# Nitric oxide-forming nitrite reductases in the anaerobic ammonium oxidizer *Kuenenia stuttgartiensis*

**DOI:** 10.1101/2025.03.25.645162

**Authors:** Femke J. Vermeir, Lotte W. Nijman, Robert S. Jansen, Laura van Niftrik, Wouter Versantvoort

**Author notes:** Authors contributed equally to the study (Femke J. Vermeir, Lotte W. Nijman). Chair of Bioinorganic Chemistry, Heinrich-Heine-Universität Düsseldorf, Universitätsstrasse 1, 40225 Düsseldorf, Germany (Wouter Versantvoort).

## Abstract

Anaerobic ammonium-oxidizing (anammox) bacteria contribute to the global nitrogen cycle by removing fixed nitrogen from the environment. They do so via the anaerobic oxidation of ammonium to dinitrogen gas, with nitrite as terminal electron acceptor. The first step in this so-called anammox reaction is the conversion of nitrite to nitric oxide by nitrite reductase. Next, nitric oxide is combined with ammonium to form hydrazine by hydrazine synthase, after which hydrazine is oxidized to dinitrogen gas by hydrazine dehydrogenase. In contrast to the other catabolic anammox enzymes, different anammox species encode different potential nitrite reductase enzymes. On top of that, there is a redundancy in genes encoding for nitrite reductase in single anammox species. The unusual diversity and redundancy in anammox nitrite reductases is unexplained. The genome of the model anammox species “*Candidatus* Kuenenia stuttgartiensis” encodes for three putative nitrite reductases. Here, we investigated which of these nitrite reductases is or are active in *K. stuttgartiensis*. Active nitric oxide-producing nitrite reductases were enriched from *K. stuttgartiensis* cells via fast protein liquid chromatography. Nitric oxide production by the enriched nitrite reductases was followed with membrane inlet mass spectrometry. Combining the activity assays with proteomics analysis indicated that the soluble nitrite reductases NirS and HAOr most strongly correlated with enzyme activity. This indicates that *K. stuttgartiensis* employs two distinct nitrite reductases to keep its nitric oxide pool replenished.

## 1. Introduction

Anaerobic ammonium-oxidizing (anammox) bacteria fulfill an important role in the global nitrogen cycle by releasing fixed nitrogen from oxygen-limited environments such as agricultural soils and oceans [1,2]. In addition to their significance in natural environments, anammox bacteria are applied in wastewater treatment plants for the cost-effective removal of nitrogen [3,4].

Anammox bacteria have a chemolithoautotrophic lifestyle and perform the catabolic anammox reaction in a dedicated intracellular compartment called the anammoxosome [5,6]. The anammox reaction proceeds via three coupled reactions interconnected by a cyclic electron flow and two reactive intermediates: nitric oxide and hydrazine [7]. First, nitrite reductase catalyzes the one-electron reduction of the substrate nitrite to nitric oxide [8,9]. Then, hydrazine synthase produces hydrazine via the combination of nitric oxide and the substrate ammonium, with the input of three electrons [10,11]. Finally, hydrazine dehydrogenase oxidizes hydrazine to the product dinitrogen gas, releasing four electrons [12]. Although the genes/enzymes that function as hydrazine synthase and hydrazine dehydrogenase in anammox bacteria are known, the identity of nitrite reductase remains ambiguous.

Anammox bacteria are cultivated in bioreactor systems as enrichment cultures that can reach up to 95% purity [13]. Enrichments and molecular approaches have identified various anammox genera, including “*Candidatus* Brocadia” [14], “*Candidatus* Kuenenia” [15], “*Candidatus* Anammoxoglobus” [16], “*Candidatus* Jettenia” [17], “*Candidatus* Anammoxibacter” [18], and “*Candidatus* Scalindua” [19]. New families and genera continue to be discovered [20,21,22]. In enrichment cultures, anammox bacteria grow slowly with doubling times between 1.8 and 12 days, depending on the bioreactor system used [23,24,25,26]. Due to slow growth, the absence of pure cultures, and a lack of standard cultivation techniques, there is currently no genetic modification toolbox available to facilitate gene and protein function studies in anammox bacteria. Therefore, our current understanding of anammox physiology and biochemistry is largely obtained from enzyme assays, transcriptomics, proteomics, and electron microscopy studies using *Kuenenia stuttgartiensis* as the anammox model species.

Different anammox genera encode different putative nitrite reductases in their genome. The genomes of *K. stuttgartiensis* and *Scalindua profunda* contain the *nirS* gene encoding a cytochrome *cd*_1_ type nitrite reductase. However, expression of this gene (kuste4136, Genbank CAJ74898) in *K. stuttgartiensis* is relatively low (1.000 RPKM [27]) compared to the expression of the genes encoding the other anammox reaction enzymes, *i*.*e*. hydrazine synthase (kuste2859, -2860 and -2861, Genbank CAJ73611, -73612 and -73613) and hydrazine dehydrogenase (kustc0694 and kustd1340, Genbank CAJ71439 and -72085) (12.000-28.000 RPKM and 25.000 RPKM, respectively [10,27]. In contrast, *nirS* is prominently expressed in *S. profunda* [28]. The genomes of *Jettenia caeni* and *Jettenia asiatica* do not contain the *nirS* gene but encode for the copper-containing nitrite reductase NirK instead [29,30,31]. *Brocadia* species seemingly lack genes encoding for canonical nitrite reductase [32,33,34], although a *nirK* gene was reported in the genome of *B. caroliensis* [35]. Due to these genomic variations in canonical nitrite reductase genes, it was postulated that anammox bacteria might harbor a novel and perhaps anammox-specific nitrite-reducing enzyme preserved throughout the different genera.

A candidate for this putative anammox nitrite reductase is the hydroxylamine oxidoreductase (HAO)-like protein HAOr. HAOr is encoded by kustc0458 (Genbank CAJ71203) and redox partner kustc0457 (Genbank CAJ71202) in *K. stuttgartiensis* and is one of the HAO-like proteins that is present in all anammox genera [36]. HAOr was isolated from *K. stuttgartiensis* and reported to reduce nitrite to nitric oxide with a rate of 0.52 nmol/min/mg protein [9]. This is relatively slow compared to the potential rate of nitrite reductases NirS and NirK in denitrifiers that produce up to 4.15 and 380 µmol/min/mg protein, respectively [37]. However, in *K. stuttgartiensis*, HAOr was detected in substantial amounts (4700 RPKM [27,38]) which could potentially meet the nitric oxide demands of hydrazine synthase. Despite its low activity, HAOr is a plausible candidate for the nitrite reductase shared among all anammox genera. Interestingly, a close homolog of HAOr, Kuste4574 (Genbank CAJ75336), has been identified. Kuste4574 is a component of one of the three Rieske/cytochrome b complexes (kuste4569–4574 (R/*b*-3) embedded in the anammoxosome membrane [36,39] and exhibits a high expression level (1800 RPKM). Similar to HAOr, this enzyme may produce nitric oxide from nitrite. However, experimental evidence confirming Kuste4574 activity is currently lacking [27,36].

In expression studies [27], *K. stuttgartiensis* HAOr and NirS both respond to changing nitrogen-species concentrations in the environment. When *K. stuttgartiensis* was grown in nitrite-limited conditions, the *nirS* and *haor* genes were significantly upregulated by 10-fold and 2.5-fold, respectively. Conversely, another study showed that the *nirS* and *haor* gene were strongly downregulated in an environment with ammonium and nitric oxide as substrates (instead of ammonium and nitrite), by 25-fold and 28-fold, respectively [40]. These data suggest that both nitrite reductases are of functional and physiological relevance in *K. stuttgartiensis* cells.

Based on current genomic and biochemical knowledge, anammox bacteria have various potential nitrite reductases at their disposal to produce nitric oxide for the activation of ammonium into hydrazine. For the model anammox species *K. stuttgartiensis* these are NirS, HAOr (both soluble) and potentially Kuste4574 (membrane-bound). In order to identify the physiological nitrite reductase in *K. stuttgartiensis*, we enriched active nitric oxide-producing nitrite reductases from *K. stuttgartiensis* cells via fast protein liquid chromatography. Nitric oxide production by the enriched nitrite reductases was followed with membrane inlet mass spectrometry and nitrite reductases were identified using proteomics.

## 2. Materials and methods

### 2.1 *Cultivation of* Kuenenia stuttgartiensis

*“Candidatus* Kuenenia stuttgartiensis*”* MBR1 [41] was grown as planktonic cells in a 12-liter membrane bioreactor (MBR) at 33°C with an OD_600_ of ∼1.2 (Applikon B.V. Schiedam, The Netherlands), as described by Kartal et al. [42]. In brief, the bioreactor was operated at 33°C and kept anoxic by continuous flushing of the bioreactor and medium vessel with argon/carbon dioxide (95/5%, 10 mL/min). The carbon dioxide in the supplied gas was sufficient to maintain the pH in the bioreactor between 7.0 and 7.4. Planktonic biomass was removed via a direct connection to a neighboring vessel at 1.1 L per day, resulting in a doubling time in the bioreactor culture of 10 days. The collected biomass was kept at room temperature under continuous sparging with dinitrogen gas to keep it anaerobic for preparation of cell-free extract.

### 2.2 Preparation of cell-free extract, soluble and membrane protein fractions

For preparation of cell-free extract and soluble and membrane protein fractions, all steps were carried out at 4°C. To collect cells, 1200 mL anaerobically collected *K. stuttgartiensis* biomass of the bioreactor described above was centrifuged at 10.000 x *g* for 10 min (Sorvall centrifuge, fixed angle). The cell pellet was resuspended in 13 mL 20 mM MOPS, 150 mM NaCl, pH 7.5 and transferred to a French press cell. Bacterial cells were broken by passing them once through the French press at 138 MPa (American Instrument Company). Whole cells and cell debris were removed by centrifugation at 5.000 x *g* for 20 min (Allegra X-15R, swinging bucket rotor, Beckman Coulter) thus obtaining the cell-free extract.

Of the cell-free extract, 1 mL was set apart for nitrite reductase activity measurements described below. To separate membranes and soluble proteins, 6 ml of the cell-free extract was subjected to ultracentrifugation at 153.300 x *g* for one hour (Optima-XE-90, Fixed-Angle 90 Ti rotor, Beckman Coulter). After centrifugation, the supernatant contained the soluble protein fraction and the pellet contained membranes and membrane proteins. The pellet was homogenized in ∼6 mL 20 mM MOPS, 150 mM NaCl, pH 7.5 with a glass potter device, thus obtaining the membrane protein fraction.

All samples were concentrated using 10 kDa molecular weight cutoff spin filters centrifuged at 4.000 x *g* (Sartorius) and their protein concentrations were measured with the Bicinchoninic acid (BCA) assay (Pierce BCA Protein Assay Kit, Thermo Scientific). Sodium dodecyl sulfate (SDS, Sigma Aldrich, 2% final concentration) was added to all samples and to the calibration curve samples (with bovine serum albumin, BSA, as standard), prior to protein quantification. SDS dissolved membranes present in the cell-free extract and membrane protein fraction, preventing their interference in the BCA assay [43].

As a control, it was evaluated whether membranes still present in cell-free extract and membrane protein fraction influenced nitrite reductase activity measurements. To this end, membranes were dissolved by incubating 1 mL of cell-free extract, membrane protein fraction, and soluble protein fraction with n-dodecyl β-D-maltoside (Thermo Scientific, 1% final concentration) in a rotating mixer overnight. The following day, insolubilized membranes and insolubilized membrane proteins were pelleted via ultracentrifugation at 142.600 x *g* for 50 min (Optima-XE-90, Fixed-Angle 90 Ti Rotor, Beckman Coulter). The remaining supernatant of cell-free extract contained soluble proteins, solubilized membrane proteins and membrane associated proteins. The supernatant of soluble protein fraction contained only soluble proteins. The supernatant of membrane protein fraction contained solubilized membrane proteins and membrane associated proteins. Samples were concentrated using

10 kDa molecular weight cutoff spin filters centrifuged at 4.000 x *g* (Sartorius) and protein concentrations were measured with the BCA assay.

### 2.3 Enrichment of soluble nitrite reductases

Note to reader: it is highly recommended to read the results section guided by Figure 8 to keep a clear overview of the experiments and fractions.

### 2.3.1 Initial separation of soluble proteins via low-resolution anion-exchange column chromatography with Q-Sepharose

Soluble protein fraction was prepared as described above. Soluble proteins (10 mL) were separated with fast protein liquid chromatography on an Äkta Purifier (GE Healthcare) with a 70 mL column packed with Q-Sepharose (XK 26/20, GE Healthcare/Cytiva). The Q-Sepharose column was equilibrated with 1.5 column volumes of 20 mM Tris-HCl buffer, pH 8.0 at a flow rate of 5 mL/min. Non-binding proteins were collected in the flow-through fraction. When the UV signal and conductivity were stable, proteins were eluted in seven column volumes via a linear gradient from 0 to 1 M NaCl and collected in 10 mL fractions. Elution of proteins was monitored at 280 nm. Collected fractions were buffer-exchanged to 20 mM MOPS, 150 mM NaCl buffer, pH 7.5 and concentrated using 10 kDa molecular weight cutoff spin filters centrifuged at 4.000 x *g* (Vivaspin 20, Sartorius Stedium Lab Ltd) for nitrite reductase activity measurements. Selected fractions that eluted from the Q-Sepharose column were pooled and further separated based on specific activity and their contribution to the total nitric oxide production.

#### 2.3.2 High-resolution anion-exchange column chromatography with Source Q15

Pooled protein fractions 9-17 (sample B) and 11-14 (sample B1) from the low-resolution anion-exchange column were further separated on a 1.7 mL Source 15Q column (4.6/100 PE, Cytiva) equilibrated with 20 column volumes of 20 mM Tris-HCl buffer, pH 8.0 with a flow rate of 2 mL/min. After the UV signal and conductivity were stable, proteins were eluted in 40 column volumes via a linear gradient from 0 to 1 M NaCl. Collected fractions of 1 mL were either pooled based on the UV signal and buffer-exchanged to 20 mM MOPS, 150 mM NaCl buffer, pH 7.5 using 10 kDa spinfilters or directly buffer-exchanged. Protein fractions were concentrated using 10 kDa spin filters centrifuged at 4.000 x *g* to measure nitrite reductase activity.

#### 2.3.3 Mixed-mode column chromatography with Hydroxyapatite

Collected flow-through of the low-resolution anion-exchange column was concentrated and buffer-exchanged to 20 mM potassium phosphate buffer, pH 7.0 using 10 kDa spinfilters. Concentrated flow-through was loaded onto a 5 mL ceramic hydroxyapatite column (EconoFit CHT Type II, 40 μm Column, Bio-Rad) equilibrated with five column volumes 20 mM potassium phosphate buffer, pH 7.0 with a flow rate of 1.5 mL/min. Proteins were eluted in nine column volumes via a linear gradient of 20 to 500 mM potassium phosphate and collected in fractions based on UV signal. The fractions were buffer-exchanged to 20 mM MOPS, 150 mM NaCl buffer, pH 7.5 and concentrated using 10 kDa molecular weight cutoff spin filters centrifuged at 4.000 x *g* to measure nitrite reductase activity.

### 2.4 Determination of protein concentration and sample purity

Protein concentrations were determined in the fractions with the absorbance of each protein sample at wavelengths 260 nm and 280 nm on a Cary 60 spectrophotometer (Agilent Technologies) with 20 mM MOPS, 150 mM NaCl buffer, pH 7.5 as baseline. From these values, protein concentrations were calculated with the formula: concentration (mg/mL) = A_280_ × 1.55 – A_260_ × 0.76 [44].

Purity of the protein samples was assessed with sodium dodecyl sulfate polyacrylamide gel electrophoresis (SDS-PAGE) [45] with a 10% resolving and 4% stacking gel. The gel was imaged on a ChemiDoc MP Imaging System (Bio-Rad).

### 2.5 Membrane-Inlet Mass Spectrometry activity assays

Nitrite reductase activity in the protein fractions was measured via ^15^N-labeled nitric oxide production followed with membrane inlet mass spectrometry (MIMS; HPR40, Positive Ion Counting detector, Hiden Analytical). The electron emission current was set to 400 μA and the measurement frequency to 1/s. Assays were performed in a custom-built 5.16 mL MIMS chamber [46] filled with 20 mM MOPS, 150 mM NaCl buffer, pH 7.5. The MIMS chamber was made anaerobic prior to measurements by flushing it with Argon and the temperature was maintained at 30°C throughout the measurements. All components of the assay were made anaerobic before they were added with a gastight syringe (Hamilton) to the MIMS chamber. Assays consisted of 200 μM phenazine ethosulfate (MP Biomedicals, LLC) and 200 μM L-ascorbic acid (Sigma) for the donation and shuttling of electrons, and 6-10 μg protein. The reaction was started by the addition of 77 μM ^15^N-labeled nitrite (99%, Cambridge Isotope Laboratories). For calibration curves, a nitric oxide stock solution was made by flushing anaerobic ultrapure water (PURELAB Chorus 1, Veolia) with 4% nitric oxide for 15 min. The nitric oxide stock concentration was calculated with the formula: concentration (M) = H^cp^ × p. In which H^cp^ for nitric oxide is 0.0019 M/atmosphere [47] and p is the partial pressure in the stock solution of nitric oxide. For each calibration level, nitric oxide was added to the MIMS chamber. Rates of nitric oxide production were determined by a linear regression fit to the initial linear part of the graph with OriginPro 2020b (OriginLab).

### 2.6 Proteomics

To identify nitrite reductases present in active protein fractions, proteomics analysis was performed. Incubations were performed at room temperature unless stated otherwise. For proteins collected from the low-resolution anion-exchange column and the high-resolution anion-exchange column, samples were prepared as follows: 8 M urea was added to the sample (1:4 (v/v)), then 10 mM dithiothreitol was added (1:500 (v/v)) and the sample was incubated for 20 min at room temperature. Subsequently, 50 mM 2-chloroacetamide was added (1:500 (v/v)) and the sample was incubated for 20 min in the dark at room temperature. Finally, proteins were digested overnight at 37°C with trypsin (1:50 (w/w). Samples were stored at -20°C until analysis.

Protein fractions collected from the mixed-mode column were diluted 1:1 (v/v) with 8 M Urea in 10 mM Tris, pH 8.0. Samples were incubated for 30 min with 1 μL 10 mM dithiothreitol per 50 μg protein in the sample. The sample volume was doubled by adding 50 mM 2-chloroacetamide in 50 mM ammonium bicarbonate (ABC) (1:1 (v/v)) and samples were incubated for 20 min in the dark. Subsequently, samples were digested overnight at 37°C with trypsin (1/50 (w/w)) and diluted 1:1 (v/v) with 2% TFA the following day. Protein fractions of 2.5 mL were desalted with OMIX C18 100 μL tips (Agilent Technologies) that were prepared by washing them two times with 100 µL 0.1% formic acid in acetonitrile, followed by two times washing with 100 µL 0.1% formic acid in ultrapure water. Protein samples were aspirated and expelled five times in order to allow proteins to bind to the OMIX C18 tip. Tips were again washed two times with 100 µL 0.1% formic acid in ultrapure water, after which proteins were eluted with 100 µL 0.1% formic acid in acetonitrile. The eluted protein samples were dried and concentrated in a vacuum centrifuge (Savant ISS110, Thermo Scientific), and resuspended in 20 μL 0.1% formic acid in ultrapure water. Samples were stored at -20°C until analysis.

Analysis of the protein fractions was performed at the Radboudumc Proteomics Center. Protein samples were analyzed by nanoflow liquid chromatography (Evosep One, Evosep Biosystems) coupled to a trapped ion mobility spectrometry – quadrupole time-of-flight mass spectrometer (timsTOF Pro2, Bruker Daltonics) via a nanoflow electrospray ionization source (CaptiveSprayer, Bruker Daltonics). Tryptic peptides (for all low- and high-resolution anion-exchange column fractions: 200 ng and for the mixed-mode column fractions 1 and 4: 200 ng, fraction 2: 100 ng and fraction 3: 10 ng) were separated by C18 reversed phase liquid chromatography (Evosep EV1106 30SPD endurance column; 150 mm length x 0.150 mm internal diameter, 1.9 μm C18AQ particles) using the pre-programmed 30 samples per day (30SPD) Evosep One method. The mass spectrometer was operated in positive ionization mode using the default data dependent acquisition – Parallel Accumulation Serial Fragmentation (dda-PASEF) [48] instrument method: 0.6 – 1.6 1/K0 mobility range, 100 – 1700 *m/z* mass range, 100 ms accumulation time, 100 ms ramp time, 10 PASEF ramps per duty cycle, 20K target PASEF intensity, 20 eV to 59 eV linear scaled collision energy between 0.6 1/K0 and 1.6 1/K0, dynamic exclusion enabled for 0.4 min.

Acquired spectra from samples of the low- and high-resolution anion-exchange column were streamed directly to ProteoScape (v2025b, Bruker Daltonics) for protein identification and label-free quantitation against the *K. stuttgartiensis* protein sequence database (Uniprot entry KSMBR1) using the following settings: Spectronaut v19 directDIA+ (Fast) workflow, 0.2 precursor PEP cutoff, 0.01 precursor Q-value cutoff, 0.01 protein Q-value cutoff global, 0.01 protein Q-value cutoff, 0.75 protein PEP cutoff, full tryptic specificity, allowed up to 2 missed cleavages, carbamidomethyl (C) as fixed modification and Oxidation (M) as variable modifications, protein group specific peptides were used for quantitation.

Acquired spectra from samples of the mixed-mode column were streamed directly to the Parallel Search Engine in Real-time (PaSER v2023, Bruker Daltonics) box for real-time database searching of acquired MS/MS spectra against the provided custom *K. stuttgartiensis* protein sequence database using the following settings: ProLuCID database search algorithm [49], 20 ppm precursor ion mass tolerance, 30 ppm fragment ion mass tolerance, full tryptic enzyme specificity, allowed up to two missed cleavages, carbamidomethyl (C) as fixed modification, deamidation (NQ) and Oxidation (M) as variable modifications, TIMScore enabled, 1% protein-level false discovery rate validation using DTA Select. Label Free Quantitation (LFQ) was performed in PaSER using Census Match Between Runs (MBR) [50] with the following parameters: 15 ppm mass accuracy, 0.03 1/K0 mobility tolerance, 30 seconds retention time tolerance, 3 isotope traces, 0.7 correlation threshold, 0.2 root mean square averaging error.

### 2.7 Calculations

For label-free quantification of proteins, the relative abundance of each protein was estimated by summing the intensity values of all detected peptide ions corresponding to that protein. Relative abundance was normalized on the injected amounts of peptides. Pearson correlations between specific enzyme activities (obtained by MIMS) and relative abundance were calculated in Excel.

Specific activity of the samples was calculated with the production of ^15^N-labeled nitric oxide in nmol/min divided by the amount of protein in the sample. From this, the purity fold per sample was calculated by dividing the specific activity by the specific activity measured in cell-free extract or soluble protein fraction. The yield of nitrite reductase activity per fraction was determined by dividing the total activity in the fraction by the total activity of cell-free extract or soluble protein fraction.

### 2.8 Figures

Figures were made in Rstudio version 4.4.1 with the ggplot2, readxl, ggrepel and cowplot packages and in Adobe Illustrator 2024.

## 3. Results

### *Soluble nitrite reductase produces most nitric oxide in* K. stuttgartiensis

To determine whether active nitrite reductase is soluble or membrane-bound, we measured nitrite reductase activity in crude cell-free extract, membrane protein fraction and soluble protein fraction. Nitrite reductase activity was followed continuously via ^15^N-labeled nitric oxide production from ^15^N-labeled nitrite using MIMS. Because the physiological electron donor for anammox nitrite reductases is unknown we chose ascorbate and phenazine ethosulfate as electron donor and carrier, respectively. These electron carriers have the ability to donate electrons to various proteins in *in vitro* enzyme assays. Moreover, their moderate midpoint potentials (E’_0_ = +80 mV for ascorbate and +55 mV for phenazine ethosulfate) make them suitable for direct nitric oxide measurement via MIMS [9].

Of the total nitrite reductase activity measured in *K. stuttgartiensis* cell-free extract, the soluble protein fraction accounted for 99 ± 17% of the activity, whereas 7 ± 4% of the activity could be attributed to the membrane protein fraction (Figure 1A and B, see also Supplementary table 1 and Figure 8). Soluble proteins produced nitric oxide at 12 ± 2 nmol/min/mg protein and nitrite reductase was two times enriched compared to cell-free extract. Membrane proteins produced nitric oxide at a rate of 0.8 ± 0.5 nmol/min/mg protein and nitrite reductase was not enriched compared to cell-free extract (purity-fold of 0.1). Notably, nitric oxide production by membrane proteins and cell-free extract stopped after four minutes whereas the production by soluble proteins continued. Experiments on DDM-solubilized membrane proteins also showed that activity ceased after four minutes (Supplementary figure 1). Thus, the membranes themselves did not interfere with nitrite reductase activity, for example, by preventing membrane-bound nitrite reductases from interacting with electron donors and carriers. The reason for the ceasing activity remains unknown. In conclusion, the majority of *K. stuttgartiensis* nitrite reductase activity could be attributed to the soluble protein fraction, indicating that the active nitrite reductase is soluble.

**Figure 1.**
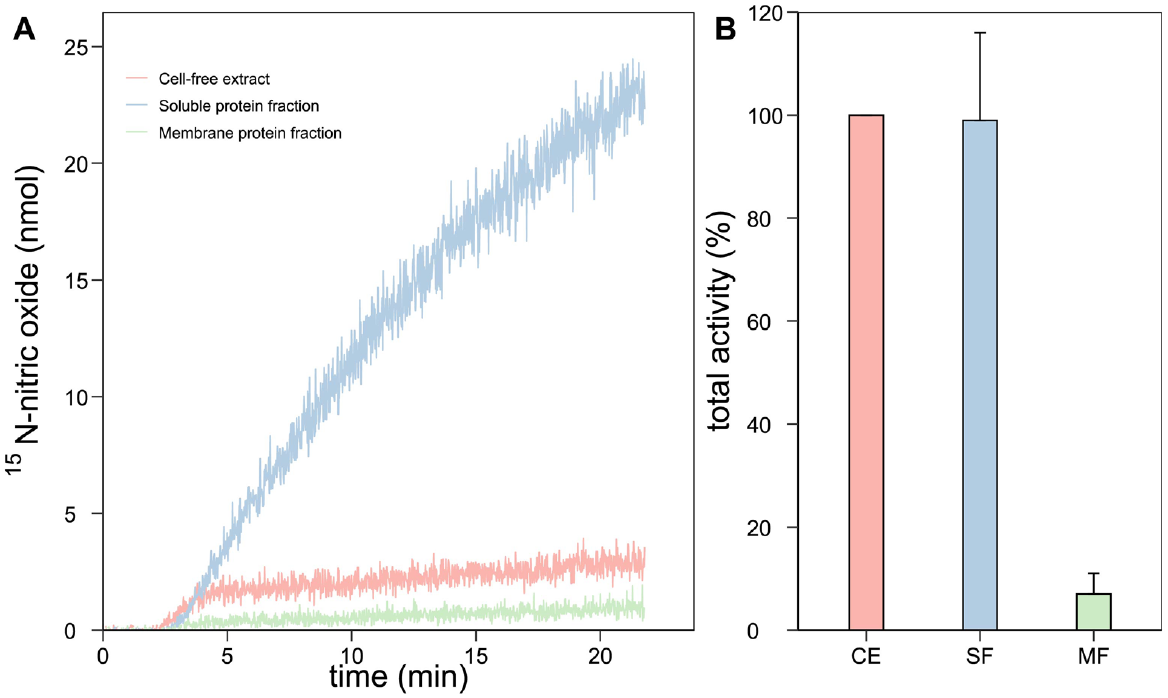
Nitrite reduction to nitric oxide in *K. stuttgartiensis* cell-free extract and protein fractions. (A) _15_N-labeled nitric oxide production from _15_N-labeled nitrite in protein fractions was followed with membrane inlet mass spectrometry. (B) Soluble proteins account for the majority of nitric oxide production from nitrite in *K. stuttgartiensis*. Activity assays contained 200 μM ascorbate and phenazine ethosulfate, and 6-10 μg protein in 20 mM MOPS, 150 mM NaCl buffer, pH 7.5. The reaction was started with 77 μM _15_N-nitrite and carried out at 30°C. CE = cell-free extract, SF = soluble protein fraction and MF = membrane protein fraction. Data are presented as mean ± SD. For A; *n*=1 biological replicate, and for B; *n*=3 biological replicates.

To identify the active nitrite reductases among the soluble proteins, we enriched nitrite reductases via FPLC. Active nitrite reductases were identified in the different protein fractions via enzyme activity assays, in which reduction of ^15^N-labeled nitrite to ^15^N-nitric oxide was followed. Subsequently, combining the measured specific activities with proteomics analysis allowed for the identification of the active nitrite reductase per protein fraction.

### 3.1 Initial separation of soluble proteins via low-resolution FPLC

For their initial separation, soluble proteins were fractionated with a low-resolution anion-exchange column (Figure 2). Nitrite reductase activity in the column fractions was measured with MIMS and showed that protein fractions 9-17 formed a peak in total nitrite reductase activity with the highest specific rate of 23 nmol nitric oxide/min/mg protein in fraction 14 (Figure 2 – please note that the black dots display the specific activity in Figure 2B). Together, the 9 fractions accounted for 65% of nitrite reductase activity measured in the soluble proteins and had a specific activity of 12 nmol nitric oxide/min/mg protein (for an overview see Supplementary table 1 and Figure 8). For clarity, the pooled fractions 9-17 were renamed to sample B. The most active fractions 11-14 within sample B were renamed to sample B1 and have a specific activity rate of 18 nmol nitric oxide/min/mg protein and account for 49% of the total nitrite reductase activity measured in all soluble proteins. The second fraction that contributed most to the total nitrite reductase activity of soluble proteins was pooled fraction 1-5. Here, nitric oxide was produced at 95 nmol/min/mg protein and the sample accounted for 11% of the total nitrite reductase activity in soluble proteins. The pooled fractions 1-5 were renamed ‘sample A’. The flow-through (FT), containing proteins that did not bind to the column, was the third fraction in terms of relative total activity. Here, nitrite reductase produced 14 nmol nitric oxide/min/mg protein which accounted for 5.3% of the total nitrite reductase activity measured in all soluble proteins. To identify the active nitrite reductase(s) in *K. stuttgartiensis*, the samples FT, A, B, and B1 were further investigated.

**Figure 2.**
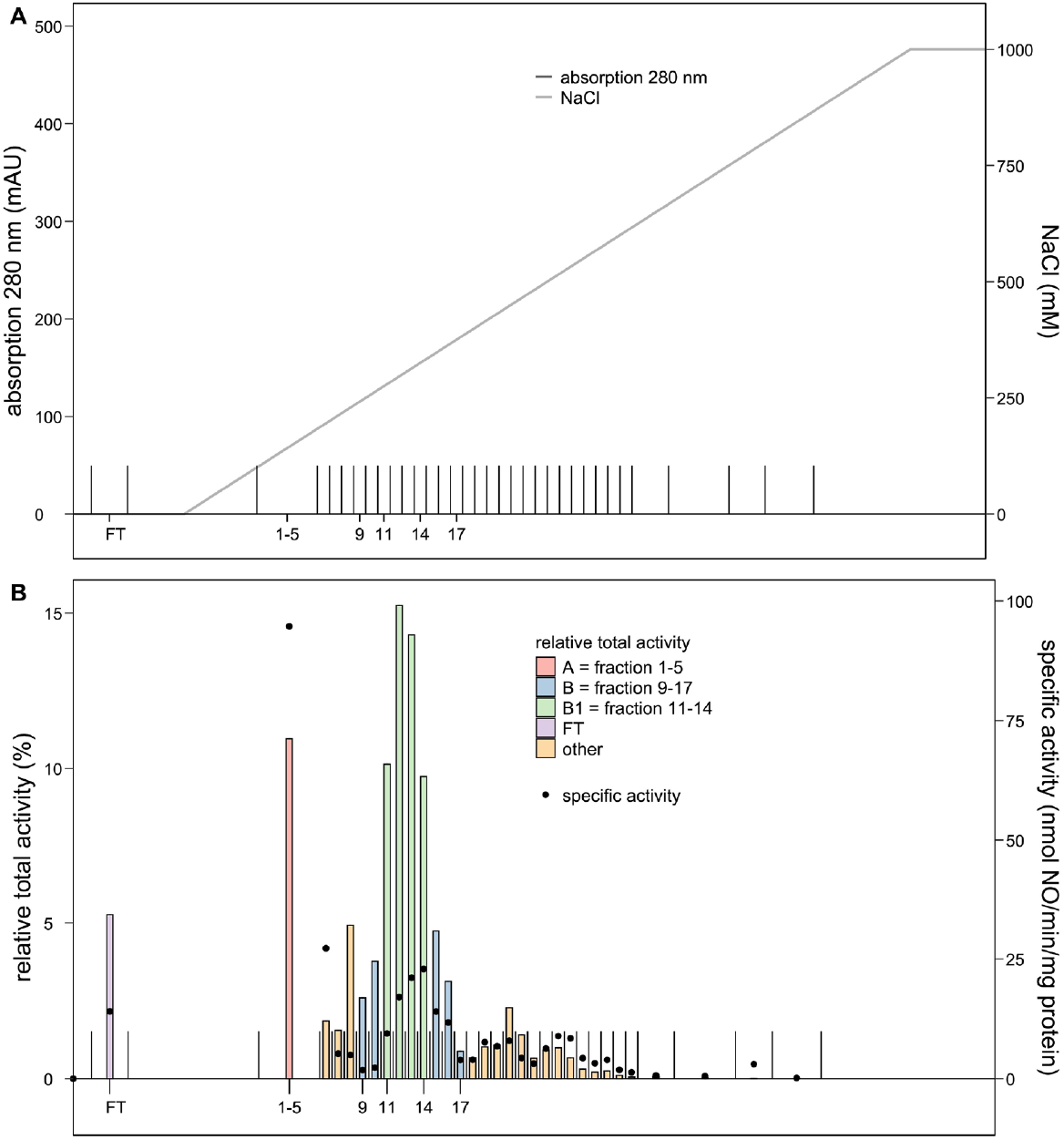
Initial separation of *K. stuttgartiensis* soluble proteins with nitrite reductase activity. (A) For the initial separation of soluble proteins, the proteins were separated on a low-resolution anion-exchange column with a linear gradient from 0 to 1 M NaCl. Eluted proteins were collected in 10 mL fractions. When the 280 nm UV signal was ≤ 25 mAU and thus indicated a low protein concentration, the fractions were pooled. (B) Of the total nitrite reductase activity of soluble proteins, 65% was measured in fractions 9-17, 50% was measured in fractions 11-14, 11% was measured in pooled fraction 1-5, and 5.3% was measured in the flow-through (FT, non-binding proteins). The majority of nitrite reductase activity was present in a single chromatographic peak of separated *K. stuttgartiensis* soluble proteins. Activity assays contained 200 μM ascorbate and phenazine ethosulfate, and 6-10 μg protein in 20 mM MOPS, 150 mM NaCl buffer, pH 7.5. The reaction was started with 77 μM _15_N-nitrite and carried out at 30°C. The relative total activity compared to activity of the total soluble proteins is expressed in percentage. The specific activity is indicated by the black dots. (*n*=1)

### 3.2 Enrichment and identification of active, soluble nitrite reductase

#### 3.2.1 High-resolution separation of sample B enriched nitrite reductase while preserving activity

The peak of nitrite reductase activity appeared in the second half of the UV absorption peak of sample B, suggesting that nitrite reductase is present in this subsection of the major UV peak (Figure 2). To further enrich nitrite reductase in sample B, we used high-resolution anion-exchange column chromatography for enhanced specificity and resolution. First, soluble proteins were again separated on a low-resolution anion-exchange column to obtain sample B. The sample accounted for 85% of the total nitrite reductase activity in soluble proteins (for an overview see Supplementary table 1 and Figure 8). Next, the proteins of sample B were loaded on a high-resolution anion-exchange column and eluted via a linear gradient from 0 to 1 M NaCl. This resulted in four UV peaks in which nitrite reductase activity was measured (Supplementary figure 2). In UV peak 4, nitrite reductase was most active and produced 36 nmol nitric oxide/min/mg protein. Furthermore, the enzyme was 5.7 times enriched compared to the soluble proteins. The proteins in UV peak 4 accounted for 60% of the total nitrite reductase activity amongst the soluble proteins (Supplementary figure 2 and for an overview see Supplementary table 1 and Figure 8). In conclusion, the two-step approach combining low- and high-resolution anion-exchange chromatography enriched nitrite reductase while largely preserving its activity.

#### 3.2.2 High-resolution separation of sample B1 enriched NirS

Next, we optimized the two-step enrichment procedure to further isolate the active nitrite reductase in sample B. Almost all nitrite reductase activity in sample B originated from fractions 11-14 (Figure 2). Thus, soluble proteins were separated on a low-resolution anion-exchange column a third time, and only fractions 11-14 were pooled to obtain sample B1. Now the sample had a specific activity of 9.7 nmol nitric oxide/min/mg protein and accounted for 22% of the total nitrite reductase activity in soluble proteins, instead of the expected rate of 18 nmol nitric oxide/min/mg protein and production of 49% that was measured in the first low-resolution separation (for an overview see Supplementary table 1 and Figure 8). Possibly, variation in elution patterns resulted in differences in total activities measured.

Alternatively, nitrite reductase may lack essential accessory components, such as cofactors or stabilizing proteins, following enrichment. However, such proteins could not be detected (Supplementary table 2). Nevertheless, separation of soluble proteins seems to lead to a loss in enzyme activity.

To further enrich active nitrite reductase in sample B1, the proteins in this sample were separated on the high-resolution anion-exchange column (Figure 3). This time, the eluted proteins were collected in 1 mL fractions independent of the UV signal. Of these collected fractions, nitric oxide was produced at 336 nmol/min/mg protein in fraction 9, and nitrite reductase was 29 times enriched compared to the soluble proteins. This most active fraction accounted for 0.8% of the total nitrite reductase activity measured in the soluble protein fraction (Figure 3, and Supplementary table 1 and Figure 8). Notably, the relative total activity of the high-resolution fractions 1-10 only attributed for 3.5% to the total nitrite reductase activity of soluble proteins. It cannot be ruled out that proteins that eluted prior to fractions 1-10 also produced nitric oxide and that the percentage of total nitrite reductase activity was higher than 3.5%.

**Figure 3.**
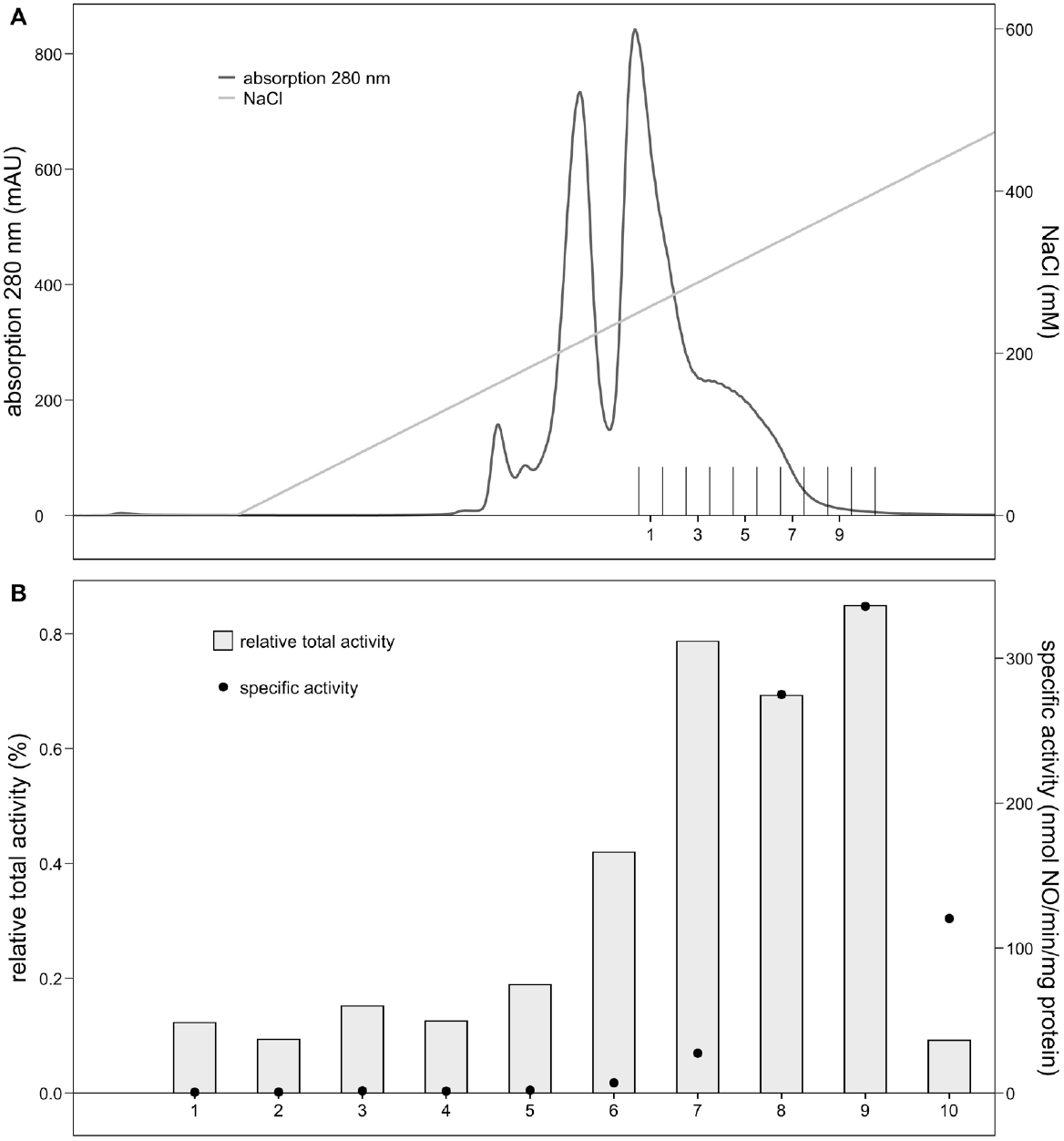
– Separation of the *K. stuttgartiensis* proteins in sample B1 by high-resolution anion-exchange column chromatography and their nitrite reductase activity. (A) Soluble proteins were first separated on a low-resolution anion-exchange column of which sample B1 showed high specific and relative total activity compared to activity measured in all soluble proteins. To enrich the active nitrite reductase in sample B1, proteins were further separated on a high-resolution anion-exchange column. Proteins were eluted with a linear gradient from 0 to 1 M NaCl in 1 mL fractions. (B) Fraction 9 of the high-resolution anion-exchange column was the most enriched and most active fraction after two-step column chromatography separation. Here, nitrite reductase produced 336 nmol nitric oxide/min/mg protein which accounted for 0.8% of the total nitrite reductase activity measured for all soluble proteins. Activity assays contained 200 μM ascorbate and phenazine ethosulfate, and 6-10 μg protein in 20 mM MOPS,150 mM NaCl buffer, pH 7.5. The reaction was started with 77 μM _15_N-nitrite and carried out at 30°C. The relative total activity compared to activity of the total soluble proteins is expressed in percentage. The specific activity is indicated by the black dots. (*n*=1)

To evaluate the purity of protein fraction 9, proteins were loaded onto an SDS-PAGE gel. However, the low protein concentration in this fraction prevented detection of the proteins (Figure 4A). To identify the active nitrite reductase in protein fraction 9, we subjected protein fractions 7-10 to proteomics analysis and surveyed the correlation between the relative abundance of identified nitrite reductase and specific activity per fraction. Proteomics revealed that NirS had the highest relative abundance measured in protein fraction 9 (Supplementary table 3). Moreover, Pearson correlation analysis showed the highest positive correlation between the relative abundance of all detected proteins and specific nitric oxide-producing activity for NirS (r = 0.87) (Figure 4B, Supplementary table 3). Other proteins with a relative abundance that positively correlated with the specific activities measured per fraction are not known to have nitrite reductase capabilities. These results strongly suggest that NirS is the enzyme responsible for the nitrite reductase activity in sample B.

**Figure 4.**
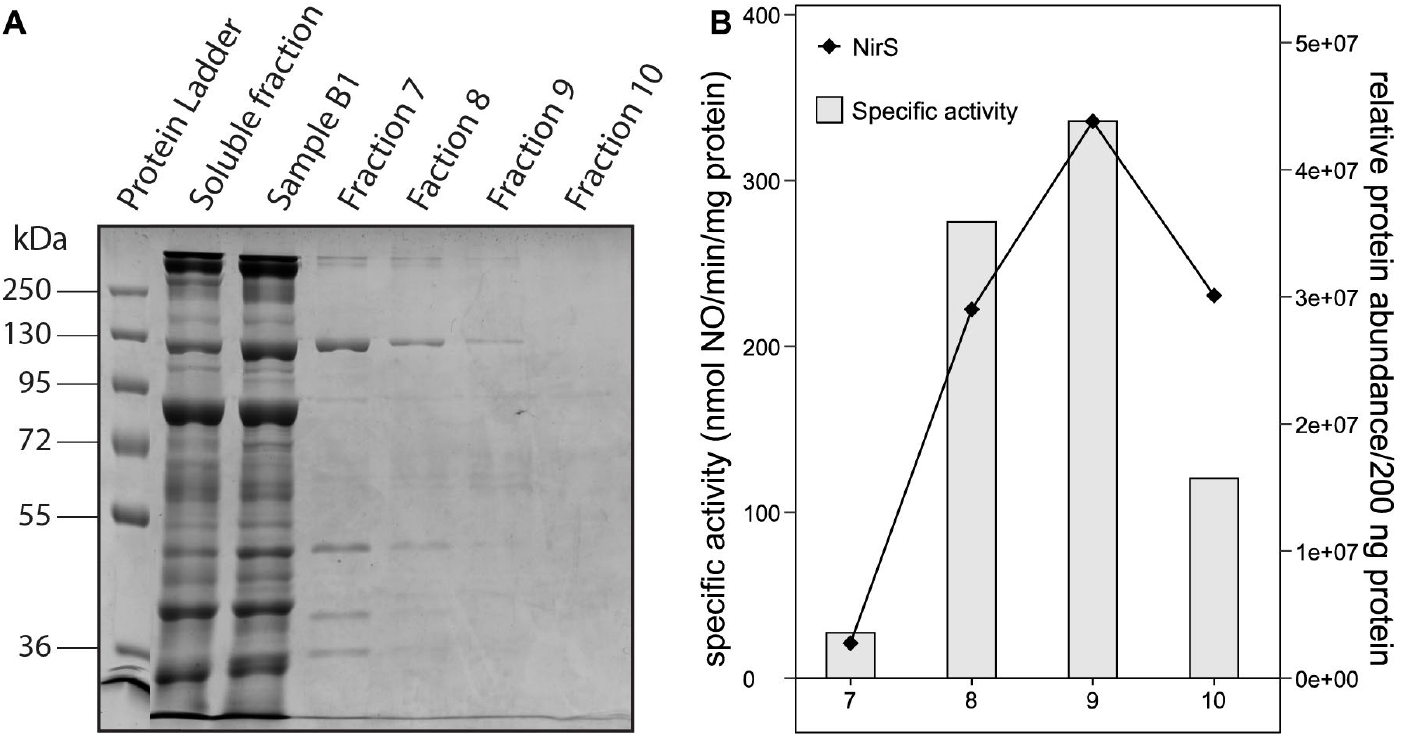
SDS-PAGE of *K. stuttgartiensis* soluble proteins and sample B1 proteins separated via high-resolution anion-exchange column chromatography, and correlation between the relative abundance of NirS and the nitrite reductase activity of these proteins. (A) SDS-PAGE of the soluble fraction, sample B1, and fractions 7, 8, 9, and 10 that showed nitrite reductase activity after separation with high-resolution anion-exchange column chromatography. The protein bands in the fractions obtained with high-resolution anion-exchange chromatography are barely visible due to the low protein concentration in the fractions. Of the soluble protein fraction and sample B1, 42 µg protein was loaded on a 10% SDS-PAGE gel. Of fractions 7, 8, 9, and 10 obtained with high-resolution anion-exchange chromatography, 5, 13, 13, and 12 µg protein was loaded on a 10% SDS-PAGE gel, respectively. (B) The relative abundance of NirS correlated well with the specific nitrite reductase activity measured in the high-resolution fractions (the specific activities are also indicated as black dots in Figure 3B). (*n*=1)

#### 3.2.3 The nitrite reductase in sample A remains unknown

Low-resolution sample A accounted for 11% of the total nitrite reductase activity amongst soluble proteins (Figure 2 and for an overview see Supplementary table 1 and Figure 8). SDS-PAGE showed that the sample contained various proteins (Figure 5). Despite the high number of different proteins present in this fraction, the calculated purity-fold of nitrite reductase in sample A compared to total soluble proteins was surprisingly high (12-fold). Proteomics analysis did not reveal any known or potential nitrite reductases (Supplementary table 4), probably because the sample was too complex and had low protein concentration.

**Figure 5.**
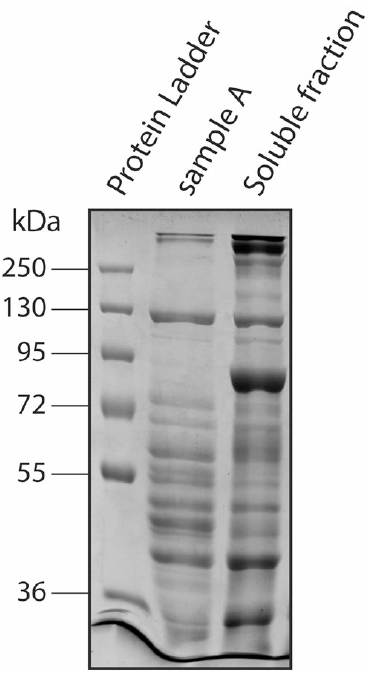
SDS-PAGE of the *K. stuttgartiensis* soluble protein fraction and sample A obtained with low-resolution anion-exchange column chromatography. The number of bands on the gel indicated that sample A contained multiple proteins. Of both samples 42 µg protein was loaded on a 10% SDS-PAGE gel.

#### 3.2.4 High-resolution separation of FT proteins enriched HAOr

The third protein fraction that contributed most to the total nitrite reductase activity, was sample FT from the low-resolution anion-exchange column (Figure 2 and for an overview see Supplementary table 1 and Figure 8). To identify the active nitrite reductase in FT, the proteins were further separated via mixed-mode chromatography on a Ceramic Hydroxyapatite column. A linear gradient of 20 to 500 mM potassium phosphate resulted in an elution pattern with four UV peaks in which nitric oxide production from nitrite was measured (Figure 6 and for an overview see Supplementary table 1 and Figure 8). Although the activities differed between biological replicates, ratios between the total activities of the UV peaks within each replica were consistent: nitrite reductase activity in UV peaks 1 and 2 were 2.5 to 3-fold lower than activity measured in UV peaks 3 and 4 (Figure 6). The highest specific activity was measured in UV peak 4, in which nitric oxide production proceeded at on average 62.5 nmol/min/mg protein. In accordance with the low number of visible bands on the SDS-PAGE gel, the enrichment of nitrite reductase was highest in this fraction: 7.5 ± 4 times compared to all soluble proteins (Figure 7A and for an overview see Supplementary table 1 and Figure 8).

**Figure 6.**
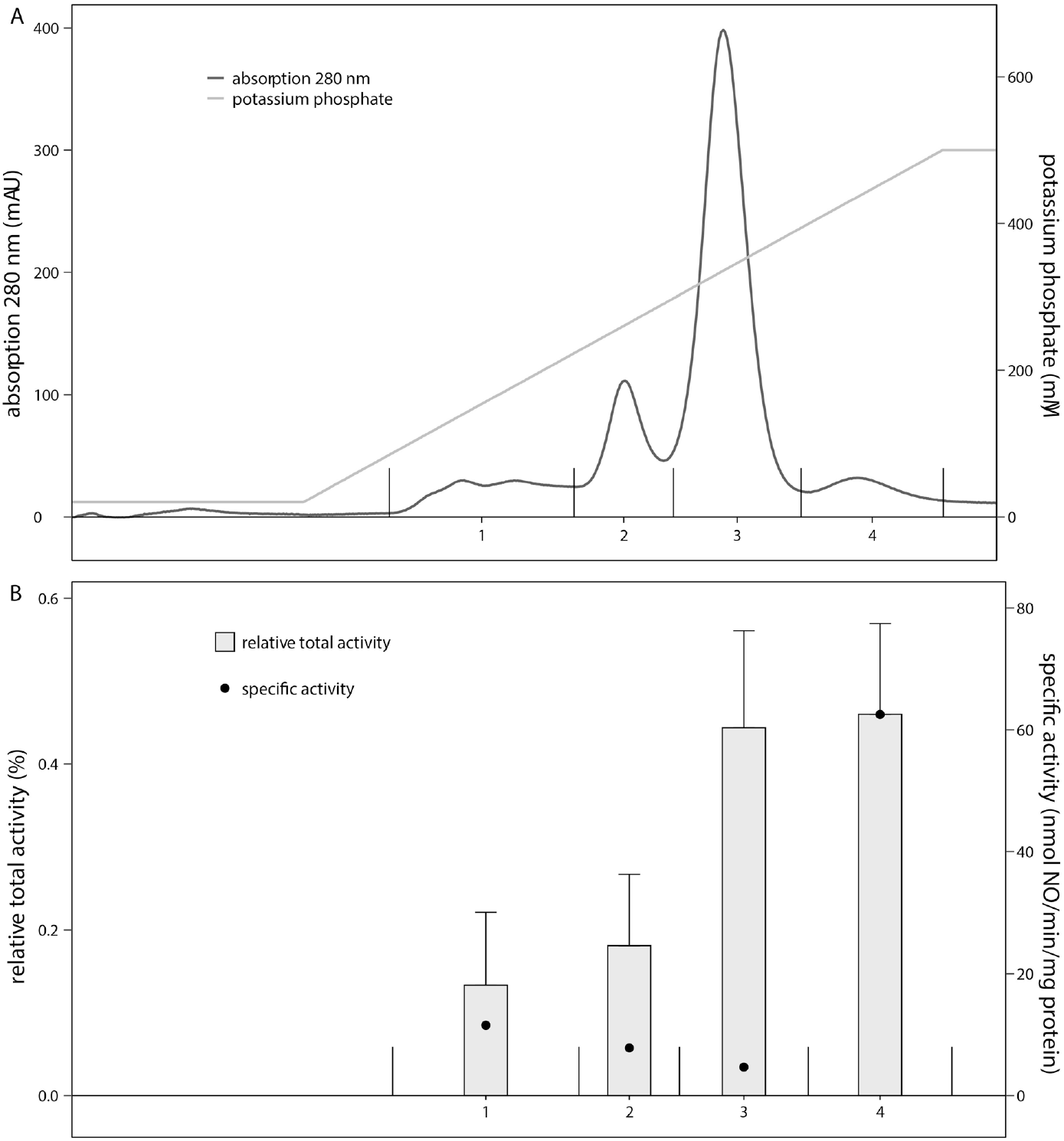
– Separation of the *K. stuttgartiensis* proteins in sample FT by mixed-mode column chromatography and their nitrite reductase activity. (A) FT proteins collected from low-resolution anion-exchange column chromatography were further fractionated via mixed-mode chromatography with a Hydroxyapatite column with a linear gradient from 20 to 500 mM potassium phosphate buffer. Proteins eluted in four distinguished UV peaks. (B) Most nitrite reductase activity in FT proteins was measured in UV peak 4 of the mixed mode chromatography column. The nitrite reductase activity accounted for 0.5% of the total nitrite reductase activity measured for all soluble proteins. Activity assays contained 200 μM ascorbate and phenazine ethosulfate, and 6-10 μg protein in 20 mM MOPS, 150 mM NaCl buffer, pH 7.5. The reaction was started with 77 μM _15_N-nitrite and carried out at 30°C. The relative total activity compared to activity of the total soluble proteins is expressed in percentage. The specific activity is indicated by the black dots. Data represented as the mean ± SD. (*n*=4 biological replicates)

**Figure 7.**
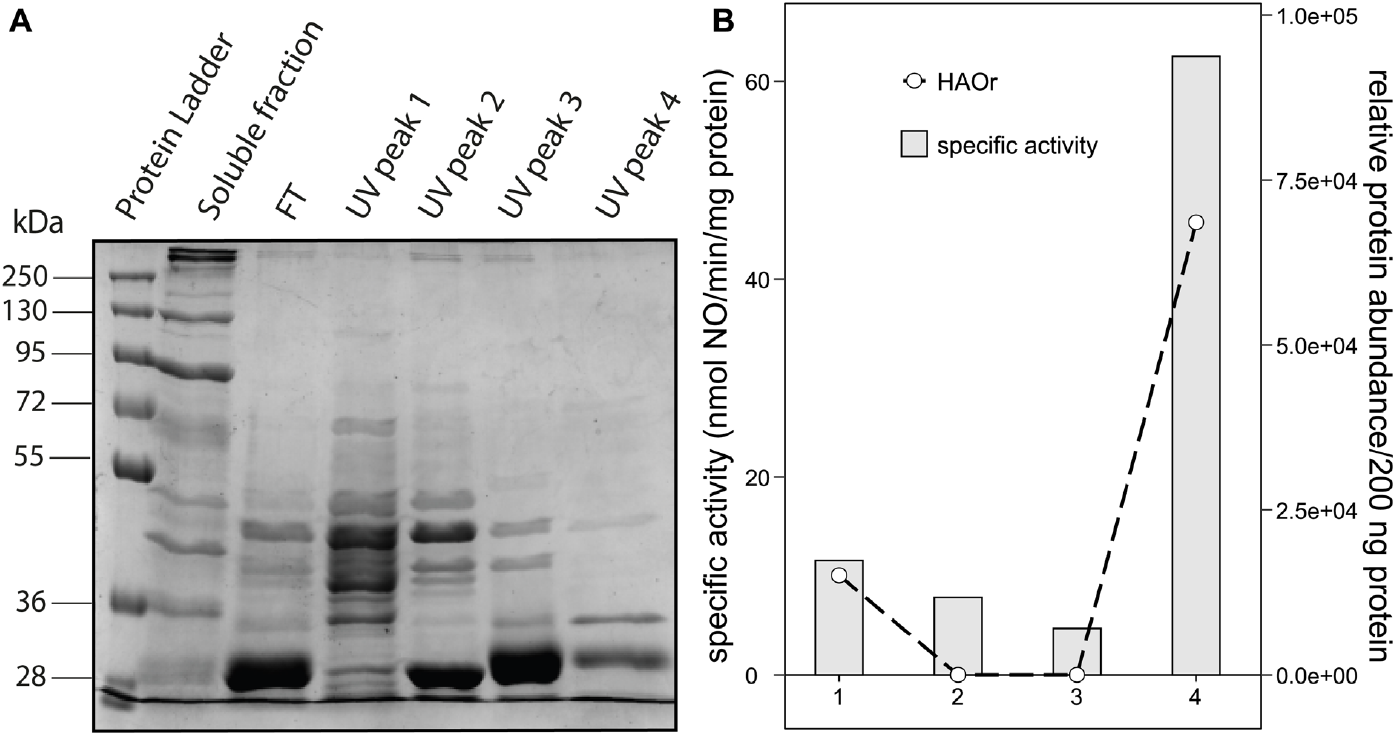
SDS-PAGE of *K. stuttgartiensis* soluble proteins and FT proteins separated via mixed-mode column chromatography, and correlation between the relative abundance of HAOr and the nitrite reductase activity of these proteins. (A) SDS-PAGE characterization of FT proteins obtained via low-resolution anion-exchange chromatography and proteins obtained via mixed-mode column chromatography. UV peak 4 showed the lowest number of visible bands. For all samples 20 µg protein was loaded on a 10% SDS gel. (B) The relative abundance of HAOr per 200 ng protein and the specific activity in nmol nitric oxide per min per mg protein measured per fraction (the specific activities are also indicated as black dots in Figure 6B). The relative abundance of HAOr correlated well with the nitrite reductase activity. (*n*=1)

**Figure 8.**
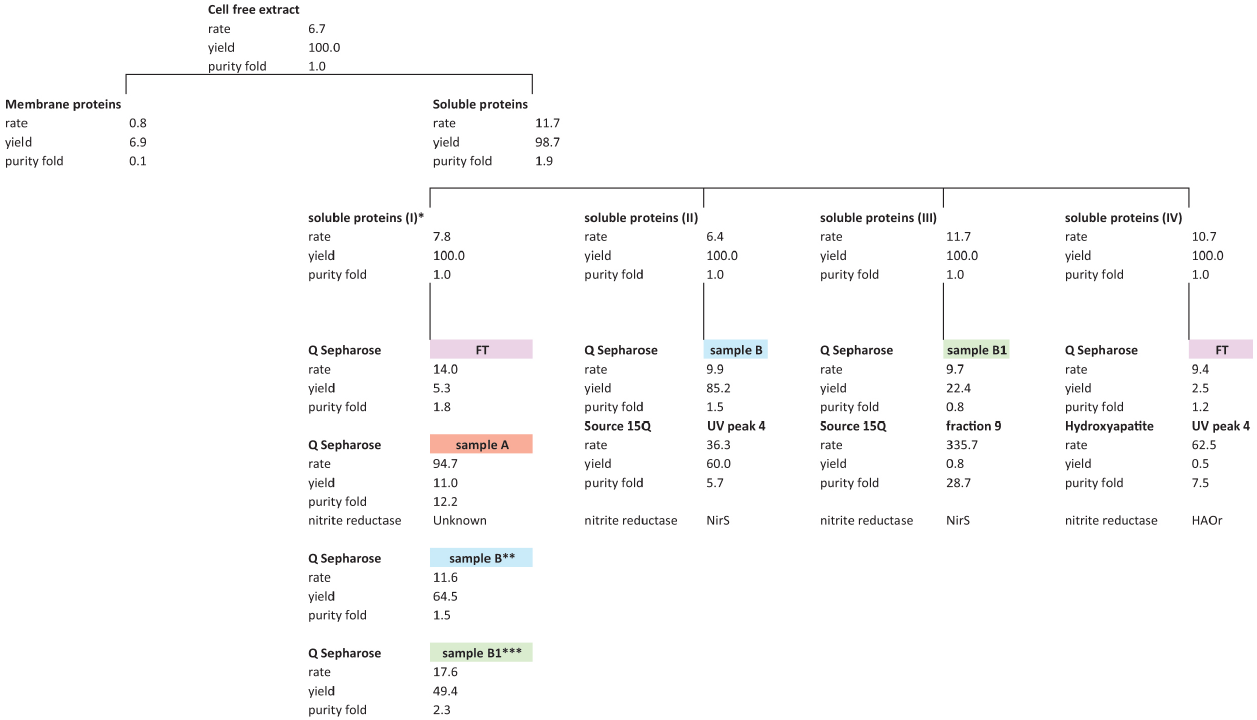
Overview of the enrichment of active nitrite reductases in *K. stuttgartiensis*. The majority of the nitrite reductase activity was measured in the soluble proteins. To identify the active nitrite reductase, soluble proteins were separated with fast protein liquid chromatography. Nitrite reductase activity per fraction was followed over time via the _15_N-nitric oxide production from _15_N-nitrite. Activity assays contained 200 μM ascorbate and phenazine ethosulfate, 6-10 μg protein in 20 mM MOPS, 150 mM NaCl buffer, pH 7.5. The reaction was started with 77 μM _15_N-nitrite and carried out at 30°C. The activity assays showed that FT, sample A, and sample B contributed most to the nitrite reductase activity. With proteomics we identified HAOr and NirS as the active nitrite reductases. The rate is expressed in nmol nitric oxide/min/mg protein, yield in % activity relative to total activity measured in the cell extract or soluble proteins, and purity fold is the specific activity per fraction compared to specific activity measured in cell extract or soluble proteins. * Roman number indicates the replicate of soluble proteins from which proteins are further separated with fast protein liquid chromatography. ** For sample B values of fractions 9-17 were combined. For the specific activity and concentration the average of fractions 9-17 was used. For the other measurements the sum of the individual fractions was used. *** For sample B1 values of fractions 11-14 were combined. For the specific activity and concentration the average of fractions 11-14 was used. For the other measurements the sum of the individual fractions was used.

Proteome analysis on fractions corresponding to UV peaks 1 to 4, identified the nitrite reductases NirS and HAOr. Pearson correlation analysis between specific nitrite reductase activity and the relative abundance of NirS and HAOr showed a strong correlation for HAOr (r=0.99) (Figure 7B) and a weak correlation for NirS (r=-0.19) (Supplementary table 5). Other proteins with a relative abundance that positively correlated with the specific activities measured per fraction are not known to function as nitrite reductase. Therefore, HAOr is presumably the active nitrite reductase in this protein fraction.

## 4. Discussion

In this study, we identified active nitrite reductases in *K. stuttgartiensis* to gain more insight in the puzzling redundancy of this enzyme in anammox bacteria. First, we showed that the active nitrite reductases are soluble proteins. In the soluble protein fraction, nitrite reductase is two times enriched compared to the cell-free extract and almost all nitrite reductase activity in the cell-free extract is recovered in the soluble protein fraction (>98%). On the contrary, activity in the membrane-bound proteins is minor and stopped shortly after the start of the assays. Using activity-guided enrichment of soluble proteins and proteomics, we identified NirS and HAOr as active nitrite reductases in two dominant activity peaks detected after low-resolution anion-exchange chromatography (65% of the total nitrite reductase activity in sample B and 5.3% in FT). Interestingly, the enriched nitrite reductases have different nitrite reduction rates. Next, we will further discuss the role of the two soluble nitrite reductases in the anammox metabolism.

### 4.1 NirS

NirS is one of the active soluble nitrite reductases in *K. stuttgartiensis* identified in this study. In this anammox species, NirS is a low abundant protein compared to the other enzymes central in the anammox catabolism *i.e*. hydrazine synthase and hydrazine dehydrogenase [27,36,39]. It could be speculated that NirS has a high specific activity rate to meet the nitric oxide demands of hydrazine synthase. Here, we measured that the most active fraction, fraction 9 of the low-resolution anion-exchange column, contained NirS and produced 336 nmol nitric oxide/min/mg protein. In previous research, *K. stuttgartiensis* batch cultures grown on nitrite and ammonium produced 30 nmol dinitrogen gas/min/mg protein [10]. Based on the measured rate in our study, NirS would need to constitute 10% of the total protein content of the cell to produce sufficient nitric oxide, which contradicts its low abundance at the proteome level. The measured rate may be underestimated due to the presence of other proteins as NirS was not purified to homogeneity in this fraction 9. Also, in a comparative study, NirS has been shown to form a highly active complex with its native electron donor, but low activity with chemical redox mediators [51]. In denitrifiers, NirS produces up to 4.15 µmol nitric oxide/min/mg protein [37], about 12 times faster than our rate measured for enriched *K. stuttgartiensis* NirS. The denitrifier rate aligns more closely with the NirS levels that could be expected in anammox cells based on transcriptome and proteome data. Further isolation of *K. stuttgartiensis* NirS could be a next step to determine whether its activity rate can be further increased compared to the value measured in this study.

### 4.2 HAOr

In addition to NirS, we identified HAOr as one of the active nitrite reductases in *K. stuttgartiensis*. The most active fraction in sample FT of the low-resolution anion-exchange column, produced 62.5 ± 30 nmol nitric oxide/min/mg protein. The activity in this sample could be attributed to HAOr. This would indicate that the catalytic activity of enriched HAOr is 125 times higher than the activity measured for the purified *K. stuttgartiensis* HAOr [9] in a nearly identical assay set-up. The difference in activity is possibly caused by the isolation procedure. While Ferousi et al. [9] eluted HAOr isocratically with 550 mM NaCl from a Q-Sepharose column, we enriched the protein from the FT of the same type of column.

In fact, we were unable to identify HAOr in the proteins bound to the low-resolution anion-exchange column in our experiments. This may be due to impurities in the fractions. Isolating HAOr from the same low-resolution anion-exchange fractions as Ferousi et al. [9], rather than the FT, would be valuable to avoid potential interference from proteins like NirS, which was identified in the FT as well, for enzyme activity determination. Further isolation of the nitrite reductase in active protein fractions bound to this column could be a next step.

### 4.3 Possible role of multiple nitrite reductases in anammox bacteria

This study demonstrates that *K. stuttgartiensis* contains two active soluble nitrite reductases, consistent with prior research showing that the *nirS* and *haor* genes are both differentially expressed under environmental nitrite limitation [27,40]. The redundancy in nitrite reductases is not a phenomenon unique to anammox bacteria. Some denitrifiers express both *nirK* and *nirS* genes, but the functional advantage of this dual expression is not fully understood [52,53,54]. For instance, it has not yet been demonstrated that NirS and NirK are both functional when present in the genome of the same denitrifier nor what mechanisms determine which nitrite reductase is expressed [52].

In *K. stuttgartiensis*, HAOr and NirS could be differentially expressed under distinct conditions aiding in the flexibility of the anammox bacteria to adapt to various environmental conditions. For instance, under low nitrite conditions it could be beneficial to express high affinity nitrite reductase to maintain growth. It would be interesting to study whether HAOr and NirS differ in nitrite affinity. NirS is a nitrite reductase with a high *K*_*m*_ between 6 and 53 µM nitrite in denitrifiers [37]. However, nitrite uptake is likely the constraining factor in nitrite-limited conditions and not the nitrite reductase rate.

In addition to improving adaptability of *K. stuttgartiensis* to environmental changes, the various anammox nitrite reductases could also be distributed to different parts of the cell. In denitrifying bacteria, nitrite reduction to nitrous oxide is coupled to energy conservation via oxidative phosphorylation [55]. Consequently, nitrite reductase is located close to the cytoplasmic membrane and mostly in the periplasm [56]. In anammox bacteria, energy is proposed to be conserved over the anammoxosome membrane [5]. Nitrite reduction to nitric oxide requires electrons that are probably donated by one of the Rieske/cytochrome b complexes of anammox bacteria [15] or the anammox nitrite oxidoreductase (NXR) [57]. Immunogold labeling studies confirmed that HAOr is located in the vicinity of the anammoxosome membrane [38]. However, the localization of NirS has not been determined yet. To understand why anammox bacteria harbor various nitrite reductases, it would be valuable to locate these enzymes in different anammox bacteria and under varying growth conditions.

## 5. Conclusion

We have shown that *K. stuttgartiensis* contains at least two nitric oxide-forming nitrite reductases, NirS and HAOr, that are active *in vitro*. This suggests that the anammox bacterium employs two nitrite reductases to keep its nitric oxide pool replenished. Containing two different nitrite reductases could improve versatility of *K. stuttgartiensis* to changes in environmental nitrite concentrations. For instance, because the enzymes have different rates and affinities. Furthermore, the different nitrite reductases could be located in different parts of the cell. While we could identify nitrite reductases that are present and active in *K. stuttgartiensis*, results of enzyme assays are difficult to translate to physiological enzyme activity. To follow up on this study, it would be interesting to evaluate nitrite reductase activity in different anammox species and within species grown in different nitrite concentrations. Moreover, the active protein fractions isolated here could be further enriched in nitrite reductase to identify and characterize the enzyme present there. Lastly, immunogold localization could enhance the understanding of nitrite reductase location and function within the cell. In conclusion, the redundancy in genes encoding potential nitrite reductases, along with the identification and activity of two distinct enzymes, underscores the critical role of nitric oxide in anammox bacteria.

## 7. Acknowledgements and funding sources

We would like to thank Guylaine Nuijten for assistance and co-maintenance of anammox bioreactor enrichment cultures, Hans Wessels from the Radboudumc Proteomics Center for help with the proteomic analyses, and Marjan Smeulders and Mike Jetten for helpful discussions.

LvN, FJV and WV were supported by VIDI grant VI.Vidi.192.001 awarded to LvN by the Dutch Research Council (NWO).

## References

[1] Lam, P. & Kuypers, M. M. 2011. Microbial nitrogen cycling processes in oxygen minimum zones. Ann Rev Mar Sci, 3, 317–45.

[2] Nie, S. A., Zhu, G.-B., Singh, B. & Zhu, Y.-G. 2019. Anaerobic ammonium oxidation in agricultural soils-synthesis and prospective. Environmental Pollution, 244, 127–134.

[3] Kartal, B., Kuenen, J. G. & Van Loosdrecht, M. C. M. 2010. Sewage treatment with anammox. Science, 328, 702–703.

[4] Lackner, S., Gilbert, E. M., Vlaeminck, S. E., Joss, A., Horn, H. & Van Loosdrecht, M. C. M. 2014. Full-scale partial nitritation/anammox experiences – An application survey. Water research, 55, 292–303.

[5] Neumann, S., Wessels, H. J., Rijpstra, W. I., Sinninghe Damste, J. S., Kartal, B., Jetten, M. S. & Van Niftrik, L. 2014. Isolation and characterization of a prokaryotic cell organelle from the anammox bacterium Kuenenia stuttgartiensis. Mol Microbiol, 94, 794–802.

[6] Van Niftrik, L., Geerts, W. J., Van Donselaar, E. G., Humbel, B. M., Yakushevska, A., Verkleij, A. J., Jetten, M. S. & Strous, M. 2008. Combined structural and chemical analysis of the anammoxosome: a membrane-bounded intracytoplasmic compartment in anammox bacteria. J Struct Biol, 161, 401–10.

[7] Kartal, B. & Keltjens, J. T. 2016. Anammox Biochemistry: a Tale of Heme c Proteins. Trends Biochem Sci, 41, 998–1011.

[8] Maalcke, W. J., Dietl, A., Marritt, S. J., Butt, J. N., Jetten, M. S., Keltjens, J. T., Barends, T. R. & Kartal, B. 2014. Structural basis of biological NO generation by octaheme oxidoreductases. J Biol Chem, 289, 1228–42.

[9] Ferousi, C., Schmitz, R. A., Maalcke, W. J., Lindhoud, S., Versantvoort, W., Jetten, M. S. M., Reimann, J. & Kartal, B. 2021. Characterization of a nitrite-reducing octaheme hydroxylamine oxidoreductase that lacks the tyrosine cross-link. J Biol Chem, 296, 100476.

[10] Kartal, B., Maalcke, W. J., De Almeida, N. M., Cirpus, I., Gloerich, J., Geerts, W., Op Den Camp, H. J. M., Harhangi, H. R., Janssen-Megens, E. M., Francoijs, K.-J., Stunnenberg, H. G., Keltjens, J. T., Jetten, M. S. M. & Strous, M. 2011b. Molecular mechanism of anaerobic ammonium oxidation. Nature, 479, 127–130.

[11] Dietl, A., Ferousi, C., Maalcke, W. J., Menzel, A., de Vries, S., Keltjens, J. T., Jetten, M. S. M., Kartal, B. & Barends, T. R. M. 2015. The inner workings of the hydrazine synthase multiprotein complex. Nature 527, 394-397.

[12] Maalcke, W. J., Reimann, J., De Vries, S., Butt, J. N., Dietl, A., Kip, N., Mersdorf, U., Barends, T. R., Jetten, M. S., Keltjens, J. T. & Kartal, B. 2016. Characterization of Anammox Hydrazine Dehydrogenase, a Key N2-producing Enzyme in the Global Nitrogen Cycle. J Biol Chem, 291, 17077–92.

[13] Peeters, S. H. & Van Niftrik, L. 2019. Trending topics and open questions in anaerobic ammonium oxidation. Curr Opin Chem Biol, 49, 45–52.

[14] Strous, M., Fuerst, J. A., Kramer, E. H., Logemann, S., Muyzer, G., Van De Pas-Schoonen, K. T., Webb, R., Kuenen, J. G. & Jetten, M. S. 1999. Missing lithotroph identified as new planctomycete. Nature, 400, 446–9.

[15] Strous, M., Pelletier, E., Mangenot, S., Rattei, T., Lehner, A., Taylor, M. W., Horn, M., Daims, H., Bartol-Mavel, D., Wincker, P., Barbe, V., Fonknechten, N., Vallenet, D., Segurens, B., Schenowitz-Truong, C., Medigue, C., Collingro, A., Snel, B., Dutilh, B. E., Op Den Camp, H. J., Van Der Drift, C., Cirpus, I., Van De Pas-Schoonen, K. T., Harhangi, H. R., Van Niftrik, L., Schmid, M., Keltjens, J., Van De Vossenberg, J., Kartal, B., Meier, H., Frishman, D., Huynen, M. A., Mewes, H. W., Weissenbach, J., Jetten, M. S., Wagner, M. & Le Paslier, D. 2006. Deciphering the evolution and metabolism of an anammox bacterium from a community genome. Nature, 440, 790–4.

[16] Kartal, B., Van Niftrik, L., Rattray, J., Van De Vossenberg, J. L., Schmid, M. C., Sinninghe Damsté, J., Jetten, M. S. & Strous, M. 2008. Candidatus ‘Brocadia fulgida’: an autofluorescent anaerobic ammonium oxidizing bacterium. FEMS microbiology ecology, 63, 46–55.

[17] Quan, Z. X., Rhee, S. K., Zuo, J. E., Yang, Y., Bae, J. W., Park, J. R., Lee, S. T. & Park, Y. H. 2008. Diversity of ammonium-oxidizing bacteria in a granular sludge anaerobic ammonium-oxidizing (anammox) reactor. Environ Microbiol, 10, 3130–9.

[18] Suarez, C., Dalcin Martins, P., Jetten, M. S. M., Karačić, S., Wilén, B. M., Modin, O., Hagelia, P., Hermansson, M. & Persson, F. 2022. Metagenomic evidence of a novel family of anammox bacteria in a subsea environment. Environ Microbiol, 24, 2348–2360.

[19] Kuypers, M. M. M., Sliekers, A. O., Lavik, G., Schmid, M., Jørgensen, B. B., Kuenen, J. G., Sinninghe Damsté, J. S., Strous, M. & Jetten, M. S. M. 2003. Anaerobic ammonium oxidation by anammox bacteria in the Black Sea. Nature, 422, 608–611.

[20] Yang, Y., Lu, Z., Azari, M., Kartal, B., Du, H., Cai, M., Herbold, C. W., Ding, X., Denecke, M. & Li, X. 2022. Discovery of a new genus of anaerobic ammonium oxidizing bacteria with a mechanism for oxygen tolerance. Water Research, 226, 119165.

[21] Zhao, R., Biddle, J. F. & Jørgensen, S. L. 2022. Introducing Candidatus Bathyanammoxibiaceae, a family of bacteria with the anammox potential present in both marine and terrestrial environments. ISME Communications, 2.

[22] Zhao, R., Le Moine Bauer, S. & Babbin, A. R. 2023. “Candidatus Subterrananammoxibiaceae,” a New Anammox Bacterial Family in Globally Distributed Marine and Terrestrial Subsurfaces. Appl Environ Microbiol, 89, e008002

[23] Isaka, K., Date, Y., Sumino, T., Yoshie, S. & Tsuneda, S. 2006. Growth characteristic of anaerobic ammonium-oxidizing bacteria in an anaerobic biological filtrated reactor. Applied Microbiology and Biotechnology, 70, 47–52.

[24] Lotti, T., Kleerebezem, R., Abelleira-Pereira, J. M., Abbas, B. & Van Loosdrecht, M. C. 2015. Faster through training: The anammox case. Water Res, 81, 261–8.

[25] Tsushima, I., Kindaichi, T. & Okabe, S. 2007. Quantification of anaerobic ammonium-oxidizing bacteria in enrichment cultures by real-time PCR. Water Research, 41, 785–794.

[26] Van Der Star, W. R. L., Abma, W. R., Blommers, D., Mulder, J.-W., Tokutomi, T., Strous, M., Picioreanu, C. & Van Loosdrecht, M. C. M. 2007. Startup of reactors for anoxic ammonium oxidation: experiences from the first full-scale anammox reactor in Rotterdam. Water research, 41, 4149–4163.

[27] Smeulders, M. J., Peeters, S. H., Van Alen, T., De Bruijckere, D., Nuijten, G. H. L., Op Den Camp, H. J. M., Jetten, M. S. M. & Van Niftrik, L. 2020. Nutrient Limitation Causes Differential Expression of Transport-and Metabolism Genes in the Compartmentalized Anammox Bacterium Kuenenia stuttgartiensis. Front Microbiol, 11, 1959.

[28] Van De Vossenberg, J., Woebken, D., Maalcke, W. J., Wessels, H. J., Dutilh, B. E., Kartal, B., Janssen-Megens, E. M., Roeselers, G., Yan, J., Speth, D., Gloerich, J., Geerts, W., Van Der Biezen, E., Pluk, W., Francoijs, K. J., Russ, L., Lam, P., Malfatti, S. A., Tringe, S. G., Haaijer, S. C., Op Den Camp, H. J., Stunnenberg, H. G., Amann, R., Kuypers, M. M. & Jetten, M. S. 2013. The metagenome of the marine anammox bacterium ‘Candidatus Scalindua profunda’ illustrates the versatility of this globally important nitrogen cycle bacterium. Environ Microbiol, 15, 1275–89.

[29] Ali, M., Oshiki, M., Awata, T., Isobe, K., Kimura, Z., Yoshikawa, H., Hira, D., Kindaichi, T., Satoh, H., Fujii, T. & Okabe, S. 2015. Physiological characterization of anaerobic ammonium oxidizing bacterium ‘Candidatus Jettenia caeni’. Environ Microbiol, 17, 2172–89.

[30] Hira, D., Toh, H., Migita, C. T., Okubo, H., Nishiyama, T., Hattori, M., Furukawa, K. & Fujii, T. 2012. Anammox organism KSU-1 expresses a NirK-type copper-containing nitrite reductase instead of a NirS-type with cytochrome cd1. FEBS Lett, 586, 1658–63.

[31] Hu, Z., Speth, D. R., Francoijs, K. J., Quan, Z. X. & Jetten, M. S. 2012. Metagenome Analysis of a Complex Community Reveals the Metabolic Blueprint of Anammox Bacterium “Candidatus Jettenia asiatica”. Front Microbiol, 3, 366.

[32] Gori, F., Tringe, S. G., Kartal, B., Marchiori, E. & Jetten, M. S. 2011. The metagenomic basis of anammox metabolism in Candidatus ‘Brocadia fulgida’. Biochem Soc Trans, 39, 1799–804.

[33] Okubo, T., Toyoda, A., Fukuhara, K., Uchiyama, I., Harigaya, Y., Kuroiwa, M., Suzuki, T., Murakami, Y., Suwa, Y. & Takami, H. 2021. The physiological potential of anammox bacteria as revealed by their core genome structure. DNA Res, 28.

[34] Oshiki, M., Shinyako-Hata, K., Satoh, H. & Okabe, S. 2015. Draft Genome Sequence of an Anaerobic Ammonium-Oxidizing Bacterium, “Candidatus Brocadia sinica”. Genome Announc, 3.

[35] Park, H., Brotto, A. C., Van Loosdrecht, M. C. M. & Chandran, K. 2017. Discovery and metagenomic analysis of an anammox bacterial enrichment related to Candidatus “Brocadia caroliniensis” in a full-scale glycerol-fed nitritation-denitritation separate centrate treatment process. Water Research, 111, 265–273.

[36] Kartal, B., De Almeida, N. M., Maalcke, W. J., Op Den Camp, H. J., Jetten, M. S. & Keltjens, J. T. 2013. How to make a living from anaerobic ammonium oxidation. FEMS Microbiol Rev, 37, 428–61.

[37] Cutruzzolà, F. 1999. Bacterial nitric oxide synthesis. Biochimica et Biophysica Acta (BBA) - Bioenergetics, 1411, 231–249.

[38] De Almeida, N. M., Neumann, S., Mesman, R. J., Ferousi, C., Keltjens, J. T., Jetten, M. S., Kartal, B. & Van Niftrik, L. 2015. Immunogold Localization of Key Metabolic Enzymes in the Anammoxosome and on the Tubule-Like Structures of Kuenenia stuttgartiensis. J Bacteriol, 197, 2432–41.

[39] De Almeida, N. M., Wessels, H. J., De Graaf, R. M., Ferousi, C., Jetten, M. S., Keltjens, J. T. & Kartal, B. 2016. Membrane-bound electron transport systems of an anammox bacterium: A complexome analysis. Biochim Biophys Acta, 1857, 1694–704.

[40] Hu, Z., Wessels, H., Van Alen, T., Jetten, M. S. M. & Kartal, B. 2019. Nitric oxide-dependent anaerobic ammonium oxidation. Nat Commun, 10, 1244.

[41] Frank, J., Lucker, S., Vossen, R., Jetten, M. S. M., Hall, R. J., Op Den Camp, H. J. M. & Anvar, S. Y. 2018. Resolving the complete genome of Kuenenia stuttgartiensis from a membrane bioreactor enrichment using Single-Molecule Real-Time sequencing. Sci Rep, 8, 4580.

[42] Kartal, B., Geerts, W. & Jetten, M. S. 2011a. Cultivation, detection, and ecophysiology of anaerobic ammonium-oxidizing bacteria. Methods Enzymol, 486, 89–108.

[43] Morton, R. E. & A., E. T. 1992. Modification of the Bicinchoninic Acid Protein Assay to Eliminate Lipid Interference in Determining Lipoprotein Protein Content. Analytical Biochemistry, 204, 332–334.

[44] Stepanchenko, N. S., Novikova, G. V. & Moshkov, I. E. 2011. Protein quantification. Russian Journal of Plant Physiology, 58, 737–742.

[45] Laemmli, U. K. 1970. Cleavage of structural proteins during the assembly of the head of bacteriophage T4. Nature, 227, 680 – 68.

[46] Schmitz, R. A., Pol, A., Mohammadi, S. S., Hogendoorn, C., van Gelder, A. H., Jetten, M. S. M., Daumann, L. & Op den Camp, H. J. M. 2020. The thermoacidophilic methanotroph Methylacidiphilum fumariolicum SolV oxidizes subatmospheric H_2_ with a high-affinity, membrane-associated [NiFe] hydrogenase. ISME J, 41, 1223–1232.

[47] Sander, R. 2015. Compilation of Henry’s law constants (version 4.0) for water as solvent. Atmospheric Chemistry and Physics, 15, 4399–4981.

[48] Meier, F., Park, M. A. & Mann, M. 2021. Trapped Ion Mobility Spectrometry and Parallel Accumulation-Serial Fragmentation in Proteomics. Mol Cell Proteomics, 20, 100138.

[49] Xu, T., Park, S. K., Venable, J. D., Wohlschlegel, J. A., Diedrich, J. K., Cociorva, D., Lu, B., Liao, L., Hewel, J., Han, X., Wong, C. C. L., Fonslow, B., Delahunty, C., Gao, Y., Shah, H. & Yates, J. R., 3rd 2015. ProLuCID: An improved SEQUEST-like algorithm with enhanced sensitivity and specificity. J Proteomics, 129, 16–24.

[50] Park, S. K. & Yates, J. R., 3rd 2010. Census for proteome quantification. Curr Protoc Bioinformatics, Chapter 13, 13.12.1-13.12.11.

[51] Pedroso, H. A., Silveira, C. M., Almeida, R. M., Almeida, A., Besson, S., Moura, I., Moura, J. J. G. & Almeida, M. G. 2016. Electron transfer and docking between cytochrome cd_1_nitrite reductase and different redox partners — A comparative study. BBA Bioenergetics, 1857, 1412–1421.

[52] Graf, D. R., Jones, C. M. & Hallin, S. 2014. Intergenomic comparisons highlight modularity of the denitrification pathway and underpin the importance of community structure for N2O emissions. PLoS One, 9, e114118.

[53] Sanchez, C. & Minamisawa, K. 2018. Redundant roles of Bradyrhizobium oligotrophicum Cu-type (NirK) and cd1-type (NirS) nitrite reductase genes under denitrifying conditions. FEMS Microbiol Lett, 365.

[54] Wittorf, L., Jones, C. M., Bonilla-Rosso, G. & Hallin, S. 2018. Expression of nirK and nirS genes in two strains of Pseudomonas stutzeri harbouring both types of NO-forming nitrite reductases. Res Microbiol, 169, 343–347.

[55] Koike, I. & Hattori, A. 1975. Energy Yield of Denitrification: An Estimate from Growth Yield in Continuous Cultures of Pseudomonas denitrificans under Nitrate-, Nitrite- and Nitrous Oxide-limited Conditions. Journal of General Microbiology 88, 11–19.

[56] Coyne, M. S., Arunakumari, A., Pankratz, H. S. & Tiedje, J. M. 1990. Localization of the Cytochrome cdl and Copper Nitrite Reductases in Denitrifying Bacteria. Journal of Bacteriology, 172, 2558–2562.

[57] Chicano, T. M., Dietrich, L., De Almeida, N. M., Akram, M., Hartmann, E., Leidreiter, F., Leopoldus, D., Mueller, M., Sánchez, R., Nuijten, G. H. L., Reimann, J., Seifert, K. A., Schlichting, I., Van Niftrik, L., Jetten, M. S. M., Dietl, A., Kartal, B., Parey, K. & Barends, T. R. M. 2021. Structural and functional characterization of the intracellular filament-forming nitrite oxidoreductase multiprotein complex. Nat Microbiol, 6, 1129–1139.

