## Supplementary Material for "Nitric oxide-forming nitrite reductases in the anaerobic ammonium oxidizer *Kuenenia stuttgartiensis*"

**Supplementary table 1** – Overview of the enrichment of active nitrite reductases in *K. stuttgartiensis*. The majority of the nitrite reductase activity was measured in the soluble protein fraction. To identify the active nitrite reductase, soluble proteins were separated with fast protein liquid chromatography. Nitrite reductase activity per fraction was followed over time via the $^{15}$N-nitric oxide production from $^{15}$N-nitrite. Activity assays contained 200 µM ascorbate and phenazine ethosulfate, 6-10 µg protein in 20 mM MOPS, 150 mM NaCl buffer, pH 7.5. The reaction was started with 77 µM $^{15}$N-labeled nitrite and carried out at 30°C. The activity assays revealed that FT, sample A, and sample B contributed most to the nitrite reductase activity. With proteomics we identified HAOr and NirS as the active nitrite reductases.

| column | sample | [protein] (mg/mL) | volume (mL) | total protein (mg) | specific activity (nmol/min/mg protein) | total activity (nmol/min) | yield (%) | purity fold | nitrite reductase |
|---|---|---|---|---|---|---|---|---|---|
| - | cell free extract | 14.1 | 13.9 | 196.44 | 6.7 | 1319.2 | 100.0 | 1.0 | |
| - | soluble proteins | 7.1 | 5.8 | 39.50 | 11.7 | 463.7 | 98.7 | 1.9 | |
| - | membrane proteins | 5.8 | 6.5 | 49.40 | 0.8 | 39.0 | 6.9 | 0.1 | |
| - | soluble proteins (I)* | 22.8 | 10.00 | 227.50 | 7.8 | 1773.3 | 100.0 | 1.0 | |
| Q Sepharose | FT | 14.8 | 0.45 | 6.65 | 14.0 | 93.4 | 5.3 | 1.8 | |
| Q Sepharose | sample A | 3.4 | 0.60 | 2.05 | 94.7 | 194.2 | 11.0 | 12.2 | Unidentified |
| Q Sepharose | sample B** | 23.1 | 6.15 | 124.13 | 11.6 | 1.4 | 64.5 | 1.5 | |
| Q Sepharose | sample B1*** | 23.4 | 2.70 | 54.5 | 17.6 | 1.0 | 49.4 | 2.3 | |
| - | soluble proteins (II) | 17.8 | 10.0 | 178.00 | 6.4 | 1140.0 | 100.0 | 1.0 | |
| Q Sepharose | sample B | 130.5 | 0.8 | 97.88 | 9.9 | 971.3 | 85.2 | 1.5 | |
| Source 15Q | UV peak 1 | 2.6 | 1.6 | 4.20 | 7.0 | 29.4 | 2.6 | 1.1 | |
| Source 15Q | UV peak 2 | 19.6 | 1.7 | 33.48 | 1.0 | 33.6 | 2.9 | 0.2 | |
| Source 15Q | UV peak 3 | 18.2 | 1.6 | 28.94 | 0.7 | 21.1 | 1.9 | 0.1 | |
| Source 15Q | UV peak 4 | 13.1 | 1.4 | 18.82 | 36.3 | 684.1 | 60.0 | 5.7 | NirS |
| - | soluble proteins (III) | 13.4 | 10.0 | 134.00 | 11.7 | 1572.1 | 100.0 | 1.0 | |
| Q Sepharose | sample B1 | 48.6 | 0.8 | 36.45 | 9.7 | 352.0 | 22.4 | 0.8 | |
| Source 15Q | Fraction 7 | 0.9 | 0.5 | 0.45 | 27.4 | 12.4 | 0.8 | 2.3 | |
| Source 15Q | Fraction 8 | 0.1 | 0.5 | 0.04 | 275.0 | 10.9 | 0.7 | 23.5 | |
| Source 15Q | Fraction 9 | 0.1 | 0.8 | 0.04 | 335.7 | 13.3 | 0.8 | 28.7 | NirS |
| Source 15Q | Fraction 10 | 0.02 | 0.6 | 0.01 | 120.5 | 1.4 | 0.1 | 10.3 | |
| - | soluble proteins (IV) | 19.4 | 22.5 | 435.87 | 10.7 | 4673.2 | 100.0 | 1.0 | |
| Q Sepharose | FT | 2.1 | 5.0 | 12.59 | 9.4 | 118.7 | 2.5 | 1.2 | |
| Hydroxyapatite | UV peak 1 | 0.8 | 0.6 | 0.55 | 11.6 | 6.4 | 0.1 | 1.0 | |
| Hydroxyapatite | UV peak 2 | 1.6 | 0.6 | 1.17 | 7.8 | 9.1 | 0.2 | 0.9 | |
| Hydroxyapatite | UV peak 3 | 7.3 | 0.6 | 4.81 | 4.7 | 22.5 | 0.5 | 0.5 | |
| Hydroxyapatite | UV peak 4 | 0.5 | 0.6 | 0.35 | 62.5 | 21.9 | 0.5 | 7.5 | HAOr |

The rate in expressed nmol nitric oxide/min/mg protein, yield in % activity relative to total activity measured in the cell extract or soluble proteins, and purity fold is the specific activity per fraction compared to specific activity measured in cell extract or soluble proteins.

* Roman number indicates the replicate of soluble proteins from which proteins are further separated with fast protein liquid chromatography.

** For sample B values of fractions 9-17 were combined. For the specific activity and concentration the average of fractions 9-17 was used. For the other measurements the sum of the individual fractions was used.

*** For sample B1 values of fractions 11-14 were combined. For the specific activity and concentration the average of fractions 11-14 was used. For the other measurements the sum of the individual fractions was used.

**Supplementary table 2** - Proteins identified in sample B1 (low-resolution fractions 11-14) and low-resolution fractions 9 and 10 did not show any missing co-factors for identified nitrite reductase NirS. The positive Pearson correlation score calculated between specific activity measured per fraction and the relative protein abundance per fraction, showed that NirS is the only known active nitrite reductase contributing to nitric oxide production from nitrite in sample B. Other proteins that showed a positive correlation are not identified as nitrite reductases. Neither the low-resolution fractions 9 and 10 nor sample B1 contained known co-factors of NirS. Thus, the loss in nitrite reductase activity in sample B1 compared to sample B, is probably not due to missing co-factors. The data is ordered based on relative protein abundance that correlates best with the specific activity. Accession numbers refer to the *K. stuttgartiensis* protein sequence database in Uniprot (entry KSMBR1).

| Protein number | Accession | Description | Relative protein abundance per fraction | | | Pearson correlation score |
|---|---|---|---|---|---|---|
| | | | Low-resolution fraction 9 | Low-resolution fraction 10 | Sample B1 | between specific activity and relative protein abundance |
| 1 | Q1Q2D9 | Putative periplasmic serine endoprotease DegP-like | 2.50E+05 | 4.20E+05 | 5.94E+05 | 0.99 |
| 2 | A0A2C9CDH8 | Corrinoid/iron-sulfur protein large subunit | 3.61E+05 | 3.29E+05 | 5.25E+05 | 0.97 |
| 3 | Q1PYP7 | RNA-binding protein | 3.10E+05 | 4.03E+05 | 7.87E+05 | 0.94 |
| 4 | Q1Q1P1 | Putative succinyl-diaminopimelate desuccinylase | 1.13E+04 | 3.92E+04 | 5.55E+04 | 0.94 |
| 5 | A0A2C9CAZ0 | Uncharacterized protein | 1.81E+06 | 1.50E+06 | 2.02E+06 | 0.92 |
| **6** | **Q1Q4F5** | **Strongly similar to cd1 nitrite reductase NirS** | **7.50E+05** | **7.28E+05** | **8.24E+05** | **0.92** |
| 7 | Q1PZK9 | phphoribyl-AMP cyclohydrolase | 2.11E+05 | 3.87E+05 | 2.98E+05 | 0.86 |
| 8 | A0A2C9CBW4 | CBS domain-containing protein | 0.00E+00 | 1.73E+05 | 2.39E+05 | 0.85 |
| 9 | Q1PX48 | Hydroxylamine oxidoreductase , HOX | 2.42E+07 | 2.42E+07 | 2.30E+07 | 0.85 |
| 10 | Q1Q1F2 | ATP phphoribyltransferase | 0.00E+00 | 1.10E+05 | 1.49E+05 | 0.85 |
| 11 | A0A2C9CJS7 | Malate dehydrogenase | 0.00E+00 | 7.89E+04 | 1.93E+05 | 0.85 |
| 12 | A0A2C9CHI2 | Ornithine carbamoyltransferase | 0.00E+00 | 5.38E+04 | 6.65E+04 | 0.84 |
| 13 | Q1Q6H8 | Peptidyl-prolyl cis-trans isomerase C (Rotamase C) | 7.23E+04 | 6.74E+04 | 7.30E+04 | 0.82 |
| 14 | Q1Q4G9 | Hemerythrin domain-containing protein | 0.00E+00 | 1.89E+05 | 2.04E+05 | 0.82 |
| 15 | A0A2C9CBU0 | Sulfatase-modifying factor enzyme domain-containing protein | 0.00E+00 | 1.70E+05 | 2.34E+05 | 0.82 |
| 16 | Q1Q7H2 | Ribe-phphate pyrophphokinase | 2.33E+05 | 3.36E+05 | 2.13E+05 | 0.75 |
| 17 | Q1PUK7 | Formate--tetrahydrofolate ligase | 5.23E+04 | 6.60E+04 | 2.56E+05 | 0.75 |
| 18 | A0A2C9CJH3 | NAD/GMP synthase domain-containing protein | 1.77E+04 | 0.00E+00 | 5.53E+04 | 0.74 |
| 19 | Q1Q6M7 | FAD:protein FMN transferase | 1.23E+04 | 0.00E+00 | 5.66E+04 | 0.74 |
| 20 | A0A2C9CFH5 | Aminotransferase class V domain-containing protein | 1.55E+05 | 3.91E+04 | 1.78E+05 | 0.74 |
| 21 | Q1Q2X4 | 3-oxoacyl-[acyl-carrier-protein] synthase 2 | 0.00E+00 | 8.18E+03 | 2.16E+05 | 0.73 |
| 22 | A0A6G7GLF5 | S-layer protein | 1.42E+05 | 2.29E+05 | 1.54E+05 | 0.72 |
| 23 | Q1Q7P4 | Class IIb small soluble cyt c | 7.10E+06 | 8.85E+06 | 5.53E+06 | 0.72 |
| 24 | Q1Q2X1 | Putative bacteriochlorophyllide d C-12(1)-methyltransferase | 0.00E+00 | 0.00E+00 | 4.64E+04 | 0.72 |
| 25 | Q1Q2W3 | Carboxymuconolactone decarboxylase-like domain-containing protein | 0.00E+00 | 0.00E+00 | 3.66E+04 | 0.72 |
| 26 | Q1Q3H0 | ATP synthase subunit beta | 0.00E+00 | 0.00E+00 | 7.73E+05 | 0.72 |
| 27 | A0A2C9CJI3 | GatB/YqeY domain-containing protein | 0.00E+00 | 0.00E+00 | 7.48E+04 | 0.72 |
| 28 | Q1PW34 | Uncharacterized protein | 0.00E+00 | 0.00E+00 | 2.27E+04 | 0.72 |
| 29 | A0A2C9CH96 | Glutamine--tRNA ligase | 0.00E+00 | 0.00E+00 | 1.88E+04 | 0.72 |
| 30 | Q1Q201 | Integration ht factor subunit beta | 0.00E+00 | 0.00E+00 | 3.10E+04 | 0.72 |
| 31 | Q1PZD7 | Uncharacterized protein | 0.00E+00 | 0.00E+00 | 1.00E+04 | 0.72 |
| 32 | A0A2C9CEK8 | D-3-phphoglycerate dehydrogenase | 0.00E+00 | 0.00E+00 | 1.26E+04 | 0.72 |
| 33 | Q1PVP3 | 3-oxoacyl-[acyl-carrier-protein] reductase FabG | 0.00E+00 | 0.00E+00 | 1.04E+04 | 0.72 |
| 34 | A0A2C9CH02 | Bifunctional oligoribonuclease/PAP phphatase NrnA | 0.00E+00 | 0.00E+00 | 6.22E+03 | 0.72 |
| 35 | A0A2C9CIY9 | Flagellar hook-length control protein-like C-terminal domain-containing protein | 0.00E+00 | 0.00E+00 | 2.13E+04 | 0.72 |
| 36 | A0A2C9CC45 | Serine--tRNA ligase | 0.00E+00 | 0.00E+00 | 1.01E+04 | 0.72 |
| 37 | Q1Q3F2 | SWIM-type domain-containing protein | 0.00E+00 | 0.00E+00 | 6.22E+04 | 0.72 |
| 38 | Q1Q5R5 | Aspartokinase | 0.00E+00 | 0.00E+00 | 1.77E+04 | 0.72 |
| 39 | A0A2C9CGI1 | Peroxiredoxin | 0.00E+00 | 0.00E+00 | 8.01E+04 | 0.72 |
| 40 | A0A2C9CFM5 | Transcription termination factor Rho | 0.00E+00 | 0.00E+00 | 7.44E+04 | 0.72 |
| 41 | A0A2C9CJI9 | site-specific DNA-methyltransferase (adenine-specific) | 0.00E+00 | 0.00E+00 | 1.23E+05 | 0.72 |
| 42 | Q1Q0M4 | 2-isopropylmalate synthase | 0.00E+00 | 0.00E+00 | 2.89E+04 | 0.72 |
| 43 | Q1Q045 | 12,18-didecarboxysiroheme deacetylase | 0.00E+00 | 0.00E+00 | 3.29E+04 | 0.72 |
| 44 | Q1PW15 | ATP-dependent Clp protease proteolytic subunit | 0.00E+00 | 0.00E+00 | 7.19E+04 | 0.72 |
| 45 | A0A2C9CFE4 | Lon protease | 0.00E+00 | 0.00E+00 | 1.49E+05 | 0.72 |
| 46 | Q1PVQ6 | Uncharacterized protein | 0.00E+00 | 0.00E+00 | 2.33E+04 | 0.72 |
| 47 | A0A2C9CI21 | Aminotransferase | 0.00E+00 | 0.00E+00 | 3.21E+04 | 0.72 |
| 48 | A0A2C9CH82 | Flavodoxin-like domain-containing protein | 0.00E+00 | 0.00E+00 | 1.19E+04 | 0.72 |

| Protein number | Accession | Description | Relative protein abundance per fraction | | | Pearson correlation score between specific activity and relative protein abundance |
|---|---|---|---|---|---|---|
| | | | Low-resolution fraction 9 | Low-resolution fraction 10 | Sample B1 | |
| 49 | Q1Q418 | SHSP domain-containing protein | 0.00E+00 | 0.00E+00 | 1.03E+04 | 0.72 |
| 50 | A0A2C9CGR6 | Thiazole synthase | 0.00E+00 | 0.00E+00 | 5.62E+04 | 0.72 |
| 51 | Q1Q679 | 2-oxoacid:acceptor oxidoreductase subunit alpha | 0.00E+00 | 0.00E+00 | 7.35E+03 | 0.72 |
| 52 | A0A2C9CD15 | Probable glycine dehydrogenase (decarboxylating) subunit 2 | 0.00E+00 | 0.00E+00 | 1.82E+04 | 0.72 |
| 53 | Q1Q7H1 | 50S ribomal protein L25 | 0.00E+00 | 0.00E+00 | 3.82E+04 | 0.72 |
| 54 | A0A2C9CLF2 | PDZ domain-containing protein | 0.00E+00 | 0.00E+00 | 1.92E+04 | 0.72 |
| 55 | Q1PXR4 | Phphoenolpyruvate carboxylase | 0.00E+00 | 0.00E+00 | 1.91E+04 | 0.72 |
| 56 | Q1Q2K7 | Carbamoyl-phphate synthase large chain | 0.00E+00 | 0.00E+00 | 3.58E+05 | 0.72 |
| 57 | Q1Q2K6 | Probable glycine dehydrogenase (decarboxylating) subunit 1 | 0.00E+00 | 0.00E+00 | 2.21E+04 | 0.72 |
| 58 | A0A6G7GWM8 | Flotillin-like protein FloA | 0.00E+00 | 0.00E+00 | 2.25E+04 | 0.72 |
| 59 | Q1Q1N7 | DUF3326 domain-containing protein | 0.00E+00 | 0.00E+00 | 6.92E+03 | 0.72 |
| 60 | A0A2C9CE92 | Adenyluccinate lyase | 0.00E+00 | 0.00E+00 | 1.63E+04 | 0.72 |
| 61 | Q1Q4Q3 | 50S ribomal subunit assembly factor BipA | 0.00E+00 | 0.00E+00 | 2.29E+05 | 0.72 |
| 62 | Q1Q3W3 | Arginine--tRNA ligase | 0.00E+00 | 0.00E+00 | 6.65E+04 | 0.72 |
| 63 | A0A2C9CAP5 | 2-dehydro-3-deoxyphphooctonate aldolase | 0.00E+00 | 0.00E+00 | 5.86E+04 | 0.72 |
| 64 | A0A6G7GXY1 | Cytochrome c | 0.00E+00 | 0.00E+00 | 3.35E+04 | 0.72 |
| 65 | A0A2C9CFI7 | Histidine--tRNA ligase | 0.00E+00 | 0.00E+00 | 4.40E+04 | 0.72 |
| 66 | A0A2C9CAN1 | 2,3-bisphphoglycerate-dependent phphoglycerate mutase | 0.00E+00 | 0.00E+00 | 2.28E+04 | 0.72 |
| 67 | Q1Q4M3 | 5-hydroxybenzimidazole synthase | 0.00E+00 | 0.00E+00 | 8.49E+04 | 0.72 |
| 68 | A0A2C9CI02 | PilT/PilU family type 4a pilus ATPase | 0.00E+00 | 0.00E+00 | 3.58E+04 | 0.72 |
| 69 | A0A2C9CEZ2 | Aspartate--tRNA(Asp/Asn) ligase | 0.00E+00 | 0.00E+00 | 9.86E+04 | 0.72 |
| 70 | A0A6G7GRR4 | peptide-methionine (S)-S-oxide reductase | 0.00E+00 | 0.00E+00 | 4.99E+04 | 0.72 |
| 71 | A0A2C9CE81 | Isoleucine--tRNA ligase | 0.00E+00 | 0.00E+00 | 8.47E+04 | 0.72 |
| 72 | A0A6G7GPV9 | PTS sugar transporter subunit IIA | 0.00E+00 | 0.00E+00 | 3.83E+04 | 0.72 |
| 73 | Q1PZ66 | DUF1858 domain-containing protein | 3.00E+05 | 4.10E+04 | 6.95E+05 | 0.69 |
| 74 | Q1Q3E2 | Amidohydrolase-related domain-containing protein | 0.00E+00 | 3.22E+04 | 2.01E+04 | 0.67 |
| 75 | A0A2C9CKW5 | Porphobilinogen deaminase | 1.35E+05 | 2.12E+05 | 1.09E+05 | 0.66 |
| 76 | Q1PW16 | Trigger factor | 2.79E+05 | 1.35E+05 | 1.66E+06 | 0.65 |
| 77 | Q1PZN3 | Chaperone protein HtpG | 3.08E+04 | 1.12E+05 | 6.81E+05 | 0.65 |
| 78 | Q1Q3V0 | Phpho-2-dehydro-3-deoxyheptonate aldolase | 0.00E+00 | 0.00E+00 | 5.35E+05 | 0.64 |
| 79 | Q1Q105 | 3-isopropylmalate dehydratase small subunit | 4.25E+04 | 0.00E+00 | 5.14E+04 | 0.63 |
| 80 | A0A2C9CKK5 | Uncharacterized protein | 1.29E+04 | 4.27E+04 | 2.24E+05 | 0.63 |
| 81 | A0A2C9CKK3 | Peptide methionine sulfoxide reductase MsrA | 0.00E+00 | 0.00E+00 | 1.17E+05 | 0.63 |
| 82 | A0A2C9CFF5 | PTS sugar transporter subunit IIA | 0.00E+00 | 0.00E+00 | 6.31E+05 | 0.56 |
| 83 | Q1Q4L7 | TIGR04255 family protein | 0.00E+00 | 0.00E+00 | 3.51E+04 | 0.55 |
| 84 | Q1Q7P3 | Class I small soluble cyt c | 4.73E+06 | 4.07E+06 | 2.28E+06 | 0.53 |
| 85 | Q1PUP9 | Adenylhomocysteinase | 0.00E+00 | 0.00E+00 | 1.99E+06 | 0.52 |
| 86 | A0A2C9CFG1 | Chaperonin GroEL | 8.43E+04 | 4.04E+04 | 1.75E+05 | 0.51 |
| 87 | Q1PYX1 | ATP-dependent Clp protease proteolytic subunit | 0.00E+00 | 0.00E+00 | 2.35E+05 | 0.50 |
| 88 | Q1PZK4 | Chaperonin GroEL | 3.11E+04 | 0.00E+00 | 5.43E+04 | 0.50 |
| 89 | A0A2C9CDZ7 | S-layer protein | 2.25E+05 | 3.36E+05 | 1.15E+05 | 0.49 |
| 90 | Q1PVI8 | MoxR family ATPase | 0.00E+00 | 0.00E+00 | 6.55E+04 | 0.49 |
| 91 | Q1Q2R8 | 3-hydroxy-5-phphonooxypentane-2,4-dione thiolase | 1.39E+05 | 4.81E+04 | 8.04E+04 | 0.49 |
| 92 | A0A2C9CN84 | Radical SAM core domain-containing protein | 0.00E+00 | 0.00E+00 | 1.54E+05 | 0.49 |
| 93 | Q1PYI1 | Iron-containing redox enzyme family protein | 5.25E+06 | 4.69E+06 | 2.24E+06 | 0.48 |
| 94 | Q1PZD5 | Nitrite oxidoreductase subunit B | 7.13E+04 | 4.66E+05 | 2.34E+06 | 0.47 |
| 95 | A0A2C9CKI8 | transketolase | 6.88E+04 | 7.76E+04 | 2.67E+04 | 0.44 |
| 96 | A0A2C9CC12 | Histidinol-phphate aminotransferase | 4.55E+05 | 1.70E+05 | 2.08E+05 | 0.43 |
| 97 | Q1Q637 | DNA-binding protein | 4.61E+04 | 1.38E+05 | 8.70E+04 | 0.43 |
| 98 | Q1Q7J1 | Putative hydroxylamine oxidoreductase hao | 1.45E+04 | 1.06E+05 | 1.77E+05 | 0.40 |
| 99 | Q1PZD8 | Nitrite oxidoreductase subunit A | 3.53E+05 | 1.39E+06 | 7.02E+06 | 0.39 |
| 100 | A0A2C9CHV5 | PQQ-like beta-propeller repeat protein | 8.88E+04 | 0.00E+00 | 5.05E+04 | 0.37 |
| 101 | A0A2C9CH14 | Hydrazine synthase subunit B | 8.49E+07 | 4.62E+07 | 3.14E+07 | 0.34 |
| 102 | Q1Q4Z4 | Homerine dehydrogenase | 1.97E+05 | 0.00E+00 | 1.03E+05 | 0.33 |
| 103 | Q1Q2X3 | Similar to beta-ketoacyl acyl carrier protein synthase II | 0.00E+00 | 1.56E+05 | 1.32E+06 | 0.33 |

| Protein number | Accession | Description | Relative protein abundance per fraction | | | Pearson correlation score between specific activity and relative protein abundance |
|---|---|---|---|---|---|---|
| | | | Low-resolution fraction 9 | Low-resolution fraction 10 | Sample B1 | |
| 104 | Q1Q3Y3 | Purine nucleide phphorylase | 0.00E+00 | 0.00E+00 | 1.34E+05 | 0.33 |
| 105 | A0A2C9CF24 | Uncharacterized protein | 0.00E+00 | 0.00E+00 | 1.36E+05 | 0.30 |
| 106 | A0A2C9CD37 | Peptide methionine sulfoxide reductase MsrA | 7.63E+05 | 2.94E+04 | 3.17E+05 | 0.28 |
| 107 | A0A2C9CEF3 | Putative NADH dehydrogenase I chain F (1st module) | 0.00E+00 | 0.00E+00 | 2.97E+05 | 0.27 |
| 108 | Q1PUT2 | PDZ domain-containing protein | 0.00E+00 | 0.00E+00 | 1.21E+05 | 0.26 |
| 109 | A0A2C9CHN2 | Hydrazine synthase subunit A | 1.16E+08 | 5.16E+07 | 3.49E+07 | 0.26 |
| 110 | A0A2C9CCP8 | PDZ domain-containing protein | 0.00E+00 | 0.00E+00 | 1.15E+05 | 0.25 |
| 111 | A0A2C9CDK8 | Gamma-glutamyl phphate reductase | 0.00E+00 | 0.00E+00 | 5.68E+04 | 0.25 |
| 112 | Q1PZD4 | Nitrite oxidoreductase subunit C | 1.55E+05 | 5.51E+05 | 2.63E+06 | 0.24 |
| 113 | A0A2C9CHM2 | Hydrazine synthase subunit C | 1.01E+08 | 4.28E+07 | 2.96E+07 | 0.24 |
| 114 | A0A2C9CBX8 | Glutamine synthetase | 6.41E+04 | 2.60E+04 | 1.49E+04 | 0.24 |
| 115 | Q1PV11 | DUF2024 domain-containing protein | 0.00E+00 | 0.00E+00 | 8.14E+04 | 0.23 |
| 116 | Q1Q2J4 | Elongation factor Ts | 2.26E+05 | 4.03E+04 | 6.69E+04 | 0.23 |
| 117 | Q1Q277 | Uncharacterized protein | 0.00E+00 | 1.68E+04 | 0.00E+00 | 0.21 |
| 118 | Q1PXN5 | Chemotaxis protein CheY | 0.00E+00 | 2.17E+04 | 0.00E+00 | 0.21 |
| 119 | Q1Q666 | Putative transcriptional repressor | 0.00E+00 | 1.15E+04 | 0.00E+00 | 0.21 |
| 120 | A0A2C9CDI4 | Glycerate 2-kinase | 0.00E+00 | 2.07E+04 | 0.00E+00 | 0.21 |
| 121 | A0A2C9CF13 | Chaperonin GroEL | 4.40E+04 | 0.00E+00 | 6.82E+04 | 0.21 |
| 122 | Q1Q3W5 | Glutamate synthase (NADPH) large chain | 7.54E+04 | 1.77E+04 | 1.48E+04 | 0.16 |
| 123 | Q1PVE2 | Type-1 blue copper-containing cupredoxin | 1.22E+05 | 3.76E+04 | 1.94E+04 | 0.15 |
| 124 | Q1PZC8 | Putative septation protein SpoVG | 8.78E+04 | 5.86E+04 | 5.09E+04 | 0.14 |
| 125 | Q1Q354 | Sulfate adenylyltransferase | 0.00E+00 | 0.00E+00 | 1.58E+06 | 0.13 |
| 126 | Q1PZK3 | Co-chaperonin GroES | 1.21E+06 | 4.33E+05 | 6.08E+05 | 0.11 |
| 127 | A0A2C9CHX8 | Dihydroxy-acid dehydratase | 0.00E+00 | 0.00E+00 | 2.22E+05 | 0.11 |
| 128 | Q1Q4H1 | Heme d1 biynthesis protein NirF | 1.26E+04 | 8.93E+03 | 0.00E+00 | 0.10 |
| 129 | Q1Q1A9 | Argininuccinate synthase | 3.45E+04 | 0.00E+00 | 6.95E+03 | 0.10 |
| 130 | A0A2C9CGP5 | Flavodoxin family protein | 0.00E+00 | 0.00E+00 | 3.54E+04 | 0.09 |
| 131 | A0A2C9CDL6 | Uncharacterized protein | 0.00E+00 | 0.00E+00 | 1.10E+06 | 0.07 |
| 132 | A0A2C9CE26 | Acetolactate synthase | 3.57E+05 | 1.28E+05 | 1.88E+04 | 0.07 |
| 133 | Q1PVQ3 | Hsp20/alpha crystallin family protein | 7.83E+05 | 2.75E+05 | 4.25E+04 | 0.07 |
| 134 | Q1PW67 | Peptidylprolyl isomerase | 2.30E+06 | 4.61E+05 | 4.18E+05 | 0.06 |
| 135 | A0A6G7GTJ9 | Uncharacterized protein | 9.51E+05 | 2.28E+05 | 2.45E+04 | 0.02 |
| 136 | A0A2C9CKL5 | Uncharacterized protein | 4.77E+04 | 1.16E+04 | 0.00E+00 | 0.00 |
| 137 | A0A2C9CHG6 | CBS domain-containing protein | 0.00E+00 | 0.00E+00 | 2.09E+04 | 0.00 |
| 138 | Q1PXW3 | Lon protease | 0.00E+00 | 0.00E+00 | 5.23E+05 | -0.01 |
| 139 | Q1PY42 | Co-chaperonin GroES | 2.11E+06 | 3.01E+05 | 0.00E+00 | -0.02 |
| 140 | Q1PWY3 | Chaperone protein ClpB | 0.00E+00 | 0.00E+00 | 8.57E+05 | -0.03 |
| 141 | Q1PW64 | Zinc ribbon domain protein | 1.34E+05 | 1.18E+04 | 0.00E+00 | -0.03 |
| 142 | Q1PYI0 | FMN-binding glutamate synthase family protein | 7.04E+04 | 6.08E+03 | 0.00E+00 | -0.03 |
| 143 | Q1Q1A6 | Uncharacterized protein | 9.74E+05 | 6.01E+04 | 0.00E+00 | -0.04 |
| 144 | A0A2C9CGA0 | Hsp20/alpha crystallin family protein | 7.15E+05 | 1.01E+04 | 7.85E+03 | -0.04 |
| 145 | A0A2C9CEW4 | transketolase | 2.99E+05 | 1.08E+04 | 0.00E+00 | -0.04 |
| 146 | A0A2C9CC59 | Bifunctional purine biynthesis protein PurH | 2.03E+05 | 5.32E+03 | 0.00E+00 | -0.05 |
| 147 | A0A2C9CDL0 | Beta-ketoacyl-[acyl-carrier-protein] synthase III | 0.00E+00 | 0.00E+00 | 1.47E+05 | -0.05 |
| 148 | A0A2C9CID2 | Acetylornithine aminotransferase | 1.67E+05 | 0.00E+00 | 0.00E+00 | -0.05 |
| 149 | Q1PZI4 | Nucleide diphphate kinase | 7.81E+04 | 0.00E+00 | 0.00E+00 | -0.05 |
| 150 | A0A2C9CBY9 | uroporphyrinogen-III C-methyltransferase | 2.38E+05 | 0.00E+00 | 0.00E+00 | -0.05 |
| 151 | Q1PVG3 | Cyclic nucleotide-binding domain-containing protein | 2.74E+04 | 0.00E+00 | 0.00E+00 | -0.05 |
| 152 | A0A2C9CA80 | DUF1326 domain-containing protein | 2.39E+05 | 0.00E+00 | 0.00E+00 | -0.05 |
| 153 | A0A2C9CIP1 | Flagellar hook protein FlgE | 6.12E+04 | 0.00E+00 | 0.00E+00 | -0.05 |
| 154 | Q1PXI4 | Uncharacterized protein | 1.73E+05 | 0.00E+00 | 0.00E+00 | -0.05 |
| 155 | A0A2C9CEZ1 | Transpase | 2.82E+05 | 0.00E+00 | 0.00E+00 | -0.05 |
| 156 | Q1Q3D6 | Carboxypeptidase regulatory-like domain-containing protein | 2.68E+04 | 0.00E+00 | 0.00E+00 | -0.05 |
| 157 | A0A2C9CLC9 | 6-phphogluconate dehydrogenase, decarboxylating | 5.95E+05 | 0.00E+00 | 0.00E+00 | -0.05 |
| 158 | Q1Q338 | Adenylyl-sulfate reductase subunit alpha | 3.92E+04 | 0.00E+00 | 0.00E+00 | -0.05 |

| Protein number | Accession | Description | Relative protein abundance per fraction Low-resolution fraction 9 | Low-resolution fraction 10 | Sample B1 | Pearson correlation score between specific activity and relative protein abundance |
|---|---|---|---|---|---|---|
| 159 | A0A2C9CDR5 | Strongly similar to sigma 54 formate hydrogen lyase transcriptional activator FlhA | 6.29E+04 | 0.00E+00 | 0.00E+00 | -0.05 |
| 160 | Q1Q416 | Small heat shock protein C2 | 5.58E+04 | 0.00E+00 | 0.00E+00 | -0.05 |
| 161 | A0A2C9CK20 | Valine--tRNA ligase | 1.75E+05 | 0.00E+00 | 0.00E+00 | -0.05 |
| 162 | Q1PX47 | Tryptophan synthase beta chain | 1.29E+04 | 0.00E+00 | 0.00E+00 | -0.05 |
| 163 | A0A2C9CDU6 | Uncharacterized protein | 4.19E+04 | 0.00E+00 | 0.00E+00 | -0.05 |
| 165 | Q1Q2L2 | Gluce-1-phphate thymidylyltransferase | 8.78E+04 | 0.00E+00 | 0.00E+00 | -0.05 |
| 166 | A0A6G7GL09 | site-specific DNA-methyltransferase (adenine-specific) | 2.81E+05 | 0.00E+00 | 0.00E+00 | -0.05 |
| 167 | Q1Q7C6 | Similar to ATP dependent transcriptional activator | 3.95E+05 | 0.00E+00 | 0.00E+00 | -0.05 |
| 168 | Q1Q0T9 | Hypothetical (Triheme) protein | 1.26E+04 | 0.00E+00 | 0.00E+00 | -0.05 |
| 169 | Q1PV60 | PEGA domain-containing protein | 1.15E+04 | 0.00E+00 | 0.00E+00 | -0.05 |
| 170 | A0A2C9CHS3 | 3-isopropylmalate dehydratase large subunit | 5.87E+04 | 0.00E+00 | 0.00E+00 | -0.05 |
| 171 | Q1Q0A1 | Putative serine-proteinase HtrA/ DegQ/ DegS family | 2.15E+04 | 0.00E+00 | 0.00E+00 | -0.05 |
| 172 | A0A2C9CHF8 | Thioredoxin domain-containing protein | 2.25E+04 | 0.00E+00 | 0.00E+00 | -0.05 |
| 173 | A0A2C9CAW2 | MULTIHEME_CYTC domain-containing protein | 2.33E+04 | 0.00E+00 | 0.00E+00 | -0.05 |
| 174 | Q1PZA6 | Amidohydrolase | 1.12E+05 | 0.00E+00 | 0.00E+00 | -0.05 |
| 175 | A0A2C9CCG2 | Uncharacterized protein | 6.25E+04 | 0.00E+00 | 0.00E+00 | -0.05 |
| 176 | A0A2C9CB13 | Strongly similar to pyruvate:ferredoxin oxidoreductase | 6.11E+04 | 0.00E+00 | 0.00E+00 | -0.05 |
| 177 | A0A2C9CC73 | UDP-N-acetylmuramoylalanine--D-glutamate ligase | 3.92E+06 | 0.00E+00 | 0.00E+00 | -0.05 |
| 178 | Q1Q697 | Quinolinate synthase | 6.19E+04 | 0.00E+00 | 0.00E+00 | -0.05 |
| 179 | Q1Q6A7 | Cold-shock protein | 4.74E+04 | 9.31E+04 | 1.48E+06 | -0.06 |
| 180 | Q1Q133 | Transcription termination/antitermination protein NusG | 1.98E+05 | 4.62E+04 | 1.59E+04 | -0.08 |
| 181 | Q1Q3G0 | PhoU domain-containing protein | 0.00E+00 | 0.00E+00 | 2.55E+05 | -0.13 |
| 182 | Q1Q2D8 | Cysteine desulfurase | 0.00E+00 | 0.00E+00 | 7.00E+04 | -0.19 |
| 183 | Q1Q123 | Elongation factor Tu | 9.03E+04 | 7.68E+04 | 6.05E+06 | -0.21 |
| 184 | A0A2C9CHA0 | 50S ribomal protein L7/L12 | 0.00E+00 | 7.62E+03 | 3.10E+05 | -0.28 |
| 185 | Q1Q1C6 | PQQ-like beta-propeller repeat protein | 0.00E+00 | 0.00E+00 | 7.78E+04 | -0.30 |
| 186 | A0A2C9CBQ3 | Chaperone protein DnaK | 0.00E+00 | 3.23E+04 | 4.48E+06 | -0.31 |
| 187 | A0A2C9CJE3 | Glyco_hydro_cc domain-containing protein | 0.00E+00 | 0.00E+00 | 4.78E+05 | -0.33 |
| 188 | Q1PYW4 | Carbohydrate kinase family protein | 0.00E+00 | 0.00E+00 | 5.28E+04 | -0.37 |
| 189 | Q1Q1N9 | Putative inine-5'-monophphate dehydrogenase related protein II | 0.00E+00 | 0.00E+00 | 6.31E+05 | -0.37 |
| 190 | Q1PY51 | Scaffold protein | 0.00E+00 | 0.00E+00 | 6.48E+04 | -0.40 |
| 191 | A0A2C9CD92 | Glutamate--tRNA ligase | 0.00E+00 | 0.00E+00 | 6.58E+04 | -0.41 |
| 192 | A0A2C9CB61 | Cofactor-independent phphoglycerate mutase | 0.00E+00 | 0.00E+00 | 1.34E+05 | -0.41 |
| 193 | Q1PUL0 | Porin | 0.00E+00 | 4.03E+04 | 2.09E+05 | -0.42 |
| 194 | Q1Q1N4 | Hypothetical phphotransacetylase protein | 0.00E+00 | 0.00E+00 | 6.10E+04 | -0.43 |
| 195 | Q1PVQ5 | Putative superoxide reductase | 5.67E+04 | 6.62E+04 | 4.10E+05 | -0.45 |
| 196 | Q1Q4X4 | HlyD_D23 domain-containing protein | 0.00E+00 | 0.00E+00 | 6.35E+04 | -0.48 |
| 197 | A0A2C9CKJ6 | Exported protein | 5.89E+04 | 6.35E+04 | 1.01E+05 | -0.56 |
| 198 | A0A2C9CBQ7 | Uncharacterized protein | 0.00E+00 | 0.00E+00 | 1.27E+05 | -0.59 |
| 199 | A0A2C9CKT8 | Sulfatase-modifying factor enzyme domain-containing protein | 0.00E+00 | 0.00E+00 | 1.43E+05 | -0.63 |
| 200 | Q1PXG4 | Ferritin family protein | 0.00E+00 | 0.00E+00 | 1.88E+05 | -0.66 |
| 201 | Q1PXG5 | Thioredoxin reductase | 0.00E+00 | 0.00E+00 | 3.61E+04 | -0.66 |
| 202 | Q1PYT0 | Enolase | 0.00E+00 | 0.00E+00 | 1.27E+05 | -0.69 |
| 203 | A0A2C9CLM3 | YfdX protein | 0.00E+00 | 0.00E+00 | 2.85E+05 | -0.69 |
| 204 | Q1Q7P0 | Uncharacterized protein | 8.31E+03 | 0.00E+00 | 2.70E+05 | -0.72 |
| 205 | Q1PXY6 | phphoenolpyruvate--protein phphotransferase | 0.00E+00 | 0.00E+00 | 9.09E+03 | -0.72 |
| 206 | A0A2C9CBT5 | Riboflavin biynthesis protein RibBA | 0.00E+00 | 0.00E+00 | 1.61E+04 | -0.74 |
| 207 | Q1Q3K7 | Elongation factor P | 0.00E+00 | 0.00E+00 | 3.10E+04 | -0.74 |
| 208 | Q1PUP8 | MoaD/ThiS family protein | 0.00E+00 | 0.00E+00 | 4.55E+04 | -0.76 |
| 209 | Q1Q5R4 | Energy-dependent translational throttle protein EttA | 0.00E+00 | 0.00E+00 | 4.23E+04 | -0.77 |
| 210 | Q1Q303 | Exported protein | 0.00E+00 | 0.00E+00 | 4.61E+05 | -0.77 |
| 211 | Q1Q7G9 | 30S ribomal protein S6 | 0.00E+00 | 0.00E+00 | 2.24E+04 | -0.78 |
| 212 | A0A2C9CIY0 | Proline--tRNA ligase | 0.00E+00 | 0.00E+00 | 4.96E+04 | -0.79 |
| 213 | Q1Q141 | Elongation factor G | 6.11E+04 | 0.00E+00 | 3.26E+05 | -0.80 |
| 214 | Q1Q0K4 | NIF system FeS cluster assembly NifU C-terminal domain-containing protein | 0.00E+00 | 0.00E+00 | 2.08E+05 | -0.81 |

| Protein number | Accession | Description | Relative protein abundance per fraction | | | Pearson correlation score |
|---|---|---|---|---|---|---|
| | | | Low-resolution fraction 9 | Low-resolution fraction 10 | Sample B1 | between specific activity and relative protein abundance |
| 215 | Q1PYS0 | Fructe-1,6-bisphphatase class 1 | 0.00E+00 | 0.00E+00 | 5.67E+04 | -0.81 |
| 216 | Q1PYY4 | NAD-dependent formate dehydrogenase alpha subunit / selenocysteine-containing | 0.00E+00 | 0.00E+00 | 1.79E+05 | -0.82 |
| 217 | Q1PW50 | FAD-dependent oxidoreductase | 0.00E+00 | 0.00E+00 | 1.31E+05 | -0.82 |
| 218 | A0A2C9CIU1 | Glutaredoxin domain-containing protein | 0.00E+00 | 0.00E+00 | 1.90E+04 | -0.83 |
| 219 | Q1PY41 | Chaperonin GroEL | 8.30E+04 | 0.00E+00 | 1.97E+05 | -0.84 |
| 220 | A0A2C9CHJ5 | FmdE domain-containing protein | 0.00E+00 | 0.00E+00 | 8.77E+04 | -0.85 |
| 221 | Q1PW49 | NADH-quinone oxidoreductase subunit NuoF | 0.00E+00 | 0.00E+00 | 3.79E+04 | -0.85 |
| 222 | A0A2C9CDQ2 | Strongly similar to proton-translocating NADH dehydrogenase I, 51 kDa subunit (NuoF) | 0.00E+00 | 0.00E+00 | 1.33E+05 | -0.85 |
| 223 | A0A2C9CBP2 | Strongly similar to proton-translocating NADH dehydrogenase I, 51 kDa subunit (NuoF) | 0.00E+00 | 0.00E+00 | 1.33E+05 | -0.85 |
| 224 | Q1Q0F8 | Exported protein | 0.00E+00 | 0.00E+00 | 1.72E+04 | -0.86 |
| 225 | A0A2C9CCN2 | Uncharacterized protein | 0.00E+00 | 0.00E+00 | 1.24E+04 | -0.87 |
| 226 | Q1Q6R3 | Strongly similar to to proton-translocating NADH dehydrogenase I, 24 kDa subunit (NuoE) | 0.00E+00 | 0.00E+00 | 1.08E+04 | -0.87 |
| 265 | Q1PV06 | Protein tyrine phphatase family protein | 7.45E+04 | 1.70E+04 | 0.00E+00 | -0.97 |

**Supplementary table 3** - Correlation between the relative abundance of identified proteins and specific nitrite reductase activity in the fractions obtained from sample B1 that were further separated on a high-resolution anion-exchanger. Pearson correlation score was calculated between the specific activity measured per fraction and the relative protein abundance measured per fraction. The data is ordered based on relative protein abundance that correlated best with the specific activity. The relative abundance of NirS correlated well with specific nitrite reductase activity. Moreover, NirS had the highest relative abundance measured in most active Fraction 9. Other proteins that showed a positive correlation are not identified as nitrite reductases. Accession numbers refer to the *K. stuttgartiensis* protein sequence database in Uniprot (entry KSMBR1).

| Protein number | Accession | Description | Relative protein abundance per fraction | | | | Pearson correlation score |
|---|---|---|---|---|---|---|---|
| | | | Fraction 7 | Fraction 8 | Fraction 9 | Fraction 10 | between specific activity and relative protein abundance |
| **1** | **Q1Q4F5** | **Strongly similar to cd1 nitrite reductase NirS** | **2.76E+06** | **2.90E+07** | **4.38E+07** | **3.01E+07** | **0.87** |
| 2 | A0A2C9CAZ0 | Uncharacterized protein | 0.00E+00 | 1.99E+03 | 1.46E+04 | 0.00E+00 | 0.77 |
| 3 | Q1Q3W5 | Glutamate synthase (NADPH) large chain | 0.00E+00 | 7.97E+04 | 9.63E+05 | 7.21E+04 | 0.74 |
| 4 | Q1PYI7 | Protein RecA | 0.00E+00 | 0.00E+00 | 5.76E+03 | 0.00E+00 | 0.69 |
| 5 | A0A2C9CN84 | Radical SAM core domain-containing protein | 0.00E+00 | 0.00E+00 | 1.13E+04 | 0.00E+00 | 0.69 |
| 6 | Q1PYI1 | Iron-containing redox enzyme family protein | 0.00E+00 | 0.00E+00 | 1.33E+05 | 0.00E+00 | 0.69 |
| 7 | Q1Q2D9 | Putative periplasmic serine endoprotease DegP-like | 0.00E+00 | 0.00E+00 | 1.23E+05 | 0.00E+00 | 0.69 |
| 8 | Q1PXC8 | Peptidylprolyl isomerase | 0.00E+00 | 0.00E+00 | 6.99E+05 | 1.22E+04 | 0.69 |
| 9 | Q1PW15 | ATP-dependent Clp protease proteolytic subunit | 4.77E+04 | 4.29E+04 | 1.24E+05 | 0.00E+00 | 0.64 |
| 10 | Q1PX48 | Hydroxylamine oxidoreductase | 3.46E+06 | 8.54E+06 | 4.67E+06 | 3.23E+06 | 0.60 |
| 11 | Q1Q6A7 | Cold-shock protein | 1.52E+05 | 1.29E+05 | 2.74E+05 | 0.00E+00 | 0.55 |
| 12 | Q1PZC8 | Putative septation protein SpoVG | 9.76E+04 | 1.01E+05 | 1.37E+05 | 2.03E+04 | 0.55 |
| 13 | Q1Q338 | Adenylyl-sulfate reductase subunit alpha | 0.00E+00 | 3.74E+04 | 0.00E+00 | 0.00E+00 | 0.40 |
| 14 | Q1Q201 | Integration ht factor subunit beta | 1.02E+05 | 9.95E+04 | 2.73E+05 | 2.90E+05 | 0.27 |
| 15 | Q1PZD5 | Nitrite oxidoreductase subunit B | 5.08E+06 | 8.28E+06 | 3.18E+06 | 8.15E+05 | 0.22 |
| 16 | Q1Q123 | Elongation factor Tu | 4.66E+06 | 4.65E+06 | 3.99E+06 | 2.06E+06 | 0.16 |
| 17 | Q1PZD8 | Nitrite oxidoreductase subunit A | 2.00E+07 | 2.97E+07 | 1.09E+07 | 3.63E+06 | 0.13 |
| 18 | A0A2C9CF13 | Chaperonin GroEL | 5.76E+04 | 5.84E+04 | 3.61E+04 | 0.00E+00 | 0.08 |
| 19 | A0A2C9CBQ3 | Chaperone protein DnaK | 2.70E+05 | 2.03E+05 | 1.92E+05 | 0.00E+00 | 0.03 |
| 20 | Q1Q7P0 | Uncharacterized protein | 1.20E+05 | 8.92E+04 | 8.57E+04 | 0.00E+00 | 0.03 |
| 21 | Q1PZD4 | Nitrite oxidoreductase subunit C | 8.14E+06 | 9.33E+06 | 3.96E+06 | 4.99E+05 | 0.01 |
| 22 | A0A2C9CFG1 | Chaperonin GroEL | 6.26E+04 | 5.84E+04 | 3.61E+04 | 0.00E+00 | 0.01 |
| 23 | Q1PY41 | Chaperonin GroEL | 1.21E+05 | 9.60E+04 | 7.75E+04 | 0.00E+00 | 0.00 |
| 24 | Q1Q0K4 | NIF system FeS cluster assembly NifU C-terminal domain-containing protein | 2.81E+04 | 0.00E+00 | 2.86E+04 | 0.00E+00 | -0.05 |
| 25 | A0A2C9CHN2 | Hydrazine synthase subunit A | 1.46E+06 | 1.43E+06 | 7.35E+05 | 1.79E+05 | -0.08 |
| 26 | Q1PVQ5 | Putative superoxide reductase | 6.47E+04 | 4.25E+04 | 4.04E+04 | 0.00E+00 | -0.09 |
| 27 | A0A2C9CH14 | Hydrazine synthase subunit B | 1.69E+06 | 1.54E+06 | 7.27E+05 | 1.64E+05 | -0.16 |
| 28 | Q1Q3G0 | PhoU domain-containing protein | 3.58E+04 | 4.70E+04 | 0.00E+00 | 0.00E+00 | -0.17 |
| 29 | A0A2C9CHM2 | Hydrazine synthase subunit C | 2.51E+06 | 2.06E+06 | 9.71E+05 | 2.05E+05 | -0.23 |
| 30 | A0A2C9CKJ6 | Exported protein | 6.26E+04 | 3.91E+04 | 2.31E+04 | 0.00E+00 | -0.31 |
| 31 | Q1PXW3 | Lon protease | 2.74E+05 | 2.22E+05 | 2.74E+04 | 0.00E+00 | -0.37 |
| 32 | Q1PYX1 | ATP-dependent Clp protease proteolytic subunit | 1.78E+05 | 1.10E+05 | 0.00E+00 | 0.00E+00 | -0.53 |
| 33 | A0A2C9CLM3 | YfdX protein | 1.24E+07 | 7.39E+06 | 3.19E+05 | 1.16E+04 | -0.53 |
| 34 | A0A2C9CDL6 | Uncharacterized protein | 2.22E+06 | 7.55E+05 | 4.48E+05 | 0.00E+00 | -0.57 |
| 35 | Q1PZD7 | Uncharacterized protein | 8.53E+04 | 2.54E+04 | 0.00E+00 | 0.00E+00 | -0.69 |
| 36 | A0A2C9CHG6 | CBS domain-containing protein | 4.53E+04 | 0.00E+00 | 0.00E+00 | 0.00E+00 | -0.77 |
| 37 | Q1PYT0 | Enolase | 4.59E+04 | 0.00E+00 | 0.00E+00 | 0.00E+00 | -0.77 |
| 38 | A0A2C9CDQ2 | Strongly similar to proton-translocating NADH dehydrogenase I, (NuoF) | 1.17E+04 | 0.00E+00 | 0.00E+00 | 0.00E+00 | -0.77 |
| 39 | A0A2C9CBP2 | Strongly similar to proton-translocating NADH dehydrogenase I, (NuoF) | 1.17E+04 | 0.00E+00 | 0.00E+00 | 0.00E+00 | -0.77 |
| 40 | A0A2C9CCN2 | Uncharacterized protein | 7.63E+04 | 0.00E+00 | 0.00E+00 | 0.00E+00 | -0.77 |

**Supplementary table 4** - Correlation between the relative abundance of identified proteins and specific nitrite reductase activity in three samples obtained with low-resolution column chromatography: FT, sample A, and low-resolution fraction 6. Pearson correlation score was calculated between the specific activity measured per fraction and the relative protein abundance measured per fraction. The data is ordered based on relative protein abundance that correlated best with the specific activity. None of the proteins that showed a positive correlation are identified as nitrite reductases. Accession numbers refer to the *K. stuttgartiensis* protein sequence database in Uniprot (entry KSMBR1).

| Protein number | Accession | Description | Relative protein abundance per fraction | | | Pearson correlation score |
|---|---|---|---|---|---|---|
| | | | FT | sample A | Low-resolution fraction 6 | between specific activity and relative protein abundance |
| 1 | Q1PYG3 | Adenylyl-sulfate kinase | 0.00E+00 | 3.95E+01 | 2.77E+01 | 1.00 |
| 2 | Q1PXM5 | Chemotaxis protein CheY-like protein | 0.00E+00 | 5.53E+03 | 8.16E+01 | 1.00 |
| 3 | A0A2C9CD35 | Methylthioribose-1-phosphate isomerase | 0.00E+00 | 5.24E+02 | 1.58E+02 | 1.00 |
| 4 | Q1Q1E9 | Protein-arginine kinase | 0.00E+00 | 1.36E+02 | 9.98E+01 | 1.00 |
| 5 | Q1Q243 | Putative deoxyribonuclease YcfH / radical SAM domain protein | 0.00E+00 | 3.68E+02 | 7.53E+01 | 1.00 |
| 6 | Q1PUP8 | Putative molybdopterin synthase subunit 1 | 9.57E+00 | 0.00E+00 | 1.27E+01 | 1.00 |
| 7 | Q1Q0C4 | 2-amino-4-hydroxy-6-hydroxymethyldihydropteridine diphosphokinase | 0.00E+00 | 3.48E+02 | 3.03E+02 | 1.00 |
| 8 | Q1Q7G3 | 3-isopropylmalate dehydrogenase | 0.00E+00 | 1.22E+02 | 6.38E+01 | 1.00 |
| 9 | Q1Q7G2 | AMMECR1 domain-containing protein | 0.00E+00 | 5.21E+01 | 2.50E+01 | 1.00 |
| 10 | Q1PYX1 | ATP-dependent Clp protease proteolytic subunit | 0.00E+00 | 2.46E+02 | 1.85E+02 | 1.00 |
| 11 | Q1Q504 | Branched-chain-amino-acid aminotransferase | 0.00E+00 | 5.00E+01 | 1.76E+01 | 1.00 |
| 12 | Q1PYY7 | Carbon monoxide dehydrogenase accessory protein CooC | 0.00E+00 | 5.34E+01 | 1.90E+01 | 1.00 |
| 13 | Q1PZB8 | Chemotaxis protein CheY | 0.00E+00 | 3.86E+02 | 2.46E+02 | 1.00 |
| 14 | Q1Q516 | Diheme cytochrome c | 0.00E+00 | 3.73E+03 | 2.70E+03 | 1.00 |
| 15 | Q1Q0K3 | DUF1858 domain-containing protein | 0.00E+00 | 1.09E+02 | 5.22E+01 | 1.00 |
| 16 | Q1Q7J5 | DUF1858 domain-containing protein | 0.00E+00 | 2.55E+03 | 5.74E+02 | 1.00 |
| 17 | Q1Q6R5 | Flagellar protein FliL | 0.00E+00 | 2.39E+02 | 9.07E+01 | 1.00 |
| 18 | Q1Q106 | Flavodoxin-like domain-containing protein | 0.00E+00 | 3.31E+01 | 3.01E+01 | 1.00 |
| 19 | Q1PZ82 | Glyceraldehyde-3-phosphate dehydrogenase | 1.73E+01 | 0.00E+00 | 2.72E+01 | 1.00 |
| 20 | Q1PXF9 | Glycogen branching enzyme, GH-57-type, archaeal | 0.00E+00 | 2.25E+01 | 1.85E+01 | 1.00 |
| 21 | Q1Q012 | HEAT repeat domain-containing protein | 0.00E+00 | 9.48E+01 | 7.33E+01 | 1.00 |
| 22 | Q1Q5K4 | histidine kinase | 0.00E+00 | 5.33E+01 | 5.17E+01 | 1.00 |
| 23 | Q1PWX7 | Imidazole glycerol phosphate synthase subunit HisH | 0.00E+00 | 1.74E+02 | 7.74E+01 | 1.00 |
| 24 | Q1PXU6 | Insulinase family protein | 0.00E+00 | 6.32E+02 | 5.60E+02 | 1.00 |
| 25 | Q1Q201 | Integration host factor subunit beta | 0.00E+00 | 8.33E+01 | 7.87E+01 | 1.00 |
| 26 | A0A2C9CC85 | Large ribosomal subunit protein bL19 | 0.00E+00 | 7.47E+01 | 4.06E+01 | 1.00 |
| 27 | Q1PVS7 | Metallo-beta-lactamase family protein, RNA-specific | 0.00E+00 | 3.90E+01 | 2.78E+01 | 1.00 |
| 28 | Q1Q1J9 | Methylthioribose-1-phosphate isomerase | 0.00E+00 | 5.32E+02 | 3.89E+02 | 1.00 |
| 29 | Q1PX75 | Methyltransferase | 0.00E+00 | 1.04E+03 | 1.51E+02 | 1.00 |
| 30 | Q1Q2I9 | MOSC domain-containing protein | 9.00E+01 | 3.21E+02 | 0.00E+00 | 1.00 |
| 31 | Q1Q242 | N-acetylglutamate synthase | 0.00E+00 | 6.99E+01 | 4.69E+01 | 1.00 |
| 32 | Q1Q0K4 | NIF system FeS cluster assembly NifU C-terminal domain-containing protein | 0.00E+00 | 7.83E+01 | 4.64E+01 | 1.00 |
| 33 | Q1Q7F3 | peptidoglycan lytic exotransglycosylase | 0.00E+00 | 2.73E+02 | 2.38E+02 | 1.00 |
| 34 | Q1PY30 | Phosphoribosylamine--glycine ligase | 0.00E+00 | 1.99E+02 | 7.48E+01 | 1.00 |
| 35 | Q1PYD1 | PilZ domain-containing protein | 0.00E+00 | 1.10E+02 | 7.75E+01 | 1.00 |
| 36 | Q1Q0Q0 | Putative acriflavine resistance protein F | 0.00E+00 | 6.01E+03 | 5.26E+02 | 1.00 |
| 37 | Q1Q7J1 | Putative hydroxylamine oxidoreductase hao | 0.00E+00 | 4.46E+01 | 4.17E+01 | 1.00 |
| 38 | Q1PXM2 | Putative N-acetyl-alpha-D-glucosaminyl L-malate deacetylase 1 | 0.00E+00 | 3.39E+02 | 6.88E+01 | 1.00 |
| 39 | Q1Q7A2 | Putative serine/threonine specific protein phosphatase 2 | 0.00E+00 | 2.35E+02 | 1.03E+02 | 1.00 |
| 40 | Q1PZS9 | Pyruvoyl-dependent arginine decarboxylase AaxB | 0.00E+00 | 1.76E+02 | 1.75E+02 | 1.00 |
| 41 | Q1PZ79 | Ribulose-phosphate 3-epimerase | 0.00E+00 | 1.78E+02 | 1.70E+02 | 1.00 |
| 42 | Q1PVM6 | Strongly similar to peroxiredoxin (Thioredoxin peroxidase) | 0.00E+00 | 4.43E+02 | 1.87E+02 | 1.00 |
| 43 | Q1Q4I1 | Type II toxin-antitoxin system RelE/ParE family toxin | 0.00E+00 | 2.02E+02 | 1.48E+02 | 1.00 |
| 44 | Q1PZV2 | Tyrosine--tRNA ligase | 0.00E+00 | 4.85E+01 | 1.35E+01 | 1.00 |
| 45 | A0A2C9CCM4 | UDP-N-acetylglucosamine 1-carboxyvinyltransferase | 0.00E+00 | 6.36E+01 | 4.11E+01 | 1.00 |
| 46 | A0A2C9CGT6 | Uncharacterized protein | 0.00E+00 | 7.85E+01 | 4.85E+01 | 1.00 |
| 47 | Q1PW80 | Uncharacterized protein | 0.00E+00 | 1.91E+02 | 7.38E+01 | 1.00 |
| 48 | Q1Q6L5 | Uncharacterized protein | 0.00E+00 | 7.10E+01 | 5.69E+01 | 1.00 |
| 49 | Q1PV39 | Secondary thiamine-phosphate synthase enzyme | 0.00E+00 | 5.55E+02 | 8.76E+01 | 1.00 |

| Protein number | Accession | Description | Relative protein abundance per fraction FT | sample A | Low-resolution fraction 6 | Pearson correlation score between specific activity and relative protein abundance |
|---|---|---|---|---|---|---|
| 50 | A0A2C9CAF9 | 2-C-methyl-D-erythritol 4-phosphate cytidylyltransferase | 0.00E+00 | 9.38E+01 | 3.11E+01 | 1.00 |
| 51 | Q1PZP7 | Uncharacterized protein | 0.00E+00 | 3.40E+02 | 9.31E+01 | 1.00 |
| 52 | Q1PWY2 | 4-hydroxy-tetrahydrodipicolinate reductase | 4.05E+01 | 1.76E+02 | 6.27E+01 | 1.00 |
| 53 | Q1PY54 | Methyltransferase domain-containing protein | 1.32E+02 | 5.55E+02 | 2.21E+02 | 1.00 |
| 54 | Q1PXH9 | ADP-dependent (S)-NAD(P)H-hydrate dehydratase | 2.56E+01 | 7.64E+01 | 2.97E+01 | 1.00 |
| 55 | Q1PZK4 | Chaperonin GroEL | 3.79E+01 | 1.80E+03 | 1.50E+02 | 1.00 |
| 56 | Q1PX08 | Regulatory protein P-II for glutamine synthetase | 3.02E+01 | 4.75E+01 | 3.05E+01 | 0.99 |
| 57 | Q1PZD9 | Blue copper protein | 2.48E+01 | 3.76E+02 | 1.75E+01 | 0.99 |
| 58 | Q1Q0Y3 | Glucose-6-phosphate isomerase | 3.71E+01 | 6.01E+01 | 3.60E+01 | 0.98 |
| 59 | Q1Q1E8 | Chaperone protein ClpB | 5.15E+01 | 2.33E+02 | 1.20E+02 | 0.97 |
| 60 | Q1PV08 | Hemerythrin-like domain-containing protein | 3.54E+01 | 5.10E+02 | 2.31E+02 | 0.96 |
| 61 | Q1PX62 | DUF4438 domain-containing protein | 4.89E+01 | 6.77E+01 | 5.73E+01 | 0.95 |
| 62 | Q1PZC1 | Addiction module component | 7.47E+01 | 3.17E+02 | 2.15E+01 | 0.95 |
| 63 | Q1Q2L9 | Aspartate-semialdehyde dehydrogenase | 2.96E+01 | 6.91E+01 | 1.70E+01 | 0.93 |
| 64 | Q1Q2J0 | Molybdenum cofactor biosynthesis protein B | 3.40E+02 | 3.75E+02 | 3.58E+02 | 0.92 |
| 65 | Q1PXR8 | Serine-pyruvate aminotransferase | 7.08E+02 | 1.45E+03 | 4.31E+02 | 0.91 |
| 66 | Q1Q3C3 | Putative NADH dehydrogenase/NAD(P)H nitroreductase | 2.38E+02 | 5.66E+02 | 1.08E+02 | 0.91 |
| 67 | Q1Q648 | Bifunctional protein FolD | 2.69E+02 | 6.05E+02 | 1.35E+02 | 0.91 |
| 68 | Q1Q087 | Pyrroline-5-carboxylate reductase | 2.75E+01 | 1.75E+02 | 1.14E+02 | 0.89 |
| 69 | Q1Q646 | Probable cytosol aminopeptidase | 3.94E+01 | 4.70E+01 | 4.42E+01 | 0.87 |
| 70 | Q1Q131 | Large ribosomal subunit protein uL1 | 2.07E+02 | 3.94E+02 | 9.42E+01 | 0.86 |
| 71 | Q1Q6K3 | Strongly similar to exoribonuclease YhaM | 7.52E+01 | 1.10E+02 | 5.36E+01 | 0.86 |
| 72 | Q1PVK4 | Similar to phosphoglucomutase | 3.06E+01 | 8.38E+01 | 6.90E+01 | 0.81 |
| 73 | Q1Q2J4 | Elongation factor Ts | 8.74E+02 | 2.74E+03 | 2.27E+03 | 0.80 |
| 74 | Q1Q156 | Adenylate kinase | 1.79E+01 | 1.65E+03 | 1.29E+03 | 0.78 |
| 75 | A0A2C9CKJ6 | Exported protein | 1.72E+02 | 3.43E+03 | 2.72E+03 | 0.77 |
| 76 | Q1Q353 | DNA polymerase beta | 8.24E+01 | 1.47E+02 | 1.36E+02 | 0.74 |
| 77 | Q1PXP1 | Acetoacetate metabolism regulatory protein AtoC | 1.64E+01 | 4.12E+01 | 4.29E+01 | 0.58 |
| 78 | Q1PV34 | 2',3'-cyclic-nucleotide 2'-phosphodiesterase | 8.20E+01 | 5.74E+02 | 6.16E+02 | 0.57 |
| 79 | Q1PYR9 | Putative fructose-biphosphate aldolase | 2.81E+02 | 3.04E+02 | 1.13E+02 | 0.46 |
| 80 | Q1PXW4 | N-acetyl-gamma-glutamyl-phosphate reductase | 1.55E+02 | 1.69E+02 | 4.37E+01 | 0.46 |
| 81 | Q1Q129 | Large ribosomal subunit protein bL12 | 4.21E+01 | 2.07E+02 | 2.93E+02 | 0.33 |
| 82 | Q1PVQ5 | Putative superoxide reductase | 3.05E+01 | 9.18E+01 | 1.29E+02 | 0.29 |
| 83 | Q1Q0V3 | Aconitate hydratase | 2.56E+02 | 2.29E+02 | 9.54E+01 | 0.21 |
| **84** | **Q1Q4F5** | **Nitrite reductase; strongly similar to NirS** | **3.62E+01** | **7.17E+01** | **1.01E+02** | **0.20** |
| 85 | Q1Q3U7 | LL-diaminopimelate aminotransferase | 7.87E+02 | 5.93E+02 | 5.99E+01 | 0.11 |
| 86 | Q1Q652 | NAD-dependent epimerase/dehydratase domain-containing protein | 1.52E+02 | 1.19E+02 | 4.06E+01 | 0.07 |
| 87 | Q1PVQ3 | Putative small heat shock protein | 1.19E+02 | 4.42E+02 | 9.17E+02 | 0.04 |
| 88 | Q1Q0L1 | Cobalt-precorrin-4 C(11)-methyltransferase | 3.19E+01 | 3.80E+01 | 4.98E+01 | -0.03 |
| 89 | Q1PYI8 | Alanine--tRNA ligase | 6.63E+01 | 1.35E+02 | 3.78E+02 | -0.16 |
| 90 | Q1PY42 | Co-chaperonin GroES | 1.48E+02 | 1.21E+03 | 9.27E+03 | -0.26 |
| 91 | Q1PY25 | Thioredoxin | 1.71E+02 | 1.08E+02 | 6.74E+01 | -0.27 |
| 92 | Q1PW67 | Peptidylprolyl isomerase | 1.42E+01 | 1.06E+02 | 9.48E+02 | -0.28 |
| 93 | Q1Q5R5 | Aspartokinase | 2.80E+02 | 3.07E+02 | 7.27E+02 | -0.31 |
| 94 | Q1PUK7 | Formate--tetrahydrofolate ligase | 1.68E+03 | 9.97E+02 | 6.47E+02 | -0.33 |
| 95 | Q1PYA3 | Putative beta-lactamase-inhibitor-like PepSY-like domain-containing protein | 6.17E+02 | 2.04E+02 | 6.47E+01 | -0.42 |
| 96 | Q1Q3B4 | HIT domain-containing protein | 5.69E+01 | 2.13E+01 | 1.05E+01 | -0.44 |
| 97 | Q1Q637 | Exported protein | 1.84E+02 | 6.90E+01 | 3.50E+01 | -0.44 |
| 98 | Q1Q0T4 | Hydrazine synthase subunit C | 6.52E+02 | 2.11E+02 | 4.70E+03 | -0.44 |
| 99 | Q1Q302 | UDP-glucose 6-dehydrogenase | 4.48E+01 | 2.04E+01 | 1.36E+01 | -0.45 |
| 100 | Q1PW28 | Uncharacterized protein | 7.56E+03 | 1.82E+03 | 3.93E+02 | -0.47 |
| 101 | Q1Q177 | Pyruvate, phosphate dikinase | 9.35E+01 | 7.29E+01 | 2.26E+02 | -0.48 |
| 102 | Q1Q1N5 | Acetate-CoA ligase [ADP-forming] I | 3.41E+02 | 7.18E+01 | 2.53E+01 | -0.51 |
| 103 | Q1PZB4 | CSD domain-containing protein | 2.31E+02 | 9.55E+01 | 7.46E+01 | -0.53 |
| 104 | Q1PUT0 | glucose-1-phosphate adenylyltransferase | 9.80E+01 | 3.47E+01 | 2.54E+01 | -0.53 |

| Protein number | Accession | Description | Relative protein abundance per fraction FT | sample A | Low-resolution fraction 6 | Pearson correlation score between specific activity and relative protein abundance |
|---|---|---|---|---|---|---|
| 105 | Q1Q3W5 | Glutamate synthase (NADPH) large chain | 5.82E+02 | 1.52E+02 | 9.52E+01 | -0.54 |
| 106 | A0A2C9CJF3 | DUF1318 domain-containing protein | 7.59E+02 | 1.21E+02 | 6.04E+01 | -0.56 |
| 107 | Q1PZE4 | Blue copper protein | 3.79E+03 | 4.08E+02 | 1.15E+02 | -0.57 |
| 108 | Q1PXI6 | Polymerase nucleotidyl transferase domain-containing protein | 2.08E+02 | 2.96E+01 | 1.56E+01 | -0.57 |
| 109 | A0A2C9CKP3 | Cobalt-precorrin-3b C17-methyltransferase | 6.69E+01 | 3.69E+01 | 3.53E+01 | -0.59 |
| 110 | Q1Q2J2 | Ribosome-recycling factor | 1.40E+03 | 1.13E+02 | 5.22E+01 | -0.59 |
| 111 | Q1Q117 | Phosphoribosylaminoimidazole-succinocarboxamide synthase | 1.03E+03 | 9.61E+01 | 5.65E+01 | -0.60 |
| 112 | Q1Q2R9 | Putative sorbitol dehydrogenase | 1.53E+03 | 3.30E+01 | 2.63E+01 | -0.62 |
| 113 | Q1PUK4 | S-adenosylmethionine synthase | 7.13E+03 | 3.23E+01 | 2.63E+01 | -0.63 |
| 114 | Q1Q0T9 | Hypothetical (Triheme) protein | 2.00E+04 | 6.86E+01 | 8.77E+01 | -0.63 |
| 115 | A0A2C9CE24 | Serine hydroxymethyltransferase | 2.75E+03 | 2.07E+01 | 2.52E+01 | -0.63 |
| 116 | Q1Q7I1 | Outer membrane protein (OmpH-like) | 3.41E+03 | 6.89E+01 | 7.91E+01 | -0.63 |
| 117 | Q1Q5X6 | Peptidyl-prolyl cis-trans isomerase ppiD | 6.78E+03 | 1.06E+02 | 1.38E+02 | -0.63 |
| 118 | Q1PZC8 | Putative septation protein SpoVG | 3.99E+02 | 8.34E+01 | 8.88E+01 | -0.64 |
| 119 | Q1PYU2 | GTPase | 6.07E+02 | 1.61E+01 | 2.87E+01 | -0.64 |
| 120 | Q1PY19 | tRNA-splicing ligase RtcB | 3.40E+02 | 5.84E+01 | 6.62E+01 | -0.64 |
| 121 | Q1PUP1 | Asl1-like glycosyl hydrolase catalytic domain-containing protein | 7.76E+02 | 4.34E+02 | 4.81E+02 | -0.72 |
| 122 | Q1PY86 | PepSY domain-containing protein | 1.06E+03 | 6.47E+01 | 2.43E+02 | -0.75 |
| 123 | Q1Q0Y1 | Cytochrome c551 peroxidase | 5.77E+01 | 1.69E+01 | 2.48E+01 | -0.76 |
| 124 | Q1Q3B7 | Exported protein | 2.09E+03 | 1.81E+02 | 6.39E+02 | -0.79 |
| 125 | Q1Q467 | LamG-like jellyroll fold domain-containing protein | 2.27E+02 | 1.41E+01 | 7.00E+01 | -0.80 |
| 126 | Q1Q664 | Dihydrolipoamide acetyltransferase component of pyruvate dehydrogenase complex | 1.54E+02 | 9.30E+01 | 1.12E+02 | -0.83 |
| 127 | Q1PZE3 | Carboxypeptidase regulatory-like domain-containing protein | 2.99E+02 | 1.72E+02 | 2.16E+02 | -0.86 |
| 128 | Q1Q4R9 | NYN domain-containing protein | 6.17E+01 | 2.61E+01 | 3.89E+01 | -0.86 |
| 129 | Q1Q4Z1 | Cytochrome c | 5.60E+02 | 2.78E+02 | 6.87E+02 | -0.90 |
| 130 | Q1PZD5 | Nitrite oxidoreductase subunit B | 4.71E+01 | 3.24E+01 | 4.57E+01 | -1.00 |
| 131 | Q1PZB1 | Aminomethyltransferase | 0.00E+00 | 1.93E+01 | 1.71E+02 | -1.00 |
| 132 | Q1Q3S4 | Cobalamin biosynthesis precorrin-8X methylmutase CobH/CbiC domain-containing protein | 0.00E+00 | 2.57E+02 | 2.65E+02 | -1.00 |
| 133 | Q1Q449 | hydroxymethylpyrimidine kinase | 1.49E+02 | 0.00E+00 | 8.94E+00 | -1.00 |
| 134 | Q1PVN8 | Putative molybdopterin oxidoreductase, molybdopterin-containing subunit | 1.83E+02 | 0.00E+00 | 2.12E+01 | -1.00 |
| 135 | Q1Q3U6 | Tryptophan--tRNA ligase | 0.00E+00 | 2.42E+02 | 1.52E+03 | -1.00 |
| 136 | Q1Q6D5 | Uncharacterized protein | 0.00E+00 | 3.55E+01 | 1.07E+02 | -1.00 |
| 137 | Q1Q4R8 | Bifunctional chorismate mutase/prephenate dehydratase | 0.00E+00 | 6.22E+01 | 1.75E+02 | -1.00 |
| 138 | Q1PZ66 | DUF1858 domain-containing protein | 0.00E+00 | 1.29E+02 | 1.47E+03 | -1.00 |
| 139 | Q1PVV7 | Similar to glutamate-1-semialdehyde 2,1-aminomutase | 0.00E+00 | 1.18E+02 | 5.44E+02 | -1.00 |
| 140 | Q1Q2R8 | 3-hydroxy-5-phosphonooxypentane-2,4-dione thiolase | 0.00E+00 | 1.05E+02 | 5.90E+02 | -1.00 |
| 141 | Q1Q122 | 3-isopropylmalate dehydratase large subunit | 0.00E+00 | 2.17E+01 | 2.99E+01 | -1.00 |
| 142 | Q1Q7P0 | 4Fe-4S ferredoxin | 0.00E+00 | 1.48E+02 | 2.20E+02 | -1.00 |
| 143 | Q1Q1D3 | 6-carboxy-5,6,7,8-tetrahydropterin synthase | 0.00E+00 | 7.72E+01 | 1.29E+02 | -1.00 |
| 144 | Q1Q2A1 | ABC transporter substrate-binding protein | 0.00E+00 | 1.09E+02 | 5.09E+02 | -1.00 |
| 145 | Q1PZV6 | Adenine phosphoribosyltransferase | 0.00E+00 | 2.06E+02 | 2.91E+02 | -1.00 |
| 146 | Q1Q338 | Adenylylsulfate reductase alpha-subunit | 0.00E+00 | 3.02E+02 | 1.03E+03 | -1.00 |
| 147 | Q1Q6F6 | Alanine racemase | 0.00E+00 | 1.18E+01 | 9.71E+01 | -1.00 |
| 148 | Q1Q1A1 | Aldehyde oxidase/xanthine dehydrogenase | 0.00E+00 | 4.15E+01 | 5.68E+01 | -1.00 |
| 149 | Q1PZS8 | Arginine biosynthesis bifunctional protein ArgJ | 0.00E+00 | 3.35E+03 | 3.54E+03 | -1.00 |
| 150 | Q1PV06 | Beta-lactamase hydrolase-like protein | 0.00E+00 | 1.41E+01 | 7.03E+02 | -1.00 |
| 151 | Q1Q3D6 | Carboxypeptidase regulatory-like domain-containing protein | 0.00E+00 | 2.38E+02 | 7.86E+02 | -1.00 |
| 152 | Q1Q1B0 | CBS domain-containing protein | 6.36E+01 | 3.89E+01 | 0.00E+00 | -1.00 |
| 153 | Q1PXW0 | Chaperonin GroEL | 0.00E+00 | 3.59E+01 | 1.20E+02 | -1.00 |
| 154 | Q1PZY2 | Chromosome partition protein Smc | 5.09E+01 | 0.00E+00 | 4.40E+01 | -1.00 |
| 155 | Q1Q5P8 | citramalate synthase | 0.00E+00 | 4.85E+01 | 9.91E+01 | -1.00 |
| 156 | Q1Q7P3 | Class I small soluble cyt c | 0.00E+00 | 7.09E+01 | 1.65E+03 | -1.00 |
| 157 | Q1PZK3 | Co-chaperonin GroES | 0.00E+00 | 2.26E+02 | 8.10E+02 | -1.00 |
| 158 | Q1Q2R6 | Cysteine synthase A, O-acetylserine sulfhydrolase A subunit | 0.00E+00 | 6.51E+01 | 8.47E+02 | -1.00 |
| 159 | Q1PZY5 | Cytochrome c551 peroxidase | 1.52E+02 | 1.04E+02 | 0.00E+00 | -1.00 |

| Protein number | Accession | Description | Relative protein abundance per fraction FT | sample A | Low-resolution fraction 6 | Pearson correlation score between specific activity and relative protein abundance |
|---|---|---|---|---|---|---|
| 160 | Q1Q787 | Cytochrome p450 hydroxylase MmcK-like protein | 0.00E+00 | 5.11E+01 | 7.61E+01 | -1.00 |
| 161 | Q1PZY1 | D-3-phosphoglycerate dehydrogenase | 0.00E+00 | 9.77E+00 | 1.99E+02 | -1.00 |
| 162 | Q1Q5N0 | Diaminopimelate decarboxylase | 0.00E+00 | 2.87E+02 | 4.11E+02 | -1.00 |
| 163 | Q1PX65 | DNA ligase | 0.00E+00 | 2.05E+01 | 1.25E+02 | -1.00 |
| 164 | Q1PYZ6 | DNA topoisomerase 1 | 0.00E+00 | 8.38E+00 | 2.46E+01 | -1.00 |
| 165 | Q1PUL9 | DUF1015 domain-containing protein | 0.00E+00 | 4.84E+01 | 5.76E+01 | -1.00 |
| 166 | Q1Q7G7 | DUF1611 domain-containing protein | 0.00E+00 | 1.15E+02 | 4.11E+02 | -1.00 |
| 167 | Q1PV11 | DUF2024 domain-containing protein | 3.96E+01 | 0.00E+00 | 1.49E+01 | -1.00 |
| 168 | Q1Q7J2 | DUF2249 domain-containing protein | 0.00E+00 | 4.25E+02 | 7.16E+02 | -1.00 |
| 169 | Q1Q598 | DUF262 domain-containing protein | 0.00E+00 | 4.09E+02 | 5.38E+02 | -1.00 |
| 170 | A0A2C9CI86 | DUF3024 domain-containing protein | 0.00E+00 | 2.10E+02 | 3.32E+02 | -1.00 |
| 171 | Q1Q1R5 | DUF4388 domain-containing protein | 0.00E+00 | 8.04E+01 | 9.38E+01 | -1.00 |
| 172 | Q1Q141 | Elongation factor G | 0.00E+00 | 1.81E+02 | 1.85E+02 | -1.00 |
| 173 | A0A2C9CCN1 | Fragment of pimeloyl-CoA synthetase (Part 2) | 2.98E+01 | 0.00E+00 | 1.08E+01 | -1.00 |
| 174 | Q1PYG4 | GDP-L-fucose synthase | 0.00E+00 | 1.33E+02 | 1.52E+02 | -1.00 |
| 175 | Q1PX01 | GMP synthase [glutamine-hydrolyzing] | 0.00E+00 | 1.33E+02 | 3.25E+02 | -1.00 |
| 176 | Q1Q060 | GTPase Obg | 0.00E+00 | 6.26E+01 | 6.46E+01 | -1.00 |
| 177 | Q1PY26 | Indole-3-glycerol phosphate synthase | 0.00E+00 | 5.84E+01 | 3.11E+02 | -1.00 |
| 178 | Q1PZD0 | Ion-translocating oxidoreductase complex subunit B | 0.00E+00 | 9.70E+01 | 4.16E+02 | -1.00 |
| 179 | Q1PYH4 | Malonyl CoA-acyl carrier protein transacylase | 0.00E+00 | 1.38E+01 | 1.02E+02 | -1.00 |
| 180 | Q1Q6F1 | Methyltransferase domain-containing protein | 0.00E+00 | 1.37E+02 | 2.38E+02 | -1.00 |
| 181 | Q1Q604 | N-acetylmuramic acid 6-phosphate etherase | 5.40E+03 | 3.65E+01 | 0.00E+00 | -1.00 |
| 182 | Q1Q4I3 | NAD(P)H-hydrate epimerase | 0.00E+00 | 3.30E+01 | 3.73E+02 | -1.00 |
| 183 | A0A2C9CMW4 | NAD-reducing hydrogenase HoxS subunit beta | 0.00E+00 | 4.22E+01 | 5.57E+01 | -1.00 |
| 184 | Q1PZI4 | Nucleoside diphosphate kinase | 0.00E+00 | 1.71E+02 | 7.44E+02 | -1.00 |
| 185 | Q1Q1G1 | Orotidine 5'-phosphate decarboxylase | 3.98E+02 | 2.82E+01 | 0.00E+00 | -1.00 |
| 186 | Q1PV60 | PEGA domain-containing protein | 0.00E+00 | 2.80E+02 | 5.37E+02 | -1.00 |
| 187 | Q1PY73 | pEK499-p136 HEPN domain-containing protein | 0.00E+00 | 3.63E+01 | 7.15E+02 | -1.00 |
| 188 | Q1PYD2 | Peptidyl-prolyl cis-trans isomerase | 4.62E+02 | 2.63E+00 | 0.00E+00 | -1.00 |
| 189 | Q1Q6P8 | Phosphoesterase | 0.00E+00 | 8.12E+01 | 1.71E+02 | -1.00 |
| 190 | Q1Q013 | phosphomannomutase | 4.44E+02 | 5.16E+01 | 0.00E+00 | -1.00 |
| 191 | Q1Q3G2 | Phosphoribosylglycinamide formyltransferase | 1.05E+03 | 0.00E+00 | 1.58E+02 | -1.00 |
| 192 | Q1Q6R0 | Prephenate dehydrogenase | 0.00E+00 | 5.25E+02 | 1.33E+03 | -1.00 |
| 193 | Q1Q355 | Putative adenylylsulfate reductase chain B | 0.00E+00 | 4.75E+02 | 7.33E+02 | -1.00 |
| 194 | Q1PVG3 | Putative catabolite gene activator (CAMP receptor protein) | 0.00E+00 | 8.10E+01 | 1.88E+02 | -1.00 |
| 195 | Q1Q2D9 | Putative periplasmic serine endoprotease DegP-like | 0.00E+00 | 6.05E+01 | 9.45E+01 | -1.00 |
| 196 | Q1PVM4 | Putative UDP-N-acetylglucosamine pyrophosphorylase | 0.00E+00 | 9.43E+00 | 1.93E+02 | -1.00 |
| 197 | Q1Q552 | Pyruvate:ferredoxin (Flavodoxin) oxidoreductase | 0.00E+00 | 1.09E+02 | 1.80E+02 | -1.00 |
| 198 | Q1Q045 | Radical SAM domain heme biosynthesis protein | 0.00E+00 | 3.41E+01 | 8.49E+01 | -1.00 |
| 199 | Q1PZE1 | Serine hydroxymethyltransferase | 1.30E+03 | 2.74E+02 | 0.00E+00 | -1.00 |
| 200 | Q1PXI1 | siroheme decarboxylase | 0.00E+00 | 1.81E+01 | 5.61E+01 | -1.00 |
| 201 | Q1Q728 | SpoVT-AbrB domain-containing protein | 0.00E+00 | 1.55E+02 | 1.77E+02 | -1.00 |
| 202 | Q1Q0W8 | Strongly similar to small heat shock protein | 0.00E+00 | 7.45E+02 | 1.29E+03 | -1.00 |
| 203 | Q1Q0I7 | Sulfur carrier protein ThiS adenylyltransferase | 0.00E+00 | 1.83E+01 | 2.84E+01 | -1.00 |
| 204 | Q1Q1K9 | Threonine synthase | 1.89E+03 | 6.65E+01 | 0.00E+00 | -1.00 |
| 205 | Q1PY20 | Thymidylate kinase | 0.00E+00 | 4.59E+01 | 1.28E+02 | -1.00 |
| 206 | Q1PX64 | TldD protein, part of proposed TldE/TldD proteolytic complex | 0.00E+00 | 1.44E+01 | 3.89E+02 | -1.00 |
| 207 | Q1Q775 | Transaldolase | 0.00E+00 | 1.68E+02 | 1.47E+03 | -1.00 |
| 208 | A0A6G7GTJ9 | Transcription factor CBF/NF-Y/archaeal histone domain-containing protein | 0.00E+00 | 5.20E+02 | 2.52E+03 | -1.00 |
| 209 | Q1Q088 | transketolase | 0.00E+00 | 3.03E+01 | 3.07E+02 | -1.00 |
| 210 | Q1PX46 | Tryptophan synthase alpha chain | 0.00E+00 | 1.86E+02 | 1.93E+03 | -1.00 |
| 211 | Q1PX47 | Tryptophan synthase beta chain | 0.00E+00 | 4.26E+02 | 9.24E+02 | -1.00 |
| 212 | Q1Q6I9 | Uncharacterized protein | 0.00E+00 | 2.10E+02 | 6.85E+02 | -1.00 |
| 213 | Q1PXU5 | Uncharacterized protein | 0.00E+00 | 2.25E+02 | 2.88E+02 | -1.00 |
| 214 | Q1Q2E9 | Conserved hypothetical iron sulfur protein | 0.00E+00 | 8.39E+01 | 3.29E+02 | -1.00 |

| Protein number | Accession | Description | Relative protein abundance per fraction FT | sample A | Low-resolution fraction 6 | Pearson correlation score between specific activity and relative protein abundance |
|---|---|---|---|---|---|---|
| 215 | Q1PXU8 | Conserved hypothetical thioredoxin protein | 1.00E+02 | 0.00E+00 | 3.64E+01 | -1.00 |
| 216 | Q1Q7J4 | Cupin type-2 domain-containing protein | 0.00E+00 | 1.26E+02 | 2.25E+03 | -1.00 |
| 217 | Q1Q5M8 | Dissimilatory sulfite reductase (Desulfoviridin), alpha and beta subunits | 1.79E+04 | 0.00E+00 | 8.43E+01 | -1.00 |
| 218 | Q1PX19 | DJ-1/PfpI domain-containing protein | 5.07E+02 | 0.00E+00 | 7.33E+00 | -1.00 |
| 219 | Q1PUZ2 | Glucose-6-phosphate 1-dehydrogenase | 2.12E+01 | 0.00E+00 | 9.54E+00 | -1.00 |
| 220 | Q1Q0T2 | Hydrazine synthase subunit A | 0.00E+00 | 5.14E+01 | 7.65E+03 | -1.00 |
| 221 | Q1PW61 | Imidazole glycerol phosphate synthase subunit HisF | 0.00E+00 | 2.96E+02 | 4.15E+02 | -1.00 |
| 222 | Q1PW68 | Ketol-acid reductoisomerase (NADP(+)) | 1.30E+04 | 4.50E+00 | 0.00E+00 | -1.00 |
| 223 | Q1Q1W6 | LamG-like jellyroll fold domain-containing protein | 0.00E+00 | 7.73E+01 | 3.04E+02 | -1.00 |
| 224 | Q1Q7J3 | Mannose-6-phosphate isomerase | 0.00E+00 | 2.10E+01 | 2.66E+02 | -1.00 |
| 225 | Q1PZD8 | Nitrite oxidoreductase subunit A | 1.03E+02 | 0.00E+00 | 4.21E+01 | -1.00 |
| 226 | Q1PVV8 | Similar to methionyl-tRNA formyltransferase | 0.00E+00 | 1.44E+03 | 5.05E+03 | -1.00 |
| 227 | Q1PVQ2 | Similar to small heat shock protein | 0.00E+00 | 1.34E+02 | 5.94E+02 | -1.00 |
| 228 | Q1PXG9 | Small heat shock protein-like protein | 0.00E+00 | 2.33E+02 | 1.20E+03 | -1.00 |
| 229 | Q1Q4D3 | TldD/PmbA family protein | 0.00E+00 | 5.86E+01 | 3.30E+02 | -1.00 |
| 230 | Q1Q4X7 | Uncharacterized protein | 0.00E+00 | 3.45E+01 | 2.33E+02 | -1.00 |
| 231 | Q1Q2M4 | Uncharacterized protein | 0.00E+00 | 1.34E+02 | 1.78E+02 | -1.00 |
| 232 | Q1PY07 | Uncharacterized protein | 0.00E+00 | 1.42E+01 | 1.71E+01 | -1.00 |
| 233 | Q1PW64 | Zinc ribbon domain protein | 0.00E+00 | 6.52E+01 | 3.82E+02 | -1.00 |

**Supplementary table 5** - Correlation between the relative abundance of identified proteins and specific nitrite reductase activity in all fractions obtained with mixed-mode column chromatography. Pearson correlation score was calculated between the specific activity measured per fraction and the relative protein abundance measured per fraction. The data is ordered based on relative protein abundance that correlated best with the specific activity. The relative abundance of HAOr correlated with specific nitrite reductase activity, while NirS does not. Other proteins that showed a positive correlation are not identified as nitrite reductases. Accession numbers refer to the *K. stuttgartiensis* protein sequence database in Uniprot (entry KSMBR1).

| Protein number | Accession | Description | Relative protein abundance per fraction | | | | Pearson correlation score |
|---|---|---|---|---|---|---|---|
| | | | UV-peak 1 | UV-peak 2 | UV-peak 3 | UV-peak 4 | between specific activity and relative protein abundance |
| 1 | A0A2C9CHX1 | strongly similar to 3naS ribosomal protein S1na | 4.75E+04 | 0.00E+00 | 0.00E+00 | 5.15E+05 | 1.00 |
| 2 | Q1Q132 | strongly similar to 5naS ribosomal protein L11 | 1.84E+04 | 0.00E+00 | 0.00E+00 | 2.45E+05 | 1.00 |
| 3 | Q1PY08 | strongly similar to 5naS ribosomal protein L11 | 2.78E+04 | 0.00E+00 | 0.00E+00 | 3.70E+05 | 1.00 |
| 4 | A0A6G7GP74 | Putative cellulose synthase, cyclic-di-GMP-binding regulatory subunit | 7.21E+05 | 0.00E+00 | 0.00E+00 | 9.99E+06 | 1.00 |
| 5 | A0A2C9CFG9 | Helicase ATP-binding domain-containing protein | 3.70E+04 | 0.00E+00 | 0.00E+00 | 5.91E+05 | 1.00 |
| 6 | A0A2C9CJB5 | Hydrazine dehydrogenase | 3.90E+04 | 0.00E+00 | 0.00E+00 | 3.29E+05 | 1.00 |
| 7 | A0A2C9CJU8 | Amine oxidase domain-containing protein | 6.74E+03 | 0.00E+00 | 0.00E+00 | 1.28E+05 | 1.00 |
| 8 | A0A2C9CJI9 | site-specific DNA-methyltransferase (adenine-specific) | 3.83E+04 | 0.00E+00 | 0.00E+00 | 1.82E+06 | 1.00 |
| 9 | Q1Q5F8 | strongly similar to bacterioferritin | 9.66E+05 | 0.00E+00 | 0.00E+00 | 1.24E+08 | 1.00 |
| 10 | Q1Q7J1 | similar to hydroxylamine oxidoreductase hao | 1.23E+05 | 2.41E+05 | 0.00E+00 | 1.80E+07 | 1.00 |
| 11 | Q1Q5E4 | hypothetical proteins | 0.00E+00 | 0.00E+00 | 0.00E+00 | 1.04E+05 | 1.00 |
| 12 | Q1PZT1 | similar to ribosomal protein S21 | 0.00E+00 | 0.00E+00 | 0.00E+00 | 5.97E+04 | 1.00 |
| 13 | A0A2C9CIR3 | Nucleoside diphosphate kinase | 0.00E+00 | 0.00E+00 | 0.00E+00 | 3.36E+05 | 1.00 |
| 14 | Q1PZY9 | similar to molybdenum cofactor biosynthesis protein | 0.00E+00 | 0.00E+00 | 0.00E+00 | 1.15E+05 | 1.00 |
| 15 | A0A2C9CGI1 | strongly similar to peroxiredoxin (thioredoxin peroxidase) | 0.00E+00 | 0.00E+00 | 0.00E+00 | 8.48E+04 | 1.00 |
| 16 | Q1Q0K4 | hypothetical protein | 0.00E+00 | 0.00E+00 | 0.00E+00 | 7.32E+05 | 1.00 |
| 17 | Q1Q5P1 | similar to naD(P) oxidoreductase, FAD-containing subunit | 0.00E+00 | 0.00E+00 | 0.00E+00 | 2.47E+05 | 1.00 |
| 19 | Q1PXG4 | hypothetical protein | 0.00E+00 | 0.00E+00 | 0.00E+00 | 1.75E+05 | 1.00 |
| 20 | Q1Q1U2 | Methyltransferase type 11 domain-containing protein | 1.44E+04 | 0.00E+00 | 0.00E+00 | 7.15E+04 | 0.99 |
| 21 | Q1PY40 | TIGR04076 family protein | 0.00E+00 | 2.39E+04 | 0.00E+00 | 3.21E+05 | 0.99 |
| 22 | Q1Q4R9 | hypothetical protein | 8.45E+04 | 0.00E+00 | 0.00E+00 | 3.88E+05 | 0.99 |
| **23** | **Q1PVE0** | **similar to hydroxylamine oxidoreductase, HAOr** | **1.51E+04** | **0.00E+00** | **0.00E+00** | **6.86E+04** | **0.99** |
| 24 | A0A2C9CFG1 | strongly similar to 6na kDa chaperonin (GroEL protein) | 4.51E+05 | 0.00E+00 | 0.00E+00 | 1.97E+06 | 0.99 |
| 25 | Q1PYJ8 | hypothetical protein | 3.51E+05 | 0.00E+00 | 5.81E+06 | 6.67E+07 | 0.99 |
| 26 | Q1PVE1 | c-type diheme-containing protein, redox partner of kustcna458 | 2.31E+04 | 0.00E+00 | 0.00E+00 | 8.69E+04 | 0.98 |
| 27 | Q1Q2J0 | similar to molybdenum cofactor biosynthesis protein B | 5.04E+03 | 0.00E+00 | 2.35E+05 | 1.85E+06 | 0.98 |
| 28 | Q1Q1C8 | hypothetical protein | 3.08E+04 | 0.00E+00 | 0.00E+00 | 1.09E+05 | 0.98 |
| 29 | A0A2C9CDA8 | Adenylyl-sulfate kinase | 0.00E+00 | 0.00E+00 | 6.35E+05 | 4.36E+06 | 0.97 |
| 30 | Q1Q3W5 | hypothetical protein | 3.46E+05 | 6.30E+04 | 0.00E+00 | 7.92E+05 | 0.94 |
| 31 | A0A2C9CF54 | hypothetical protein | 2.06E+05 | 0.00E+00 | 0.00E+00 | 4.53E+05 | 0.93 |
| 32 | Q1PXH4 | hypothetical protein | 4.93E+06 | 3.04E+05 | 0.00E+00 | 1.00E+07 | 0.92 |
| 33 | Q1PUY9 | UPF0234 protein KsCSTR_01580 | 2.17E+04 | 0.00E+00 | 0.00E+00 | 3.09E+04 | 0.81 |
| 34 | Q1PYP7 | rna binding protein | 1.07E+06 | 0.00E+00 | 0.00E+00 | 1.48E+06 | 0.80 |
| 35 | A0A2C9CGH9 | hypothetical protein | 3.90E+06 | 0.00E+00 | 0.00E+00 | 5.22E+06 | 0.78 |
| 36 | A0A2C9CJX5 | hypothetical protein | 6.89E+04 | 0.00E+00 | 0.00E+00 | 8.32E+04 | 0.74 |
| 37 | A0A2C9CB72 | Putative Alkaline phosphatase | 1.82E+06 | 0.00E+00 | 0.00E+00 | 2.14E+06 | 0.73 |
| 38 | Q1PZC1 | hypothetical protein | 3.01E+05 | 0.00E+00 | 0.00E+00 | 3.01E+05 | 0.64 |
| 39 | A0A2C9CGA7 | 4Fe-4S Mo/W bis-MGD-type domain-containing protein | 8.39E+04 | 1.00E+06 | 0.00E+00 | 9.90E+05 | 0.56 |
| 40 | Q1PXI6 | hypothetical protein | 0.00E+00 | 2.08E+06 | 0.00E+00 | 1.99E+06 | 0.54 |
| 41 | Q1Q0V3 | similar to aconitase 1 (aconitate hydratase 1; citrate hydro-lyase 1) | 6.50E+05 | 7.18E+05 | 2.29E+07 | 1.92E+07 | 0.40 |
| 42 | A0A2C9CJF3 | hypothetical protein | 1.44E+05 | 3.02E+06 | 0.00E+00 | 2.22E+06 | 0.38 |
| 43 | A0A2C9CF13 | strongly similar to 6na kDa chaperonin (groEL protein) | 2.92E+06 | 0.00E+00 | 0.00E+00 | 1.74E+06 | 0.35 |
| 44 | Q1Q2D4 | Phosphate-binding protein | 1.93E+06 | 0.00E+00 | 0.00E+00 | 1.07E+06 | 0.31 |
| 46 | Q1Q131 | strongly similar to 5naS ribosomal protein L1 | 1.53E+06 | 2.28E+05 | 0.00E+00 | 7.76E+05 | 0.23 |
| 47 | A0A2C9CDC4 | dihydropteroate synthase | 2.13E+05 | 1.41E+05 | 0.00E+00 | 1.43E+05 | 0.22 |
| 48 | Q1Q782 | Hypervirulence associated protein TUDOR domain-containing protein | 1.49E+06 | 5.87E+05 | 0.00E+00 | 8.21E+05 | 0.20 |
| 49 | Q1Q5X6 | hypothetical protein | 1.50E+07 | 2.43E+07 | 4.50E+08 | 2.56E+08 | 0.15 |
| 50 | Q1PVQ3 | similar to small heat shock protein | 5.82E+05 | 2.83E+05 | 0.00E+00 | 3.15E+05 | 0.15 |
| 51 | Q1PZP6 | probable nicotinate-nucleotide adenylyltransferase | 3.93E+05 | 1.11E+06 | 0.00E+00 | 5.34E+05 | 0.06 |

| Protein number | Accession | Description | Relative protein abundance per fraction UV-peak 1 | UV-peak 2 | UV-peak 3 | UV-peak 4 | Pearson correlation score between specific activity and relative protein abundance |
|---|---|---|---|---|---|---|---|
| 52 | Q1Q0Y1 | strongly similar to cytochrome c551 peroxidase | 7.60E+06 | 3.15E+05 | 0.00E+00 | 2.28E+06 | 0.03 |
| 53 | Q1PXP1 | Acetoacetate metabolism regulatory protein AtoC | 7.15E+05 | 0.00E+00 | 0.00E+00 | 2.01E+05 | 0.03 |
| 54 | A0A2C9CJS6 | Vitamin B12-dependent ribonucleotide reductase | 2.47E+06 | 1.23E+05 | 0.00E+00 | 6.81E+05 | 0.01 |
| 55 | Q1PXV3 | Fido domain-containing protein | 7.72E+04 | 1.98E+06 | 0.00E+00 | 7.03E+05 | 0.00 |
| 56 | Q1PUV5 | Methyltransferase domain-containing protein | 1.19E+06 | 0.00E+00 | 8.92E+05 | 6.51E+05 | -0.01 |
| 57 | A0A2C9CHM2 | hydrazine synthase subunit C | 4.25E+05 | 0.00E+00 | 0.00E+00 | 9.13E+04 | -0.04 |
| 59 | A0A2C9CAJ2 | Rhamnogalacturonan lyase domain-containing protein | 1.69E+07 | 2.47E+06 | 0.00E+00 | 4.33E+06 | -0.05 |
| 60 | A0A2C9CG37 | hypothetical protein | 4.11E+07 | 1.16E+06 | 0.00E+00 | 7.88E+06 | -0.08 |
| 61 | A0A2C9CKJ6 | hypothetical protein | 5.63E+05 | 3.25E+05 | 0.00E+00 | 2.17E+05 | -0.08 |
| 62 | Q1Q4B6 | hypothetical protein | 8.41E+05 | 0.00E+00 | 0.00E+00 | 1.29E+05 | -0.10 |
| 63 | Q1PZD8 | nitrite oxidoreductase subunit A | 7.25E+06 | 4.38E+05 | 0.00E+00 | 1.13E+06 | -0.12 |
| 64 | A0A2C9CJW9 | Macro domain-containing protein | 5.68E+06 | 0.00E+00 | 0.00E+00 | 7.36E+05 | -0.13 |
| 65 | A0A2C9CHN2 | hydrazine synthase subunit A | 1.41E+06 | 0.00E+00 | 0.00E+00 | 1.80E+05 | -0.13 |
| 66 | A0A2C9CB53 | Periplasmic zinc binding protein-like protein | 8.73E+04 | 5.19E+06 | 0.00E+00 | 1.06E+06 | -0.15 |
| 67 | Q1Q6B5 | Cell division protein FtsI (Peptidoglycan synthetase) | 1.30E+06 | 0.00E+00 | 0.00E+00 | 1.22E+05 | -0.16 |
| 68 | Q1Q2D9 | similar to heat shock protease DegP/HtrA | 1.99E+06 | 0.00E+00 | 0.00E+00 | 1.69E+05 | -0.17 |
| 69 | Q1PZD5 | nitrite oxidoreductase subunit B | 3.19E+06 | 4.82E+05 | 0.00E+00 | 4.63E+05 | -0.17 |
| 70 | A0A2C9CAF9 | 2-C-methyl-D-erythritol 4-phosphate cytidylyltransferase | 8.44E+06 | 0.00E+00 | 0.00E+00 | 6.38E+05 | -0.18 |
| 71 | A0A2C9CDZ7 | S-layer protein | 9.01E+05 | 0.00E+00 | 0.00E+00 | 6.59E+04 | -0.18 |
| 72 | A0A2C9CCT8 | Aldehyde oxidase/xanthine dehydrogenase | 1.73E+07 | 0.00E+00 | 0.00E+00 | 1.25E+06 | -0.18 |
| 73 | Q1Q2I9 | MOSC domain-containing protein | 1.06E+06 | 0.00E+00 | 0.00E+00 | 7.29E+04 | -0.19 |
| 74 | Q1PZD4 | nitrite oxidoreductase subunit C | 1.61E+06 | 0.00E+00 | 0.00E+00 | 1.09E+05 | -0.19 |
| 75 | A0A2C9CH14 | hydrazine synthase subunit B | 8.55E+05 | 0.00E+00 | 0.00E+00 | 5.63E+04 | -0.19 |
| 76 | A0A2C9CHG0 | hypothetical protein | 1.39E+05 | 4.21E+06 | 2.42E+05 | 8.44E+05 | -0.19 |
| 77 | Q1PXR8 | strongly similar to aspartate transaminase | 2.26E+07 | 3.73E+05 | 2.59E+06 | 2.86E+06 | -0.19 |
| **78** | **Q1Q4F5** | **strongly similar to cd1 nitrite reductase NirS** | **1.32E+06** | **1.90E+05** | **0.00E+00** | **1.64E+05** | **-0.19** |
| 79 | Q1Q353 | DNA polymerase beta | 1.62E+06 | 6.66E+06 | 0.00E+00 | 1.55E+06 | -0.20 |
| 80 | Q1Q4U2 | Putative methylmalonyl-CoA epimerase | 3.04E+06 | 0.00E+00 | 0.00E+00 | 1.78E+05 | -0.20 |
| 81 | A0A2C9CL22 | Strongly similar to rhodanese sulfur transferase and phage shock protein pspE | 2.24E+06 | 0.00E+00 | 0.00E+00 | 1.29E+05 | -0.20 |
| 82 | A0A2C9CCM4 | UDP-N-acetylglucosamine 1-carboxyvinyltransferase | 1.16E+05 | 0.00E+00 | 0.00E+00 | 6.49E+03 | -0.20 |
| 83 | Q1PYZ7 | hypothetical protein | 4.36E+05 | 0.00E+00 | 0.00E+00 | 2.41E+04 | -0.20 |
| 84 | Q1PZY5 | strongly similar to cytochrome c peroxidase | 9.66E+07 | 4.10E+06 | 0.00E+00 | 6.93E+06 | -0.20 |
| 85 | A0A2C9CJI3 | hypothetical protein | 1.84E+06 | 0.00E+00 | 0.00E+00 | 8.82E+04 | -0.21 |
| 86 | A0A2C9CKU3 | precorrin-2 dehydrogenase | 4.00E+06 | 1.63E+04 | 0.00E+00 | 1.43E+05 | -0.22 |
| 87 | Q1Q3G3 | similar to matrilysin (metalloproteinase) | 8.18E+06 | 0.00E+00 | 0.00E+00 | 2.55E+05 | -0.22 |
| 88 | A0A2C9CEG0 | hypothetical protein/ part of the Nxr 'operon' | 1.01E+08 | 6.94E+06 | 1.57E+06 | 7.06E+06 | -0.22 |
| 89 | A0A6G7GLF5 | S-layer protein | 3.44E+06 | 0.00E+00 | 0.00E+00 | 8.39E+04 | -0.23 |
| 90 | Q1Q1X2 | Putative mannosyltransferase B | 8.84E+05 | 0.00E+00 | 0.00E+00 | 2.15E+04 | -0.23 |
| 91 | Q1PY41 | stronlgy similar to 6na kDa chaperonin (GroEL protein) | 9.49E+05 | 9.60E+04 | 3.41E+06 | 9.85E+05 | -0.23 |
| 92 | Q1PVK4 | similar to phosphoglucomutase | 1.58E+07 | 8.58E+04 | 0.00E+00 | 3.76E+05 | -0.23 |
| 93 | Q1Q1B0 | hypothetical protein | 7.25E+06 | 0.00E+00 | 0.00E+00 | 1.46E+05 | -0.23 |
| 94 | Q1Q2J4 | strongly similar to elongation factor Ts (EF-Ts) | 3.04E+07 | 2.62E+05 | 0.00E+00 | 6.97E+05 | -0.23 |
| 95 | Q1Q3G2 | Phosphoribosylglycinamide formyltransferase | 5.25E+07 | 5.54E+05 | 0.00E+00 | 1.24E+06 | -0.23 |
| 96 | Q1PYD2 | strongly similar to peptidylprolylisomerase | 1.48E+08 | 1.30E+06 | 0.00E+00 | 3.31E+06 | -0.23 |
| 97 | A0A2C9CAG3 | HIT domain-containing protein | 4.41E+06 | 1.51E+04 | 0.00E+00 | 8.10E+04 | -0.24 |
| 98 | A0A2C9CC36 | N-(5'-phosphoribosyl)anthranilate isomerase | 3.13E+06 | 0.00E+00 | 0.00E+00 | 5.14E+04 | -0.24 |
| 99 | A0A2C9CDY7 | hypothetical protein | 1.92E+06 | 0.00E+00 | 0.00E+00 | 2.41E+04 | -0.24 |
| 100 | A0A2C9CIZ7 | hypothetical protein | 4.47E+08 | 2.60E+06 | 0.00E+00 | 6.67E+06 | -0.24 |
| 101 | Q1Q4J7 | strongly similar to UDP-glucuronate 5'-epimerase | 0.00E+00 | 1.44E+06 | 0.00E+00 | 1.57E+05 | -0.24 |
| 102 | A0A2C9CET7 | similar to high affinity sulfate transporter (plant) | 4.75E+06 | 0.00E+00 | 0.00E+00 | 5.32E+04 | -0.24 |
| 103 | Q1Q648 | strongly similar to bifunctional methylene-tetrahydrofolate dehydrogenase | 1.36E+08 | 4.49E+06 | 6.20E+05 | 3.75E+06 | -0.24 |
| 104 | Q1Q3U7 | similar to N-succinyldiaminopimelate aminotransferase | 3.42E+08 | 3.63E+06 | 7.74E+03 | 4.88E+06 | -0.24 |
| 105 | Q1PUK7 | strongly similar to formyltetrahydrofolate synthetase | 2.58E+08 | 1.69E+07 | 1.17E+07 | 1.63E+07 | -0.24 |
| 106 | Q1Q515 | DNA-binding protein | 1.35E+07 | 3.33E+05 | 0.00E+00 | 2.70E+05 | -0.24 |
| 107 | Q1Q1G1 | Orotidine 5'-phosphate decarboxylase | 1.28E+08 | 1.53E+05 | 0.00E+00 | 8.79E+05 | -0.25 |

| Protein number | Accession | Description | Relative protein abundance per fraction UV-peak 1 | UV-peak 2 | UV-peak 3 | UV-peak 4 | Pearson correlation score between specific activity and relative protein abundance |
|---|---|---|---|---|---|---|---|
| 108 | A0A2C9CJJ9 | NAD-dependent epimerase/dehydratase domain-containing protein | 2.77E+07 | 7.03E+05 | 8.48E+05 | 9.22E+05 | -0.25 |
| 109 | A0A2C9CDL3 | GTPase | 3.77E+07 | 2.54E+05 | 0.00E+00 | 2.96E+05 | -0.25 |
| 110 | Q1Q6P7 | HD-GYP domain-containing protein | 2.98E+07 | 8.89E+04 | 0.00E+00 | 1.65E+05 | -0.25 |
| 111 | A0A2C9CI57 | dITP/XTP pyrophosphatase | 8.22E+06 | 0.00E+00 | 0.00E+00 | 2.73E+04 | -0.25 |
| 112 | Q1Q0Y3 | Glucose-6-phosphate isomerase | 3.30E+06 | 0.00E+00 | 0.00E+00 | 1.09E+04 | -0.25 |
| 113 | Q1Q117 | strongly similar to SAICAR synthase | 1.37E+08 | 7.95E+04 | 0.00E+00 | 4.20E+05 | -0.25 |
| 114 | A0A2C9CAC1 | strongly similar to glutamate-1-semialdehyde aminomutase | 5.55E+08 | 1.73E+07 | 4.03E+06 | 1.08E+07 | -0.25 |
| 115 | Q1PUT0 | strongly similar to glucose-1-phosphate adenylyltransferase | 1.66E+07 | 0.00E+00 | 0.00E+00 | 0.00E+00 | -0.25 |
| 116 | Q1PV39 | Secondary thiamine-phosphate synthase enzyme | 1.12E+07 | 0.00E+00 | 0.00E+00 | 0.00E+00 | -0.25 |
| 117 | Q1Q0L1 | Tetrapyrrole methylase domain-containing protein | 9.33E+06 | 0.00E+00 | 0.00E+00 | 0.00E+00 | -0.25 |
| 118 | Q1PZV4 | Anthranilate synthase component 1 | 5.95E+06 | 0.00E+00 | 0.00E+00 | 0.00E+00 | -0.25 |
| 119 | A0A2C9CGY3 | Anhydro-N-acetylmuramic acid kinase | 4.83E+06 | 0.00E+00 | 0.00E+00 | 0.00E+00 | -0.25 |
| 120 | Q1Q253 | UDP-N-acetylenolpyruvoylglucosamine reductase | 2.99E+06 | 0.00E+00 | 0.00E+00 | 0.00E+00 | -0.25 |
| 121 | A0A6G7GJQ2 | ATPase AAA-type core domain-containing protein | 2.77E+06 | 0.00E+00 | 0.00E+00 | 0.00E+00 | -0.25 |
| 122 | A0A6G7GX23 | Aspartate aminotransferase family protein | 2.19E+06 | 0.00E+00 | 0.00E+00 | 0.00E+00 | -0.25 |
| 123 | A0A2C9CK09 | hypothetical protein | 2.16E+06 | 0.00E+00 | 0.00E+00 | 0.00E+00 | -0.25 |
| 124 | Q1PZ73 | dTDP-4-dehydrorhamnose 3,5-epimerase | 2.16E+06 | 0.00E+00 | 0.00E+00 | 0.00E+00 | -0.25 |
| 125 | A0A6G7GU28 | LamG-like jellyroll fold domain-containing protein | 1.90E+06 | 0.00E+00 | 0.00E+00 | 0.00E+00 | -0.25 |
| 126 | Q1PVD3 | Probable 6-oxopurine nucleoside phosphorylase | 1.26E+06 | 0.00E+00 | 0.00E+00 | 0.00E+00 | -0.25 |
| 127 | A0A2C9CCA4 | hypothetical protein | 1.16E+06 | 0.00E+00 | 0.00E+00 | 0.00E+00 | -0.25 |
| 128 | Q1Q0Z5 | hypothetical protein | 1.08E+06 | 0.00E+00 | 0.00E+00 | 0.00E+00 | -0.25 |
| 129 | Q1PZ46 | similar to RND multidrug efflux membrane fusion protein MexC precursor | 8.41E+05 | 0.00E+00 | 0.00E+00 | 0.00E+00 | -0.25 |
| 130 | A0A2C9CHD9 | strongly similar to dihydrodipicolinate reductase | 8.26E+05 | 0.00E+00 | 0.00E+00 | 0.00E+00 | -0.25 |
| 131 | Q1PXP9 | Protein-glutamate methylesterase/protein-glutamine glutaminase | 7.90E+05 | 0.00E+00 | 0.00E+00 | 0.00E+00 | -0.25 |
| 132 | A0A2C9CCG0 | hypothetical protein | 6.76E+05 | 0.00E+00 | 0.00E+00 | 0.00E+00 | -0.25 |
| 133 | A0A2C9CGM9 | hypothetical protein | 5.53E+05 | 0.00E+00 | 0.00E+00 | 0.00E+00 | -0.25 |
| 134 | Q1PYC5 | Hydrazine dehydrogenase | 4.84E+05 | 0.00E+00 | 0.00E+00 | 0.00E+00 | -0.25 |
| 135 | Q1Q7Q7 | hypothetical protein | 4.81E+05 | 0.00E+00 | 0.00E+00 | 0.00E+00 | -0.25 |
| 136 | Q1Q0A1 | similar to serine-proteinase HtrA/ DegQ/ DegS family protein | 4.27E+05 | 0.00E+00 | 0.00E+00 | 0.00E+00 | -0.25 |
| 137 | A0A2C9CJS3 | hypothetical protein | 3.91E+05 | 0.00E+00 | 0.00E+00 | 0.00E+00 | -0.25 |
| 138 | Q1Q4R8 | strongly similar to chorismate mutase / prephenate dehydratase | 3.91E+05 | 0.00E+00 | 0.00E+00 | 0.00E+00 | -0.25 |
| 139 | A0A6G7GY31 | hypothetical protein | 3.89E+05 | 0.00E+00 | 0.00E+00 | 0.00E+00 | -0.25 |
| 140 | Q1PXY8 | hypothetical protein | 3.04E+05 | 0.00E+00 | 0.00E+00 | 0.00E+00 | -0.25 |
| 141 | A0A2C9CBC8 | Formiminotetrahydrofolate cyclodeaminase | 1.90E+05 | 0.00E+00 | 0.00E+00 | 0.00E+00 | -0.25 |
| 142 | A0A2C9CBD6 | Putative enzyme | 1.49E+05 | 0.00E+00 | 0.00E+00 | 0.00E+00 | -0.25 |
| 143 | A0A2C9CGE2 | Strongly similar to NAD(P)H:quinone oxidoreductase chain 5 | 1.04E+05 | 0.00E+00 | 0.00E+00 | 0.00E+00 | -0.25 |
| 144 | A0A2C9CCG2 | hypothetical protein | 1.03E+05 | 0.00E+00 | 0.00E+00 | 0.00E+00 | -0.25 |
| 145 | A0A2C9CC83 | strongly similar to 3naS ribosomal protein S2 | 9.27E+04 | 0.00E+00 | 0.00E+00 | 0.00E+00 | -0.25 |
| 146 | A0A2C9CKI0 | Adenylyl-sulfate kinase | 5.15E+04 | 0.00E+00 | 0.00E+00 | 0.00E+00 | -0.25 |
| 147 | Q1Q1N0 | Undecaprenyl phosphate-alpha-4-amino-4-deoxy-L-arabinose transferase | 3.78E+04 | 0.00E+00 | 0.00E+00 | 0.00E+00 | -0.25 |
| 148 | A0A2C9CFR0 | glutamate formimidoyltransferase | 1.69E+07 | 0.00E+00 | 0.00E+00 | 0.00E+00 | -0.25 |
| 149 | A0A2C9CIE4 | Transposase IS204/IS1001/IS1096/IS1165 DDE domain-containing protein | 5.77E+06 | 0.00E+00 | 0.00E+00 | 0.00E+00 | -0.25 |
| 150 | A0A2C9CLN1 | Amidohydrolase-related domain-containing protein | 5.44E+06 | 0.00E+00 | 0.00E+00 | 0.00E+00 | -0.25 |
| 151 | A0A2C9CEK8 | similar to D-3-phosphoglycerate dehydrogenase (PGDH) | 3.32E+06 | 0.00E+00 | 0.00E+00 | 0.00E+00 | -0.25 |
| 152 | Q1Q3C3 | hypothetical protein | 2.99E+06 | 0.00E+00 | 0.00E+00 | 0.00E+00 | -0.25 |
| 153 | Q1PXZ8 | Polar-differentiation response regulator DivK | 2.95E+06 | 0.00E+00 | 0.00E+00 | 0.00E+00 | -0.25 |
| 154 | A0A2C9CE43 | hypothetical protein | 2.52E+06 | 0.00E+00 | 0.00E+00 | 0.00E+00 | -0.25 |
| 155 | A0A2C9CH82 | similar to flavoproteins norVW and fprA | 2.29E+06 | 0.00E+00 | 0.00E+00 | 0.00E+00 | -0.25 |
| 156 | A0A2C9CIJ9 | GGDEF domain-containing protein | 2.25E+06 | 0.00E+00 | 0.00E+00 | 0.00E+00 | -0.25 |
| 157 | Q1PZ80 | strongly similar to phosphoglycerate kinase | 2.12E+06 | 0.00E+00 | 0.00E+00 | 0.00E+00 | -0.25 |
| 158 | A0A2C9CDM5 | Glutamine--fructose-6-phosphate aminotransferase [isomerizing] | 1.93E+06 | 0.00E+00 | 0.00E+00 | 0.00E+00 | -0.25 |
| 159 | A0A2C9CIF4 | Transposase IS200 like protein | 1.77E+06 | 0.00E+00 | 0.00E+00 | 0.00E+00 | -0.25 |
| 160 | A0A2C9CKJ5 | Gluconeogenesis factor | 1.66E+06 | 0.00E+00 | 0.00E+00 | 0.00E+00 | -0.25 |
| 161 | Q1PXW3 | strongly similar to ATP-dependent protease La | 1.58E+06 | 0.00E+00 | 0.00E+00 | 0.00E+00 | -0.25 |
| 162 | A0A2C9CHA0 | strongly similar to 5naS ribosomal protein L7/L12 | 1.29E+06 | 0.00E+00 | 0.00E+00 | 0.00E+00 | -0.25 |

| Protein number | Accession | Description | Relative protein abundance per fraction | | | | Pearson correlation score |
|---|---|---|---|---|---|---|---|
| | | | UV-peak 1 | UV-peak 2 | UV-peak 3 | UV-peak 4 | between specific activity and relative protein abundance |
| 163 | A0A2C9CKL5 | Regulator of chromosome condensation (RCC1) repeat protein | 1.27E+06 | 0.00E+00 | 0.00E+00 | 0.00E+00 | -0.25 |
| 164 | A0A2C9CDL6 | hypothetical protein | 1.23E+06 | 0.00E+00 | 0.00E+00 | 0.00E+00 | -0.25 |
| 165 | A0A2C9CK66 | Multifunctional fusion protein | 1.19E+06 | 0.00E+00 | 0.00E+00 | 0.00E+00 | -0.25 |
| 166 | Q1Q4S6 | hypothetical protein | 9.71E+05 | 0.00E+00 | 0.00E+00 | 0.00E+00 | -0.25 |
| 167 | Q1PZE3 | Carboxypeptidase regulatory-like domain-containing protein | 9.55E+05 | 0.00E+00 | 0.00E+00 | 0.00E+00 | -0.25 |
| 168 | A0A2C9CCR3 | hypothetical protein | 9.42E+05 | 0.00E+00 | 0.00E+00 | 0.00E+00 | -0.25 |
| 169 | Q1PUH0 | hypothetical protein | 9.15E+05 | 0.00E+00 | 0.00E+00 | 0.00E+00 | -0.25 |
| 170 | Q1Q6W7 | Putative mannose-1-phosphate guanylyltransferase | 8.96E+05 | 0.00E+00 | 0.00E+00 | 0.00E+00 | -0.25 |
| 171 | Q1PX51 | DUF3368 domain-containing protein | 8.77E+05 | 0.00E+00 | 0.00E+00 | 0.00E+00 | -0.25 |
| 172 | Q1Q3Y4 | Purine nucleoside phosphorylase | 8.74E+05 | 0.00E+00 | 0.00E+00 | 0.00E+00 | -0.25 |
| 173 | A0A2C9CK74 | Cobalamin biosynthesis precorrin-8X methylmutase CobH protein | 8.62E+05 | 0.00E+00 | 0.00E+00 | 0.00E+00 | -0.25 |
| 174 | A0A2C9CHW3 | Chorismate synthase | 8.60E+05 | 0.00E+00 | 0.00E+00 | 0.00E+00 | -0.25 |
| 175 | Q1Q325 | 3-oxoacyl-[acyl-carrier-protein] reductase | 8.25E+05 | 0.00E+00 | 0.00E+00 | 0.00E+00 | -0.25 |
| 176 | Q1Q3U9 | hypothetical protein | 7.91E+05 | 0.00E+00 | 0.00E+00 | 0.00E+00 | -0.25 |
| 177 | Q1PV08 | hypothetical protein | 7.55E+05 | 0.00E+00 | 0.00E+00 | 0.00E+00 | -0.25 |
| 178 | Q1PZK1 | hypothetical protein | 7.45E+05 | 0.00E+00 | 0.00E+00 | 0.00E+00 | -0.25 |
| 179 | A0A2C9CBT8 | ABC transporter domain-containing protein | 7.41E+05 | 0.00E+00 | 0.00E+00 | 0.00E+00 | -0.25 |
| 180 | A0A2C9CBS9 | strongly similar to aspartate-semialdehyde dehydrogenase Asd | 7.29E+05 | 0.00E+00 | 0.00E+00 | 0.00E+00 | -0.25 |
| 181 | A0A2C9CJI8 | similar to aminopeptidase A | 6.79E+05 | 0.00E+00 | 0.00E+00 | 0.00E+00 | -0.25 |
| 182 | A0A2C9CCE6 | HD-GYP domain-containing protein | 6.08E+05 | 0.00E+00 | 0.00E+00 | 0.00E+00 | -0.25 |
| 183 | Q1Q6N2 | Transposase IS4-like domain-containing protein | 6.00E+05 | 0.00E+00 | 0.00E+00 | 0.00E+00 | -0.25 |
| 184 | Q1Q1A6 | hypothetical protein | 5.47E+05 | 0.00E+00 | 0.00E+00 | 0.00E+00 | -0.25 |
| 185 | Q1Q156 | strongly similar to adenylate kinase | 4.45E+05 | 0.00E+00 | 0.00E+00 | 0.00E+00 | -0.25 |
| 186 | Q1PVS7 | Metallo-beta-lactamase family protein, RNA-specific | 4.38E+05 | 0.00E+00 | 0.00E+00 | 0.00E+00 | -0.25 |
| 187 | Q1Q3W8 | hypothetical protein | 3.70E+05 | 0.00E+00 | 0.00E+00 | 0.00E+00 | -0.25 |
| 188 | Q1PZ79 | Ribulose-phosphate 3-epimerase | 3.63E+05 | 0.00E+00 | 0.00E+00 | 0.00E+00 | -0.25 |
| 189 | Q1PZ47 | DNA-(apurinic or apyrimidinic site) lyase | 3.00E+05 | 0.00E+00 | 0.00E+00 | 0.00E+00 | -0.25 |
| 190 | Q1PX52 | Ribbon-helix-helix protein CopG domain-containing protein | 2.96E+05 | 0.00E+00 | 0.00E+00 | 0.00E+00 | -0.25 |
| 191 | Q1PY25 | strongly similar to thioredoxin | 2.67E+05 | 0.00E+00 | 0.00E+00 | 0.00E+00 | -0.25 |
| 192 | Q1Q247 | Small ribosomal subunit protein bS16 | 2.57E+05 | 0.00E+00 | 0.00E+00 | 0.00E+00 | -0.25 |
| 193 | Q1Q1K7 | Ferric uptake regulation protein FUR | 2.48E+05 | 0.00E+00 | 0.00E+00 | 0.00E+00 | -0.25 |
| 194 | Q1PUZ1 | strongly similar to 6-phosphogluconate dehydrogenase (decarboxylating) | 2.36E+05 | 0.00E+00 | 0.00E+00 | 0.00E+00 | -0.25 |
| 195 | Q1Q5Y4 | Reverse transcriptase domain-containing protein | 2.27E+05 | 0.00E+00 | 0.00E+00 | 0.00E+00 | -0.25 |
| 196 | Q1PY42 | strongly similar to 1na kDa chaperonin (GroES protein) | 2.09E+05 | 0.00E+00 | 0.00E+00 | 0.00E+00 | -0.25 |
| 197 | Q1PUZ9 | hypothetical protein | 2.08E+05 | 0.00E+00 | 0.00E+00 | 0.00E+00 | -0.25 |
| 198 | A0A2C9CFP5 | Transposase (putative) YhgA-like domain-containing protein | 1.69E+05 | 0.00E+00 | 0.00E+00 | 0.00E+00 | -0.25 |
| 199 | A0A2C9CFA5 | strongly similar to pyrroline-5-carboxylate reductase | 1.56E+05 | 0.00E+00 | 0.00E+00 | 0.00E+00 | -0.25 |
| 200 | Q1Q5M6 | hypothetical protein | 1.47E+05 | 0.00E+00 | 0.00E+00 | 0.00E+00 | -0.25 |
| 201 | Q1Q4Q2 | strongly similar to Dna binding protein HU | 1.17E+05 | 0.00E+00 | 0.00E+00 | 0.00E+00 | -0.25 |
| 202 | A0A2C9CH33 | Strongly similar to molybdenum cofactor biosynthesis protein C | 1.16E+05 | 0.00E+00 | 0.00E+00 | 0.00E+00 | -0.25 |
| 203 | Q1Q205 | hypothetical protein | 1.05E+05 | 0.00E+00 | 0.00E+00 | 0.00E+00 | -0.25 |
| 204 | A0A2C9CH91 | hypothetical protein | 1.04E+05 | 0.00E+00 | 0.00E+00 | 0.00E+00 | -0.25 |
| 205 | Q1PX08 | strongly similar to nitrogen regulatory protein P-II | 9.15E+04 | 0.00E+00 | 0.00E+00 | 0.00E+00 | -0.25 |
| 206 | Q1PXM3 | Response regulatory domain-containing protein | 8.53E+04 | 0.00E+00 | 0.00E+00 | 0.00E+00 | -0.25 |
| 207 | Q1Q6A7 | strongly similar to cold shock protein A | 7.96E+04 | 0.00E+00 | 0.00E+00 | 0.00E+00 | -0.25 |
| 208 | Q1PYG6 | hypothetical protein | 6.33E+04 | 0.00E+00 | 0.00E+00 | 0.00E+00 | -0.25 |
| 209 | Q1PX26 | hypothetical protein | 5.40E+04 | 0.00E+00 | 0.00E+00 | 0.00E+00 | -0.25 |
| 210 | Q1PYI8 | strongly similar to alanyl-tRna synthetase | 3.90E+04 | 0.00E+00 | 0.00E+00 | 0.00E+00 | -0.25 |
| 211 | A0A6G7GJL0 | DUF3368 domain-containing protein | 3.42E+04 | 0.00E+00 | 0.00E+00 | 0.00E+00 | -0.25 |
| 212 | A0A6G7GSL2 | Putative divergent AAA domain family | 2.86E+04 | 0.00E+00 | 0.00E+00 | 0.00E+00 | -0.25 |
| 213 | A0A2C9CKH9 | Hypoxanthine phosphoribosyltransferase | 2.09E+04 | 0.00E+00 | 0.00E+00 | 0.00E+00 | -0.25 |
| 214 | A0A2C9CL87 | similar to UDP-glucose 4-epimerase | 6.30E+06 | 0.00E+00 | 0.00E+00 | 0.00E+00 | -0.25 |
| 215 | A0A2C9CME9 | Nitroreductase domain-containing protein | 1.13E+06 | 0.00E+00 | 0.00E+00 | 0.00E+00 | -0.25 |
| 216 | Q1Q1J8 | Catecholate siderophore receptor CirA | 4.34E+04 | 0.00E+00 | 0.00E+00 | 0.00E+00 | -0.25 |
| 217 | Q1PYG4 | GDP-L-fucose synthase | 1.44E+04 | 0.00E+00 | 0.00E+00 | 0.00E+00 | -0.25 |

| Protein number | Accession | Description | Relative protein abundance per fraction | | | | Pearson correlation score |
|---|---|---|---|---|---|---|---|
| | | | UV-peak 1 | UV-peak 2 | UV-peak 3 | UV-peak 4 | between specific activity and relative protein abundance |
| 218 | Q1PZT0 | Anthranilate phosphoribosyltransferase | 9.59E+06 | 2.28E+04 | 0.00E+00 | 0.00E+00 | -0.25 |
| 219 | A0A2C9CIN1 | hypothetical protein | 6.63E+06 | 4.66E+04 | 0.00E+00 | 1.40E+04 | -0.25 |
| 220 | Q1Q2R9 | Putative sorbitol dehydrogenase | 3.42E+08 | 5.58E+06 | 4.32E+06 | 4.34E+06 | -0.25 |
| 221 | Q1PZ82 | strongly similar to glyceraldehyde-3-phosphate dehydrogenase | 1.68E+08 | 2.53E+07 | 0.00E+00 | 1.11E+07 | -0.25 |
| 222 | Q1PZC7 | 4-diphosphocytidyl-2-C-methyl-D-erythritol kinase | 4.04E+07 | 2.93E+05 | 0.00E+00 | 4.10E+04 | -0.25 |
| 223 | Q1PYA3 | Putative beta-lactamase-inhibitor-like PepSY-like domain-containing protein | 1.27E+07 | 3.60E+05 | 0.00E+00 | 9.86E+04 | -0.26 |
| 224 | Q1Q013 | strongly similar to phosphomannomutase | 1.01E+08 | 1.08E+07 | 0.00E+00 | 3.88E+06 | -0.26 |
| 225 | Q1Q1I6 | Benzoyl-CoA reductase subunit BadG | 7.81E+06 | 0.00E+00 | 3.66E+06 | 1.67E+06 | -0.26 |
| 226 | Q1Q5S6 | 3-phosphoshikimate 1-carboxyvinyltransferase | 4.57E+06 | 1.28E+05 | 0.00E+00 | 0.00E+00 | -0.26 |
| 227 | Q1Q6L2 | Glycogen synthase | 9.63E+06 | 4.21E+05 | 0.00E+00 | 6.42E+04 | -0.26 |
| 228 | A0A2C9CMF8 | hypothetical protein | 7.62E+05 | 2.58E+04 | 0.00E+00 | 0.00E+00 | -0.27 |
| 229 | Q1PY54 | Methyltransferase domain-containing protein | 1.13E+06 | 4.52E+04 | 0.00E+00 | 0.00E+00 | -0.27 |
| 230 | Q1Q468 | Exported protein | 4.83E+07 | 2.41E+06 | 0.00E+00 | 1.38E+05 | -0.27 |
| 231 | A0A2C9CD72 | hypothetical protein | 2.12E+06 | 1.40E+05 | 0.00E+00 | 0.00E+00 | -0.28 |
| 232 | A0A2C9CKP3 | Cobalt-precorrin-3b C17-methyltransferase | 2.31E+06 | 1.60E+05 | 0.00E+00 | 0.00E+00 | -0.28 |
| 233 | Q1PV12 | Aminotransferase | 8.43E+07 | 9.90E+06 | 0.00E+00 | 8.63E+05 | -0.29 |
| 234 | Q1Q637 | hypothetical protein | 1.23E+08 | 1.50E+07 | 2.23E+06 | 2.72E+06 | -0.29 |
| 235 | A0A2C9CE24 | strongly similar to serine hydroxymethyl transferase SHMT | 3.12E+08 | 6.05E+07 | 1.26E+07 | 1.89E+07 | -0.30 |
| 236 | Q1Q724 | similar to serine protease Do | 1.93E+06 | 2.23E+05 | 0.00E+00 | 0.00E+00 | -0.30 |
| 237 | A0A2C9CGA0 | Small heat shock protein-like protein | 1.08E+06 | 2.56E+05 | 0.00E+00 | 5.78E+04 | -0.30 |
| 238 | Q1PZJ0 | similar to flagellar motor protein MotB | 2.24E+06 | 2.64E+05 | 0.00E+00 | 0.00E+00 | -0.30 |
| 239 | Q1Q4B8 | Nucleotidyl transferase AbiEii/AbiGii toxin family protein | 9.45E+06 | 8.81E+05 | 2.53E+05 | 2.33E+04 | -0.30 |
| 240 | Q1Q7N4 | similar to (3R)-hydroxymyristoyl acyl carrier protein dehydrase | 0.00E+00 | 6.34E+05 | 0.00E+00 | 1.90E+04 | -0.32 |
| 241 | A0A2C9CFE9 | Fragment of pimeloyl-CoA synthetase (Part 1) | 4.65E+06 | 7.42E+05 | 0.00E+00 | 0.00E+00 | -0.32 |
| 242 | A0A2C9CCN1 | Fragment of pimeloyl-CoA synthetase (Part 2) | 4.75E+06 | 7.59E+05 | 0.00E+00 | 0.00E+00 | -0.32 |
| 243 | Q1PXW4 | N-acetyl-gamma-glutamyl-phosphate reductase | 1.29E+07 | 3.87E+06 | 0.00E+00 | 7.60E+05 | -0.32 |
| 244 | A0A2C9CBM9 | Fibronectin type-III domain-containing protein | 4.50E+06 | 7.51E+05 | 0.00E+00 | 0.00E+00 | -0.32 |
| 245 | Q1PY88 | similar to superoxide dismutase | 5.88E+07 | 1.33E+06 | 9.88E+06 | 1.25E+06 | -0.33 |
| 246 | Q1PZC8 | similar to unknown protein involved in septum location | 2.01E+06 | 5.07E+05 | 0.00E+00 | 6.89E+04 | -0.33 |
| 247 | A0A2C9CFE7 | Thymidylate kinase | 2.21E+06 | 6.71E+05 | 0.00E+00 | 1.23E+05 | -0.33 |
| 248 | A0A2C9CJJ8 | Exoribonuclease YhaM-like protein | 1.33E+06 | 2.35E+05 | 0.00E+00 | 0.00E+00 | -0.33 |
| 249 | A0A2C9CJE3 | hypothetical protein | 1.22E+07 | 2.60E+06 | 0.00E+00 | 1.56E+05 | -0.33 |
| 250 | A0A2C9CHS3 | similar to 3-isopropylmalate dehydratase, large subunit | 1.88E+06 | 3.54E+05 | 0.00E+00 | 0.00E+00 | -0.33 |
| 251 | Q1Q1N5 | Acetate-CoA ligase [ADP-forming] I | 1.09E+08 | 2.19E+07 | 3.56E+05 | 4.66E+05 | -0.33 |
| 252 | Q1Q4S4 | L,D-transpeptidase YkuD | 1.89E+07 | 4.05E+06 | 0.00E+00 | 1.18E+05 | -0.34 |
| 253 | A0A2C9CLA0 | Methionyl-tRNA formyltransferase | 2.95E+06 | 9.82E+06 | 1.67E+05 | 1.24E+06 | -0.34 |
| 254 | A0A2C9CIL6 | Phospholipase D-like domain-containing protein | 0.00E+00 | 5.93E+06 | 0.00E+00 | 0.00E+00 | -0.34 |
| 255 | A0A6G7GQY5 | hypothetical protein | 0.00E+00 | 3.83E+05 | 0.00E+00 | 0.00E+00 | -0.34 |
| 256 | A0A2C9CHK8 | DUF11 domain-containing protein | 0.00E+00 | 6.75E+06 | 0.00E+00 | 0.00E+00 | -0.34 |
| 257 | Q1PXN9 | Putative purine | 0.00E+00 | 1.88E+06 | 0.00E+00 | 0.00E+00 | -0.34 |
| 258 | Q1PZ64 | Competence protein ComM | 0.00E+00 | 1.45E+06 | 0.00E+00 | 0.00E+00 | -0.34 |
| 259 | Q1PV42 | similar to naD(P) oxidoreductase, FAD-containing subunit | 0.00E+00 | 8.21E+05 | 0.00E+00 | 0.00E+00 | -0.34 |
| 260 | Q1Q5P4 | Putative desampylase | 0.00E+00 | 6.45E+05 | 0.00E+00 | 0.00E+00 | -0.34 |
| 261 | A0A2C9CG62 | DUF262 domain-containing protein | 0.00E+00 | 3.04E+05 | 0.00E+00 | 0.00E+00 | -0.34 |
| 262 | A0A2C9CF87 | TonB-dependent receptor | 0.00E+00 | 3.03E+05 | 0.00E+00 | 0.00E+00 | -0.34 |
| 263 | Q1PXU8 | hypothetical protein | 0.00E+00 | 1.49E+05 | 0.00E+00 | 0.00E+00 | -0.34 |
| 264 | Q1PVL5 | Helicase C-terminal domain-containing protein | 0.00E+00 | 2.85E+06 | 0.00E+00 | 0.00E+00 | -0.34 |
| 265 | A0A2C9CFN1 | Aminoglycoside phosphotransferase domain-containing protein | 0.00E+00 | 1.05E+06 | 0.00E+00 | 0.00E+00 | -0.34 |
| 266 | A0A2C9CCF9 | hypothetical protein | 0.00E+00 | 2.13E+05 | 0.00E+00 | 0.00E+00 | -0.34 |
| 267 | A0A2C9CI35 | hypothetical protein | 8.43E+06 | 4.53E+06 | 0.00E+00 | 1.03E+06 | -0.35 |
| 268 | A0A2C9CAV3 | hypothetical protein | 1.32E+06 | 2.82E+07 | 3.77E+05 | 2.75E+05 | -0.36 |
| 269 | Q1Q5R5 | Aspartokinase | 1.64E+06 | 4.63E+05 | 0.00E+00 | 2.53E+03 | -0.37 |
| 270 | Q1Q0T9 | hypothetical (triheme) protein | 1.60E+06 | 1.29E+07 | 5.08E+09 | 1.65E+08 | -0.37 |
| 271 | Q1PUI8 | Thiamine-phosphate synthase | 1.89E+05 | 5.32E+05 | 0.00E+00 | 4.61E+04 | -0.38 |
| 272 | Q1PY86 | hypothetical protein | 4.21E+06 | 3.76E+06 | 0.00E+00 | 7.09E+05 | -0.39 |

| Protein number | Accession | Description | Relative protein abundance per fraction UV-peak 1 | UV-peak 2 | UV-peak 3 | UV-peak 4 | Pearson correlation score between specific activity and relative protein abundance |
|---|---|---|---|---|---|---|---|
| 273 | Q1Q277 | hypothetical protein | 2.31E+06 | 8.37E+05 | 0.00E+00 | 2.67E+04 | -0.39 |
| 274 | Q1Q1K9 | hypothetical protein | 6.95E+06 | 1.40E+07 | 1.85E+08 | 1.31E+07 | -0.39 |
| 275 | A0A2C9CHA8 | similar to intracellular proteinase I | 2.35E+07 | 9.38E+07 | 1.32E+06 | 4.19E+06 | -0.40 |
| 276 | A0A2C9CAR7 | Murein hydrolase activator NlpD | 1.30E+06 | 4.73E+05 | 0.00E+00 | 0.00E+00 | -0.40 |
| 277 | Q1PZB4 | hypothetical protein | 7.79E+05 | 2.85E+05 | 0.00E+00 | 0.00E+00 | -0.40 |
| 278 | A0A2C9CD11 | Acetoacetate metabolism regulatory protein AtoC | 1.66E+06 | 1.45E+06 | 0.00E+00 | 2.40E+05 | -0.40 |
| 279 | A0A2C9CHE2 | strongly similar to glutamine amidotransferase class I | 4.93E+05 | 1.99E+05 | 0.00E+00 | 0.00E+00 | -0.41 |
| 280 | A0A2C9CJG4 | Dihydrolipoamide acetyltransferase component of pyruvate dehydrogenase | 4.19E+04 | 4.14E+05 | 1.19E+07 | 1.20E+04 | -0.42 |
| 281 | A0A2C9CJ81 | strongly similar to S-adenosylmethionine synthetase | 8.02E+06 | 3.70E+08 | 1.84E+08 | 6.82E+07 | -0.42 |
| 282 | Q1PW67 | similar to peptidyl-prolyl cis-trans isomerase | 2.23E+04 | 7.18E+04 | 0.00E+00 | 0.00E+00 | -0.45 |
| 283 | Q1Q1P5 | Ketoacid-binding protein | 1.17E+06 | 6.66E+05 | 0.00E+00 | 0.00E+00 | -0.46 |
| 284 | Q1PY19 | tRNA-splicing ligase RtcB | 1.11E+07 | 9.04E+06 | 0.00E+00 | 6.57E+05 | -0.46 |
| 285 | A0A2C9CC13 | similar to ribosome recycling factor | 5.24E+07 | 9.21E+07 | 4.93E+04 | 3.78E+06 | -0.47 |
| 286 | Q1Q2M7 | Putative NADP-dependent glyceraldehyde-3-phosphate dehydrogenase GapN | 5.49E+06 | 8.35E+06 | 0.00E+00 | 4.30E+05 | -0.47 |
| 287 | Q1Q1G5 | strongly similar to translation initiation factor IF-2 | 1.16E+06 | 4.65E+06 | 3.73E+05 | 0.00E+00 | -0.48 |
| 288 | A0A2C9CLS3 | hydroxymethylpyrimidine kinase | 7.45E+06 | 5.45E+06 | 0.00E+00 | 9.55E+04 | -0.48 |
| 289 | Q1PUW1 | CBS domain-containing protein | 7.80E+05 | 8.18E+05 | 0.00E+00 | 0.00E+00 | -0.52 |
| 290 | A0A6G7GXG7 | Monoheme cytochrome | 3.29E+05 | 4.24E+05 | 0.00E+00 | 0.00E+00 | -0.52 |
| 291 | Q1Q096 | Uroporphyrin-III C-methyltransferase | 7.04E+04 | 8.66E+04 | 0.00E+00 | 0.00E+00 | -0.52 |
| 292 | A0A2C9CIK3 | Outer membrane protein | 1.17E+07 | 1.77E+08 | 1.32E+08 | 3.09E+07 | -0.54 |
| 293 | A0A2C9CGT8 | c-type diheme-containing protein, redox partner of kustc0458 | 2.47E+08 | 9.09E+08 | 2.58E+08 | 5.19E+07 | -0.57 |
| 294 | A0A2C9CAZ0 | hypothetical protein | 1.68E+07 | 8.17E+06 | 3.66E+06 | 2.63E+05 | -0.58 |
| 295 | A0A2C9CH11 | Ketol-acid reductoisomerase (NADP(+)) | 2.61E+07 | 3.75E+08 | 2.97E+08 | 2.01E+07 | -0.64 |
| 296 | A0A2C9CJP1 | similar to hydroxylamine oxidoreductase hao | 0.00E+00 | 3.00E+06 | 3.14E+06 | 0.00E+00 | -0.65 |
| 297 | Q1Q2C3 | Phospholipase D precursor | 3.72E+06 | 2.18E+06 | 2.73E+06 | 2.93E+04 | -0.88 |
| 298 | Q1PZ43 | Putative pterin-4-alpha-carbinolamine dehydratase | 8.91E+05 | 1.09E+06 | 1.07E+06 | 2.87E+05 | -0.99 |

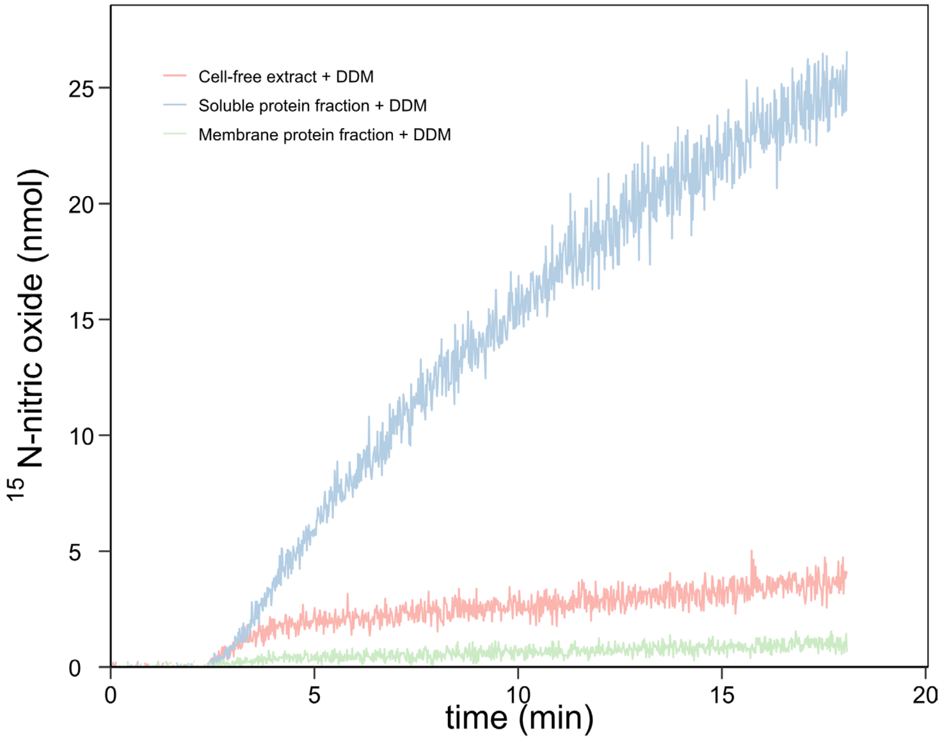

**Supplementary figure 1** – $^{15}$N-labeled nitric oxide production from $^{15}$N-labeled nitrite by cell-free anammox extract, soluble protein fraction, and membrane protein fraction incubated with 1% n-dodecyl β-D-maltoside (DDM). Nitric oxide production in membrane proteins and cell-free extract stopped around four minutes whereas the production in soluble proteins continued. Activity assays contained 200 µM ascorbate and phenazine ethosulfate, and 6-10 µg protein in 20 mM MOPS, 150 mM NaCl buffer, pH 7.5. The reaction was started with 77 µM $^{15}$N-nitrite and carried out at 30°C. (*n*= 1)

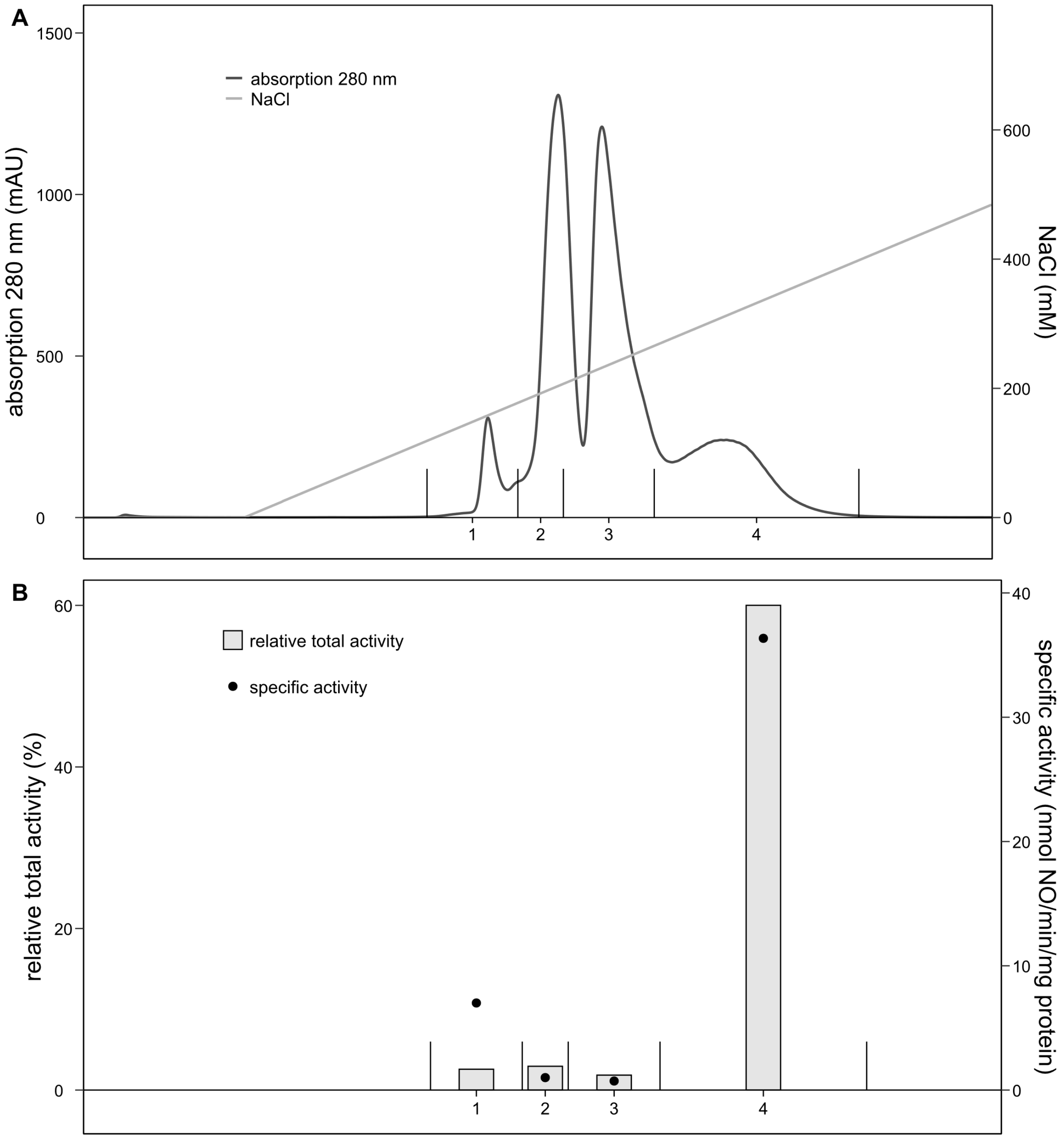

**Supplementary figure 2** – Separation of the proteins in sample B by high-resolution anion-exchange column chromatography and their nitrite reductase activity. (A) Soluble proteins were initially separated on a low-resolution column of which sample B showed high specific and relative total activity compared to activity measured in all soluble proteins. To enrich the active nitrite reductase in sample B, proteins were further separated on a high-resolution anion-exchanger. Proteins were eluted with a linear gradient from 0 to 1 M NaCl and fractionated based on their UV signal. (B) UV peak 4 was the most enriched and most active fraction after two-step column chromatography separation. Here, nitrite reductase produced 36 nmol nitric oxide/min/mg protein which accounted for 60% of the total nitrite reductase activity measured for all soluble proteins. Activity assays contained 200 µM ascorbate and phenazine ethosulfate, 6-10 µg protein in 20 mM MOPS. 150 mM NaCl buffer, pH 7.5. The reaction was started with 77 µM $^{15}$N-nitrite and carried out at 30°C. The relative total activity compared to activity in the total soluble protein fraction is expressed in percentage. The specific activity is indicated by the black dots. (*n*=1)
